# Emergence and evolution of heterocyte glycolipid biosynthesis enabled specialized nitrogen fixation in cyanobacteria

**DOI:** 10.1101/2024.05.17.594646

**Authors:** Ruth Pérez Gallego, F. A. Bastiaan von Meijenfeldt, Nicole J. Bale, Jaap S. Sinninghe Damsté, Laura Villanueva

## Abstract

Paleontological and phylogenomic observations have shed light on the evolution of cyanobacteria. Nevertheless, the emergence of heterocytes, specialized cells for nitrogen fixation, remains unclear. Heterocytes are surrounded by heterocyte glycolipids (HGs), which contribute to protection of the nitrogenase enzyme from oxygen. Here, by comprehensive HG identification and screening of HG biosynthesis genes throughout cyanobacteria, we identify HG analogs produced by specific and distantly related non-heterocytous cyanobacteria. These structurally less complex molecules probably acted as precursors of HGs, suggesting that HGs arose after a genomic reorganization and expansion of ancestral biosynthetic machinery, enabling the rise of cyanobacterial heterocytes in an increasingly oxygenated atmosphere. Subsequently, HG chemical structure evolved convergently in response to environmental pressures. Our results open a new chapter in the potential use of diagenetic products of HGs and HG analogs as fossils for reconstructing the evolution of multicellularity and division of labor in cyanobacteria.

---

Nitrogen-fixing (diazotrophic) cyanobacteria play a major role in nitrogen cycling by transforming N_2_ to biologically available NH_4_^+^ (ref. ^1–4^). To overcome the inhibition by oxygen of the nitrogenase enzyme responsible for nitrogen fixation, diazotrophic cyanobacteria have evolved strategies that separate oxygen-sensitive nitrogen fixation from oxygen-producing photosynthesis^5^. One strategy involves the confinement of the nitrogen fixation reaction to heterocytes, specialized non-photosynthetic cells. Heterocytes are surrounded by a thick-walled cell envelope, composed of polysaccharides and long-chain lipids with sugar headgroups called heterocyte glycolipids (HGs), that limits oxygen diffusion into the cell^6^.

HGs detected in cultures of heterocytous cyanobacteria and in the environment are structurally diverse (Fig. 1a,b), spanning a variety of alkyl chain lengths (C_26_, C_28_, C_30_, and C_32_), attached to different sugar headgroups (hexoses (C_6_) and pentose (C_5_)), with two or three keto or alcohol groups at fixed positions in their alkyl chain^7–11^. 19 distinct HGs have been structurally identified to date^12^. HGs are biosynthesized by polyketide synthases (PKSs) that extend and reduce a growing acyl chain in successive rounds leading to the formation of a so-called aglycone (AG), which is subsequently attached to a sugar headgroup (Fig. 1a,b, Supplementary Fig. 1; refs. ^6,13–15)^. Thus, enzymatic differences between cyanobacteria may result in the production of structurally different HGs, which may reflect adaptive differences in membrane impermeabilization to N_2_ and oxygen^9,16^. HGs can be preserved in the sedimentary record and have been used as indicators of past nitrogen fixation by cyanobacteria^17^, to differentiate specific families and genera of heterocytous cyanobacteria as taxonomic markers^18^, and to detect cyanobacterial symbionts of marine diatoms^19^. Additionally, specific HGs found in the sedimentary record have been used to reconstruct past surface water temperatures^20^. However, despite the relevance of HGs for nitrogen fixation and their potential as biomarkers, knowledge on their structural diversity is limited because to date, only few cyanobacterial cultures have been examined with high-resolution mass spectrometry techniques allowing for comprehensive HG identification, including those present in low abundance.

**Fig. 1.**
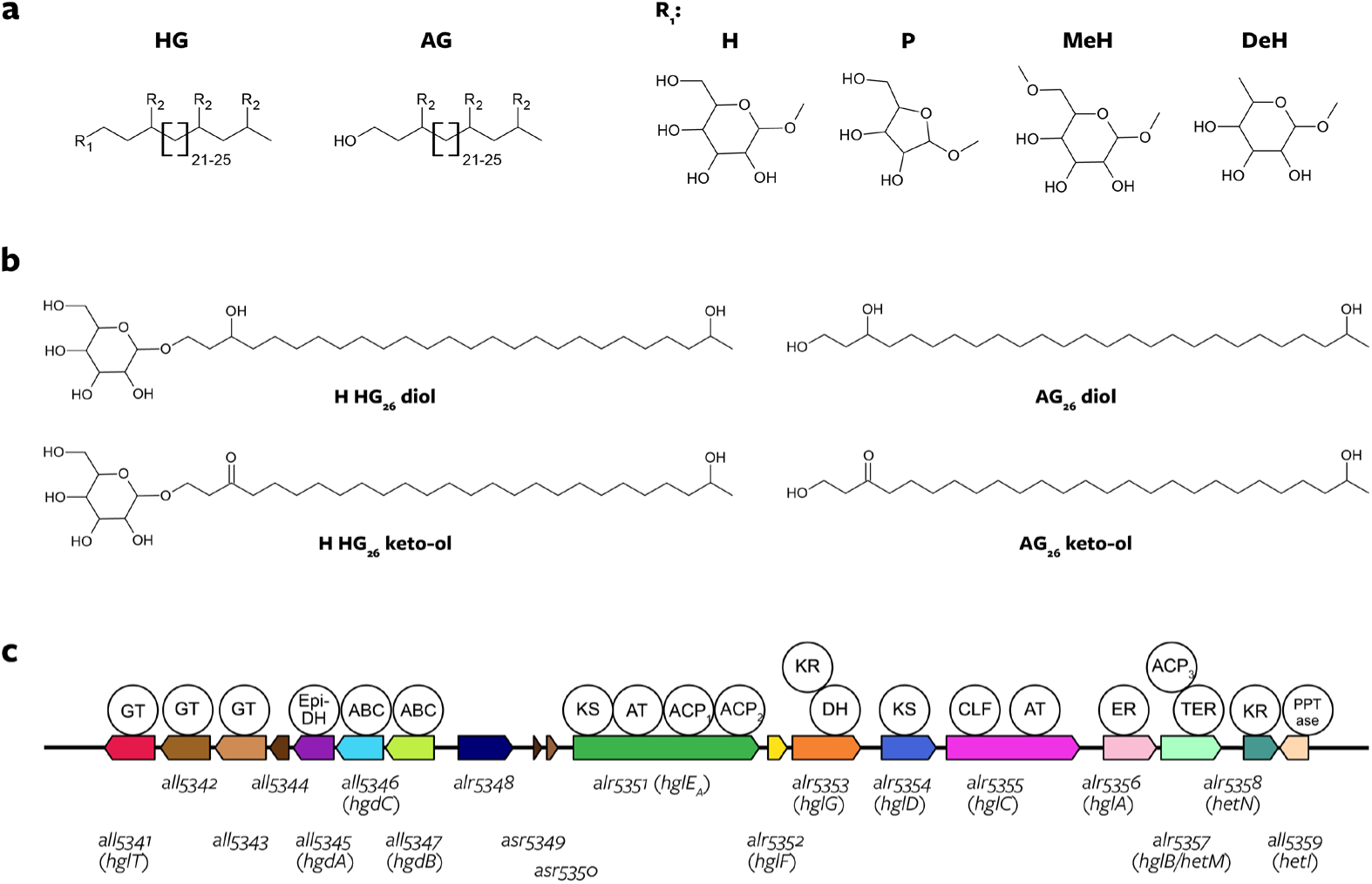
Chemical structure of heterocyte glycolipids (HGs), and schematic representation of the *hgl* island of *Anabaena* sp. PCC 7120 and its most common products. **a**, HGs consist of a sugar headgroup that is attached to an aglycone (AG). Headgroups (R_1_) are H, hexose; P, pentose; MeH, methyl hexose; and DeH, deoxyhexose. Functional groups on the alkyl chain (R_2_) are H, -OH (alcohol), and =O (ketone). 19 distinct HG structures have previously been identified in heterocytous cyanobacterial cultures and in the environment (refs. *^7,9,10,18,39,40^*). **b**, Chemical structure of the most abundant HGs and AGs produced by *Anabaena* sp. PCC 7120. **c**, Gene cluster in *Anabaena* sp. PCC 7120 that contains genes involved in HG biosynthesis and transport (the *hgl* island). Similar clusters had also been identified in several other heterocyte-forming cyanobacteria*^6,13,15^*. Catalytic domains (circles) are plotted on top. Numbered genes depict the genomic naming scheme of *Anabaena* sp. PCC 7120, gene names assigned in previous publications are depicted within brackets. The known functions of *Anabaena* sp. PCC 7120 *hgl* island genes are shown in Supplementary Table 4, and the complete HG biosynthetic pathway is depicted in Supplementary Fig. 1. GT, glycosyl transferase; Epi-DH, epimerase-dehydratase; ABC, ATP-binding cassette (ABC) transporter; KS, ketoacyl synthase; AT, acyl transferase; ACP, acyl carrier protein; KR, ketoreductase; DH, dehydratase; CLF, chain length factor; ER, enoyl reductase; TER, thioester reductase; PPTase, phosphopantetheinyltransferase.

Fossil evidence and phylogenetic analyses suggest that cyanobacteria capable of heterocyte differentiation arose between 2,450 and 2,100 million years ago (Ma), around the same time and possibly simultaneously with multicellularity and nitrogen fixation^21–23^. Evolutionary pressure for an oxygen-impermeable lipid layer to shelter the nitrogenase enzyme was likely driven by the rising oxygen levels of Earth’s atmosphere around 2,400 Ma^21,24^. Heterocytes have been suggested to be evolutionary related to akinetes^25^, spore-like cells produced under stress conditions by some heterocytous cyanobacteria, which are also surrounded by HGs^25,26^. Here, we reconstructed the acquisition and evolution of the biosynthetic capability to produce HGs by screening ∼3,600 cyanobacterial genomes and plasmids for key genes involved in HG formation and their deposition in the cell envelope of the heterocyte. We further analyzed the lipid composition of 26 heterocytous and of 2 non-heterocytous cyanobacterial cultures using high-resolution mass spectrometry to elucidate the connection between biosynthetic capability encoded by the genome and the generated biosynthetic product. Our results reveal that HG structure evolved convergently within heterocytous cyanobacteria and suggest an origin of HGs from structurally similar molecules still produced by some non-heterocytous cyanobacteria and that are not involved in nitrogen fixation nor akinete formation today.

## Results and Discussion

### Genomic prediction of HG biosynthesis in Cyanobacteriia

To investigate the evolution of HG biosynthesis within cyanobacteria, we searched for characteristic genetic signatures in 3,579 publicly available cyanobacterial genomes and plasmids^27^ and 14 newly sequenced genomes of heterocytous cyanobacteria (this study, see Supplementary Tables 1-3, Supplementary Data 1 and 2) and focused our analysis on photosynthetic cyanobacteria (taxonomic class *Cyanobacteriia* within the phylum *Cyanobacteriota*). We screened for the genomic presence and colocalization of 19 protein-coding sequences which are clustered on the genome of *Anabaena* sp. PCC 7120 (Fig. 1c, Supplementary Fig. 1; refs. ^13,28^), the well-studied ‘model’ cyanobacterium for heterocyte formation. The encoded proteins are involved in the biosynthesis of HGs, and in their export across the inner membrane and cell wall—leading to HG deposition in the cell envelope just outside the outer membrane and beneath the polysaccharide layer (Supplementary Table 4, refs. ^6,13–15,29,30^). This genomic cluster has been called the ‘*hgl* island’^13,28^ because it contains, among others, several ‘*hgl*’ genes (Fig. 1c). Here, we defined *hgl* islands as gene clusters containing homologs of at least 7 of the 19 queried genes (Supplementary Information).

To validate our genomic ‘island’ identification approach, we evaluated our results against a subset of cyanobacteria with known morphology (Supplementary Fig. 2, ref. ^31^), and analyzed the genomic co-occurrence of two processes taking place in heterocytes, which are both encoded by genomic clusters, HG and nitrogenase biosynthesis (the latter encoded by *nif* genes, Supplementary Tables 4 and 5, Supplementary Fig. 3). This benchmark confirmed that the ability to make heterocytes can be confidently inferred from the genome sequence based on the presence of an *hgl* island as defined here (Supplementary Information, Supplementary Table 6).

### Hgl and hgl-like islands are present throughout Cyanobacteriia

We investigated the presence of *hgl* islands in the context of the phylogeny of the *Cyanobacteriia* based on 24 concatenated core genes (Fig. 2a). This revealed a monophyletic group that includes cyanobacteria known to produce heterocytes. Although the morphology of many of the 479 genomes in this clade is unknown, 367 (77%) contain an *hgl* island composed of homologs of ≥10 of the queried genes (Fig. 1c), and almost all can fix nitrogen based on the presence of a *nif* island (Fig. 2a). This confirms previous suggestions of the occurrence of a single heterocytous clade within the *Cyanobacteriia*^21,31^.

**Fig. 2.**
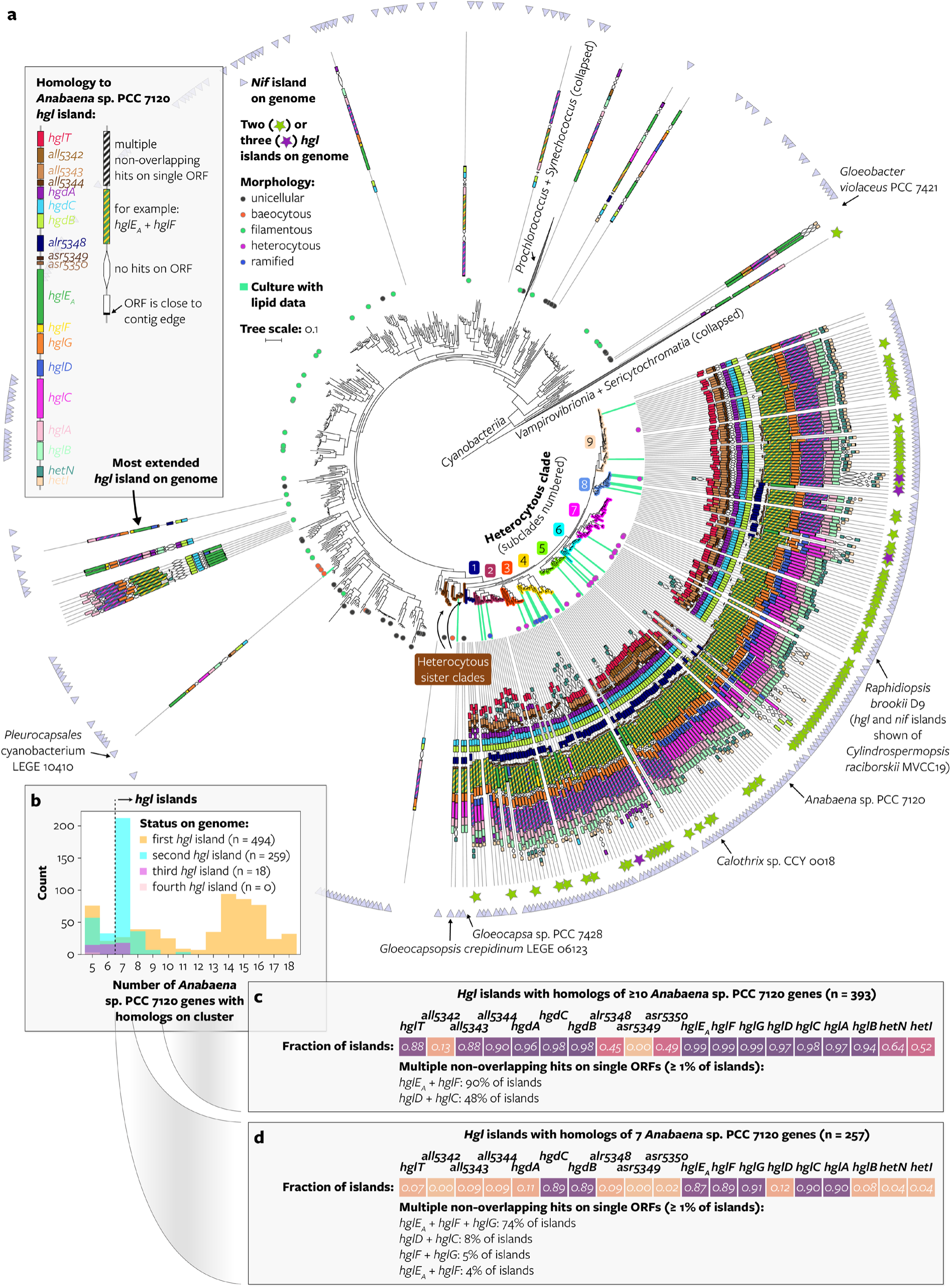
**a,** Maximum likelihood cyanobacterial phylogeny created using a concatenated alignment of 24 core vertically-transferred genes*^42^* of 1,260 genomes representing species groups (2,758 genomes of cultured and uncultured cyanobacteria clustered at ≥95% average nucleotide identity). The tree is rooted between *Cyanobacteriia* and the non-photosynthetic *Vampirovibrionia* and *Sericytochromatia^43,44^*. Morphology of specific genomes is indicated based on ref. *^31^*: unicellular (grey); baeocytous (orange), strains capable of forming internal small cells (or endospores) by multiple fissions in parent cells*^45^*; filamentous (green), chain of cells (or trichome) without an investing sheath which grows by intercalary cell division*^45,46^*; heterocytous (purple) and ramified (blue) are filamentous strains that when grown in absence of combined nitrogen contain heterocytes in their trichome and that divide in one or in more than one plane, respectively*^46^*. HG lipid data was obtained in this study and in previous studies (Supplementary Table 1). The *hgl* and *nif* islands of one selected genome per branch are drawn on the tree (Online methods). The composition of the most extended *hgl* island in terms of number of *Anabaena* sp. PCC 7120 genes with homologs on the island is shown. For size reference, the length of the coding region of *hglE_A_* of *Anabaena* sp. PCC 7120 is 4,626 base pairs. Genomes that contain more than one *hgl* island are indicated with stars, the composition of their additional islands is not shown. The ‘heterocytous clade’ is defined based on the ubiquitous presence of both *hgl* and *nif* islands and contains genomes with unknown morphology. Monophyletic heterocytous subclades were manually selected. Scale bar represents the mean number of substitutions per site. ORF, open reading frame; contig, contiguous sequence. Source files in Supplementary Data 3. **b**, Size distribution of all identified gene clusters. Gene clusters are grouped according to their status in the genome, where the ‘first’ cluster (orange) is the most extended island in terms of number of *Anabaena* sp. PCC 7120 genes with homologs on the island, and the ‘second’ cluster (cyan) the second-most extended island. Sample sizes in the legend are only for *hgl* islands (here defined as genomic clusters containing homologs of ≥7 *Anabaena* sp. PCC 7120 genes). **c**, Frequency of homologs of HG biosynthesis genes on *hgl* islands containing homologs of ≥10 genes, and **d**, containing homologs of 7 genes. Color-coding in panels c and d is for legibility only and reflects the numbers within the cells, with darker colors representing a higher fraction.

The gene composition of *hgl* islands with homologs of ≥10 genes is largely conserved, with homologs of 13 out of the 19 queried genes present on ≥88% of the islands (Fig. 2b,c), and their relative position fixed within the island (Supplementary Fig. 4). This indicates that evolutionary conservation of genes and their location within the *hgl* island are associated with the preservation of their function in heterocyte formation. Queried genes that are not conserved may be non-essential for HG biosynthesis, be essential but present somewhere else on the genome (e.g. *hetI*, see Supplementary Information and Supplementary Fig. 5), or their catalytic function may have been replaced by unrelated genes (Supplementary Information). The identification of 13 genes that are evolutionary conserved on *hgl* islands is in good agreement with previous mutagenesis studies showing that 12 of the genes are essential in *Anabaena* sp. PCC 7120 for proper HG biosynthesis and deposition (*hgdACB*, *hglTE_A_FGDCAB*, and *all5343*; Supplementary Information, Supplementary Table 4; refs. ^13,32–34^).

Besides, the genomes of 35 out of 2,279 cyanobacteria that are not part of the heterocytous clade— and are thus not expected to make heterocytes—remarkably also encode an *hgl* island by our definition (Fig. 2a, Supplementary Table 7). They include unicellular cyanobacteria like *Gloeocapsa* sp. PCC 7428, part of a closely related sister clade of heterocytous cyanobacteria in our phylogeny (Fig. 2a), and *Gloeobacter violaceus* PCC 7421 (ref. ^13^), a distantly-related cyanobacterium found at the base of the *Cyanobacteriia* (Fig. 2a), which has a uniquely simple life cycle and cellular composition lacking thylakoids^35,36^. Some of the non-heterocytous cyanobacteria possessing an *hgl* island do not encode a *nif* island (Fig. 2a), and are thus likely non-diazotrophic. Most of the *hgl* islands of non-heterocytous cyanobacteria have a more variable gene composition than the *hgl* islands of heterocytous cyanobacteria, with occasional gene duplications, fusions, rearrangements, and absences, and, importantly, all lack homologs of the gene *hglT* (Fig. 2a) encoding the glycosyltransferase (GT) that attaches the glucose headgroup to the AG in the last step of HG biosynthesis (Supplementary Fig. 1). Hence, although these cyanobacteria possess clusters of homologs of HG biosynthesis genes, their divergent gene composition and phylogenetic position outside the heterocytous clade suggests enigmatic biosynthetic products and a function unrelated to heterocyte formation.

Our genomic screening also revealed that 259 of the 494 *Cyanobacteriia* that have an *hgl* island possess two or more *hgl* islands (Fig. 2a,b). Many of the cyanobacteria with multiple islands encode an island with a highly conserved gene composition, containing homologs of only 7 HG biosynthesis genes (*hgdCB* and *hglE_A_FGCA*) encoded by 5 open reading frames (ORFs) (Fig. 2d), thus differing from the more extended *hgl* islands with homologs of ≥10 genes discussed above (cf. Fig. 2c). Additional *hgl* islands have previously been reported for four heterocytous cyanobacteria^37^, however our data show they occur widespread within heterocytous cyanobacteria. Because its gene composition is highly conserved and more limited in size compared to the queried island it may encode biosynthetic products other than HGs, and we call this island within heterocytous cyanobacteria the ‘*hgl*-like’ island hereafter. *Hgl*-like islands are absent in some akinete-forming heterocytous cyanobacteria like *Anabaena variabilis* ATCC 29413, and present as the only island in a few cyanobacteria within the heterocytous clade which have lost the ability to form heterocytes like *Raphidiopsis brookii* D9 (refs. ^23,38^), suggesting that the biosynthetic products of *hgl*-like islands are not involved in akinete nor heterocyte formation (Supplementary Information).

Hence, our results demonstrate that HG biosynthesis genes are conserved within a monophyletic heterocytous clade in *Cyanobacteriia* as a cluster resembling the *Anabaena* sp. PCC 7120 *hgl* island. However, several distantly related non-heterocytous cyanobacteria also encode *hgl* islands with a more divergent gene composition, and various heterocytous cyanobacteria encode an additional ‘*hgl*-like’ island, providing genomic evidence to elucidate the origin of HGs in *Cyanobacteriia*.

### Heterocytous cyanobacteria produce a wide diversity of new HGs

To characterize the structural diversity of HGs produced by heterocytous cyanobacteria, we carried out the largest HG screen to date using ultra-high-performance liquid chromatography coupled with multistage high-resolution mass spectrometry (UHPLC-HRMS*^n^*). We analyzed the lipid extracts of 26 cyanobacterial cultures evenly sampled throughout the heterocytous phylogeny, all grown under nitrogen-limiting conditions (Figs. 2a and 3, Supplementary Fig. 6, Supplementary Table 1). We screened 88 HG structures with different chain lengths, headgroups, and functional groups on their alkyl chain (Supplementary Information, Supplementary Tables 8-11), 19 of which were previously reported in heterocytous cyanobacteria (refs. ^7,9,10,18,39,40^). This revealed the presence of 30 previously unreported HGs, increasing the number of described HGs to 49 (Supplementary Information, Supplementary Table 9, Supplementary Figs. 7-10). Previously, only HGs with an even number of carbon atoms in their alkyl chain had been described^8,11^, yet we also detect odd-numbered HGs (hexose HG_27_ diol and hexose HG_29_ keto-ol) albeit in small amounts. Although most HGs described thus far contain a hexose headgroup, we discovered HGs with various other headgroups, including a still undetermined C_6_ sugar. In addition, our broad screening reveals novel chain length and headgroup combinations. The functional groups on the alkyl chain of the HGs are usually diols, keto-ols, keto-diols, or triols, but we also observe novel combinations of keto and alcohol groups such as diketo-ols, monofunctional HGs, and an HG with 4 functional groups. The identification of these new lipids highlights the wide diversity of HGs that are produced by heterocytous cyanobacteria today and opens a new avenue in the study of the role of HGs in heterocyte impermeabilization and its link to nitrogen fixation.

**Fig. 3.**
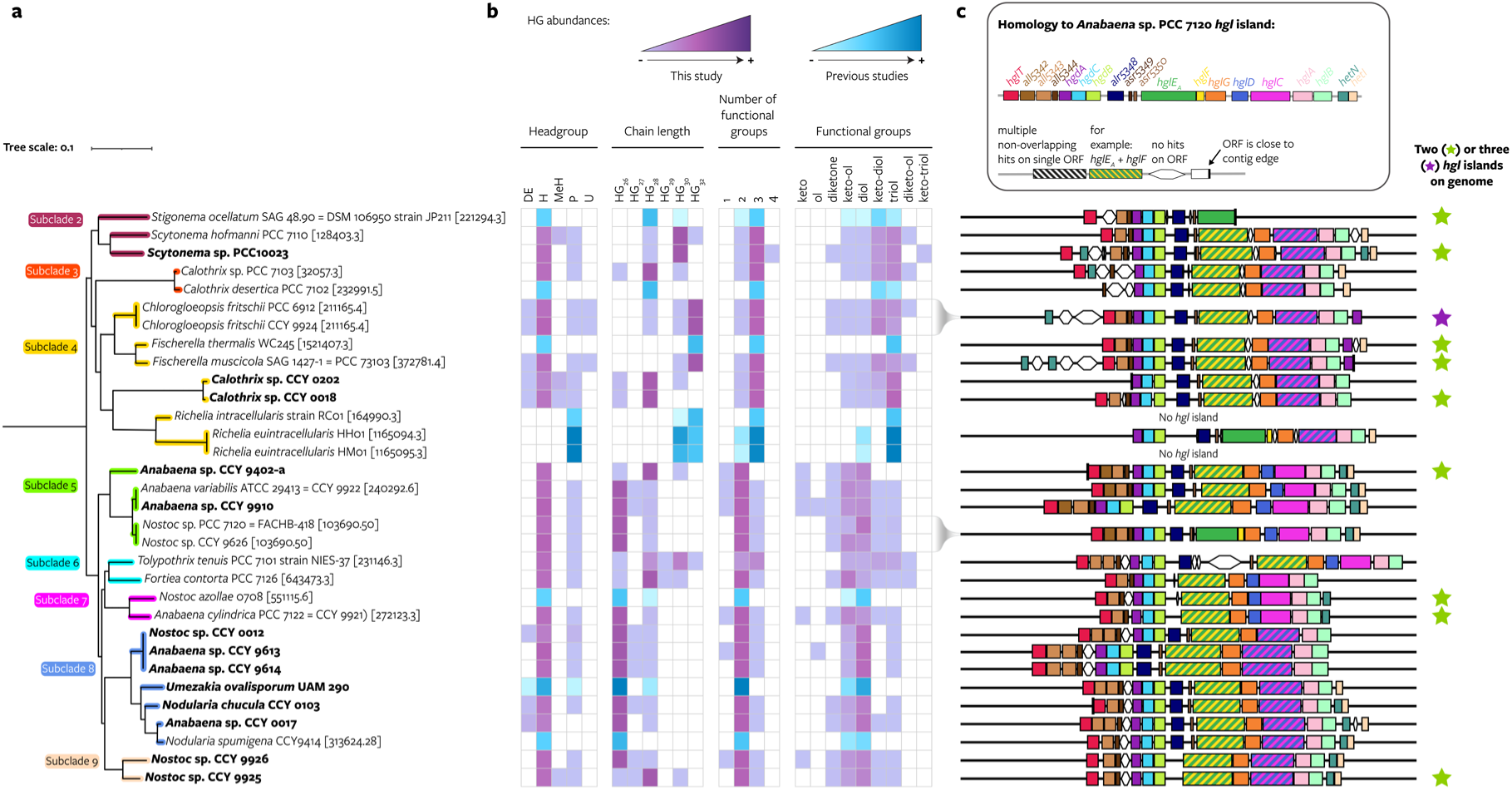
Distribution of 49 HGs grouped according to their structure’s characteristics throughout heterocytous *Cyanobacteriia*. **a**, Maximum likelihood cyanobacterial phylogeny based on 24 core vertically transferred genes including genomes with a known HG lipid profile (pruned from Fig. 2a). Genomes sequenced in this study are shown in bold. Scale bar represents the mean number of substitutions per site. **b**, Heatmap of HG relative abundances grouped according to the headgroup (H, hexose; DeH, deoxyhexose; MeH, methyl-hexose; P, pentose; U, unknown), chain length, number and type of functional groups of each HG. The same figure with all individual HGs is shown in Supplementary Fig. 6. HG abundances obtained from this study are shown in purple and those obtained from literature are shown in blue. **c**, Schematic representation of the genes present on the most extended *hgl* island on the genome in terms of number of *Anabaena* sp. PCC 7120 genes with homologs on the island. Stars indicate the presence of two (green) or three (purple) *hgl* islands on the genome; these additional islands are not drawn. ORF, open reading frame; contig, contiguous sequence.

### HG structure is mostly independent of hgl island gene composition

We next analyzed the distribution of HGs grouped according to their structure’s characteristics— headgroup, alkyl chain length, and type and number of functional groups on their alkyl chain (Fig. 3). Most cyanobacteria produce predominant HGs with a single combination of headgroup and chain length and with 2 or 3 functional groups, and a variety of other HGs in low abundance (Fig. 3b). This confirms previous observations of heterocytous cyanobacteria harboring two predominant HGs that are structurally identical except for differences in the number and stereochemistry of keto and alcohol groups they carry^7^. For example, the two most abundant HGs produced by *Chlorogloeopsis fritschii* PCC 6912 are hexose HG_32_ keto-diol and hexose HG_32_ triol (this study and ref. ^7^). However, others such as *Tolypothrix tenuis* PCC 7101 produce multiple HGs with different headgroups, chain lengths, and number of functional groups in high abundance (Fig. 3b, Supplementary Information; this study and ref. ^7^).

No direct relationship was found between the type of HGs cyanobacteria produced and the composition of their (most extended) *hgl* island, which is similar across the various species (Fig. 3c, Supplementary Fig. 6). Moreover, we did not find specific HGs associated with the additional presence of an *hgl*-like island on the genome, further suggesting that the more compact *hgl*-like islands are not involved in HG biosynthesis. The observed variation in HG distributions (Fig. 3b) is thus likely due to differences in enzymatic activity of the proteins encoded by the *hgl* island (see section ‘HG structure evolved convergently’), available precursors, cell physiology, environmental conditions, or a combination of these factors. An exception is *hglT*, the glycosyltransferase (GT) that attaches the glucose headgroup to the AG (Supplementary Fig. 1), which is absent from the genomes of three *Richelia* species that form symbiotic relationships with marine diatoms (Supplementary Information, Supplementary Figs. 6 and 11). *Richelia euintracellularis* HM01 and *Richelia intracellularis* RC01 both produce only HGs with a pentose instead of the common hexose headgroup (Fig. 3b), which have been suggested as biomarkers for diatom-diazotroph associations^9,19^, and the HG profile of *Richelia rhizosoleniae* SC01 is unknown. Thus, the production of pentose HGs may be directly related to the absence of *hglT* from the *hgl* island, and possible replacement by an alternative GT with pentose specificity. We thoroughly searched for this alternative GT on the genome of *Richelia euintracellularis* HH01 and identified four candidate genes based on sequence homology to *hglT*, and functional annotation as GT together with gene localization close to the *hgl* island (see Supplementary information for details). However, heterologous expression of the four candidate genes in an *hglT*-deficient *Anabaena* sp. PCC 7120 strain did not result in production of pentose HGs (Supplementary Information, Supplementary Figs. 12-14, Supplementary Tables 12-16).

### HG structure evolved convergently

Next, we assessed whether the variation in HG production between cyanobacteria can be explained by evolutionary relatedness, by evaluating the observed HG structural diversity in the context of the core gene phylogeny (Fig. 3, Supplementary Fig. 6), which represents the evolutionary history of the vertically transferred part of the genome. Since clusters of HG biosynthesis genes can be horizontally transferred—as suggested by their identification on a plasmid (Supplementary Information, Supplementary Fig. 2)—and may thus have a different evolutionary history, we also constructed a phylogeny based on the concatenated alignment of 7 genes that are shared among most *hgl* and *hgl*-like islands (Fig. 4, Supplementary Fig. 15). In this phylogeny, the most extended *hgl* islands of heterocytous cyanobacteria cluster together distantly from the *hgl* islands of non-heterocytous cyanobacteria and *hgl*-like islands within heterocytous cyanobacteria and have a tree topology largely similar to that of the core vertically transferred genes (Fig. 5). This suggests a shared evolutionary history of the core genes and *hgl* islands within heterocytous cyanobacteria (Supplementary Information). The core gene and *hgl* gene phylogenies thus show similar HG distribution patterns (Fig. 3 versus Supplementary Fig. 16).

**Fig. 4.**
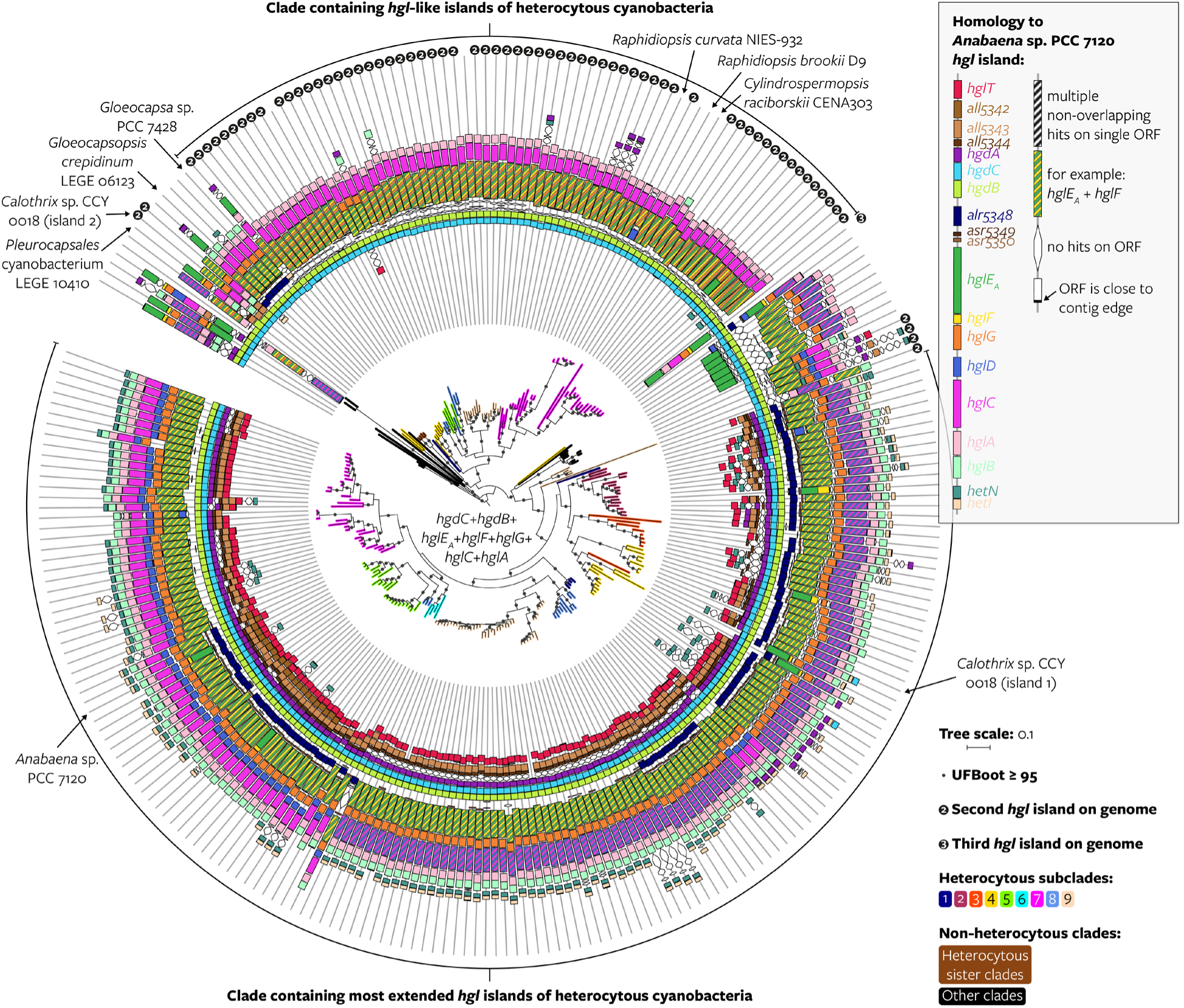
Evolutionary history of *hgl* islands within *Cyanobacteriia*. Maximum likelihood phylogeny of the *hgl* island created using a concatenated alignment of homologous sequences of 7 HG biosynthesis genes (*hgdCB* and *hglE_A_FGCA*) that are often present on *hgl* islands. Heterocytous and non-heterocytous subclades represent the position of the genome in the manually defined monophyletic clades of Fig. 2a based on a concatenation of 24 core vertically transferred genes. Genomes that are not part of a manually defined clade in Fig. 2a fall within the ‘other clades’ category in this figure. ‘First’, ‘second’, and ‘third’ *hgl* islands are based on the presence of other islands on the genome, where the ‘first’ *hgl* island (not marked) is the most extended island in terms of number of *Anabaena* sp. PCC 7120 genes with homologs on the island, the second *hgl* island (marked with a 2) the second-most extended island, and the third island (marked with 3) the third-most extended island. The *hgl*-like islands of *Raphidiopsis curvata* NIES-932, *Cylindrospermopsis raciborskii* CENA303, and *Raphidiopsis brookii* D9 are indicated because they are non-diazotrophic cyanobacteria from within the heterocytous clade. The tree is rooted at midpoint. Scale bar represents the mean number of substitutions per site. ORF, open reading frame; contig, contiguous sequence; UFBoot, ultrafast bootstrap approximation. Source files in Supplementary Data 4.

**Fig. 5.**
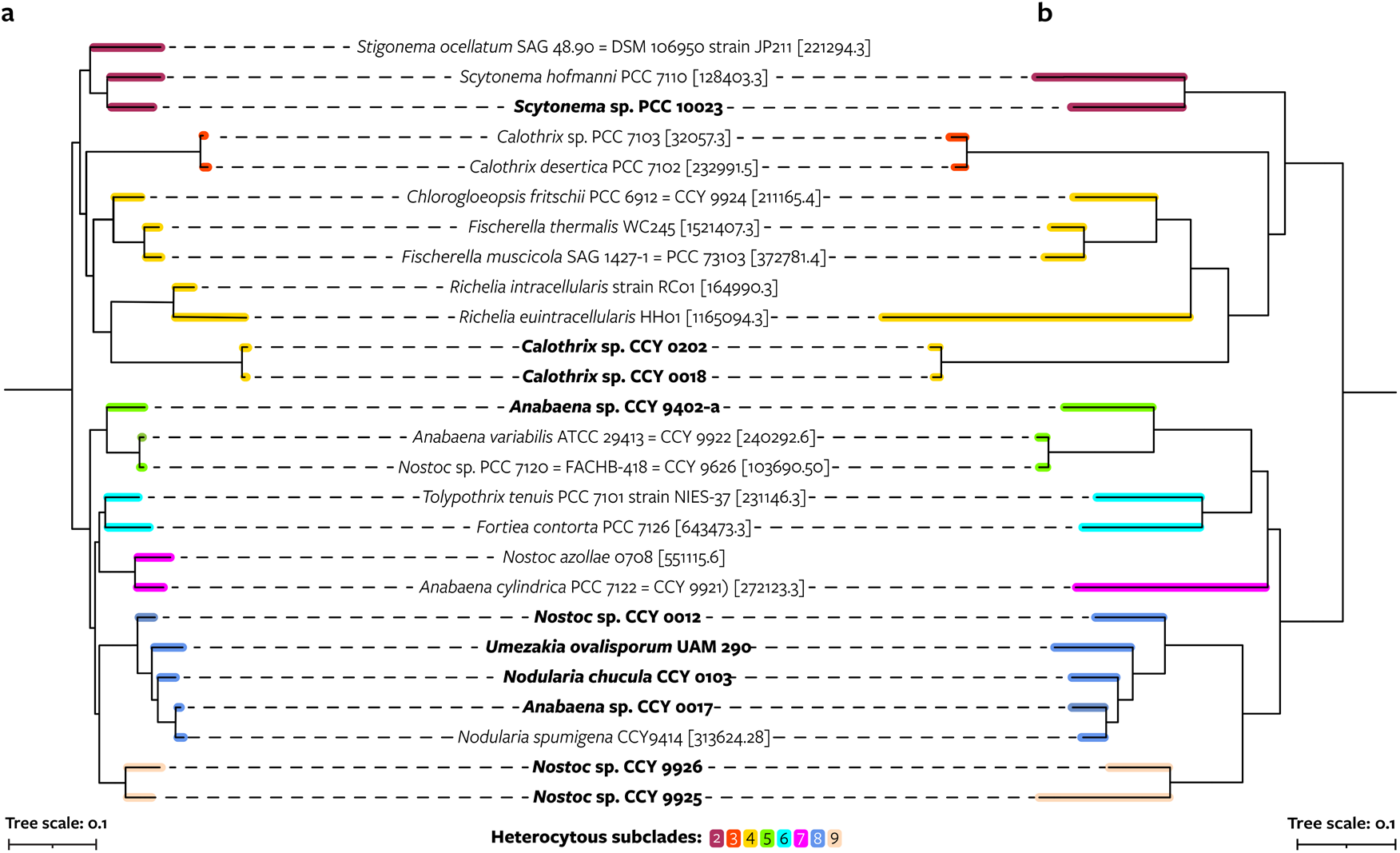
Comparison between maximum likelihood phylogenies based on 24 core vertically transferred genes and 7 *hgl* island genes of genomes with a known HG profile. **a**, Cyanobacterial phylogeny based on 24 core vertically transferred genes (pruned from Fig. 2a). **b**, Pruned phylogeny of the *hgl* island based on homologous sequences of 7 HG biosynthesis genes (*hgdCB* and *hglE_A_FGCA*) that are often present on *hgl* islands. For each cyanobacterium, only the most extended *hgl* island in terms of number of *Anabaena* sp. PCC 7120 genes with homologs on the island is shown. These *hgl* islands closely resemble the *hgl* island of *Anabaena* sp. PCC 7120 in terms of gene composition as opposed to the more compact *hgl*-like islands. The tree is rooted outside the depicted clade. Genomes sequenced in this study are shown in bold. No genomes from subclade 1 are present in the figure because this subclade does not contain cultured representatives with lipid data. Scale bars represent the mean number of substitutions per site.

Specific HGs are common within heterocytous subclades, such as the above discussed pentose HGs of two *Richelia* species. Strains from subclades 2-4 mainly produce dominant HGs with 3 functional groups and even-numbered alkyl chains with ≥28 carbon atoms. Strains from subclades 5-9 produce primarily HG_26_’s with 2 functional groups. However, few HGs are unique to a subclade, and most HGs—especially those that are dominant in one of the cultures—are found throughout the phylogeny even if in low abundance (Fig. 3, Supplementary Figs. 6 and 16). For example, the pentose headgroup, which was previously detected in only a few strains, is present in most cultures in relatively low abundance, although possibly in pentopyranose form (Supplementary Information). Exceptions are HGs with a single functional group and odd-chained HGs, which are both restricted to subclades 5-9, and pentose HG_30_ keto-triol—only found in *Scytonema* sp. PCC 10023 (Supplementary Fig. 6). In addition, although cultures from closely related strains usually produce similar HGs, this is not always the case. For example, *Nostoc* sp. CCY 9925 and *Nostoc* sp. CCY 9926 synthesize primarily hexose HG_28_ and hexose HG_26_, respectively.

Thus, the same HGs can be found in distantly related strains and closely related strains can differ in their abundant HGs, hence challenging the common use of HGs as taxonomic biomarkers, for example for members of the *Nostocaceae* and *Aphanizomenonaceae* families and for the genus *Fortieaceae* (Supplementary Information, Supplementary Table 17). These findings suggest that the biosynthetic process leading to HGs is flexible and can evolve convergently, i.e. independent evolution of the same HG structures in different taxonomic groups (Supplementary Information), possibly reflecting similar environmental pressures, leading to a chemically diverse pool of HGs to accommodate them.

### Hgl islands predate heterocyte formation

The identification of *hgl* islands in non-heterocytous cyanobacteria and of *hgl*-like islands allowed for a reconstruction of the acquisition of HG biosynthesis within the *Cyanobacteriia*. As discussed above, the most extended *hgl* islands of heterocytous cyanobacteria cluster together in the phylogeny based on 7 HG biosynthesis genes, as do their *hgl*-like islands (Fig. 4), and topologies of both phylogenetic clusters resemble that of the core genes when a root is placed in between, suggesting that a duplication that gave rise to both island types in heterocytous cyanobacteria predates the Last Heterocytous Cyanobacterial Common Ancestor (LHeCCA) (Supplementary Information). Phylogenetic placement of the *hgl* islands of non-heterocytous cyanobacteria—in between the two heterocytous phylogenetic clusters—suggests they were not acquired via recent horizontal transfer from heterocytous cyanobacteria (Fig. 4 and Supplementary Fig. 17, Supplementary Table 18) and supports the presence of *hgl* islands further back in time. Given that *Gloeobacter violaceus* PCC 7421 contains an *hgl* island, the duplication may predate thylakoids or the last common ancestor of *Cyanobacteriia*. Alternatively, multiple horizontal transfer events predating LHeCCA may have given rise to the *hgl* islands in non-heterocytous cyanobacteria (Supplementary Information). In both cases, widespread loss explains the sparse distribution of *hgl* islands in *Cyanobacteriia* today. Hence, the *hgl* islands in specific non-heterocytous cyanobacteria likely represent a phylogenetic lineage of the island from before the emergence of the heterocyte. Their diverse gene compositions suggest that this ancestral island already contained many homologs of the contemporary HG biosynthesis genes—except for possibly *hglT* which is absent from all non-heterocytous *hgl* islands—and reflect subsequent differential loss of individual genes.

### Potential origin of HGs from ancient 1,3-diols

To elucidate the evolutionary origin of HG biosynthesis, we selected two cultured strains from outside the heterocytous clade that contain an *hgl* island and are capable of nitrogen fixation— *Pleurocapsales* cyanobacterium LEGE 10410 and *Gloeocapsopsis crepidinum* LEGE 06123 (Fig. 6a,b)—and exposed them to heterocyte- and akinete-inducing stresses, namely nitrogen deficiency and ageing^26,41^. Microscopic analysis revealed that upon division, daughter cells of both strains remain attached to the parent cell and are surrounded by a polysaccharide layer, thus appearing as aggregates (Supplementary Fig. 18), under all growth conditions. Because their *hgl* islands lack *hglT* homologs and contain potential ABC transporters expressed by the *hgdBC* homologs, we hypothesize they can be involved in the biosynthesis and export of AGs.

**Fig. 6.**
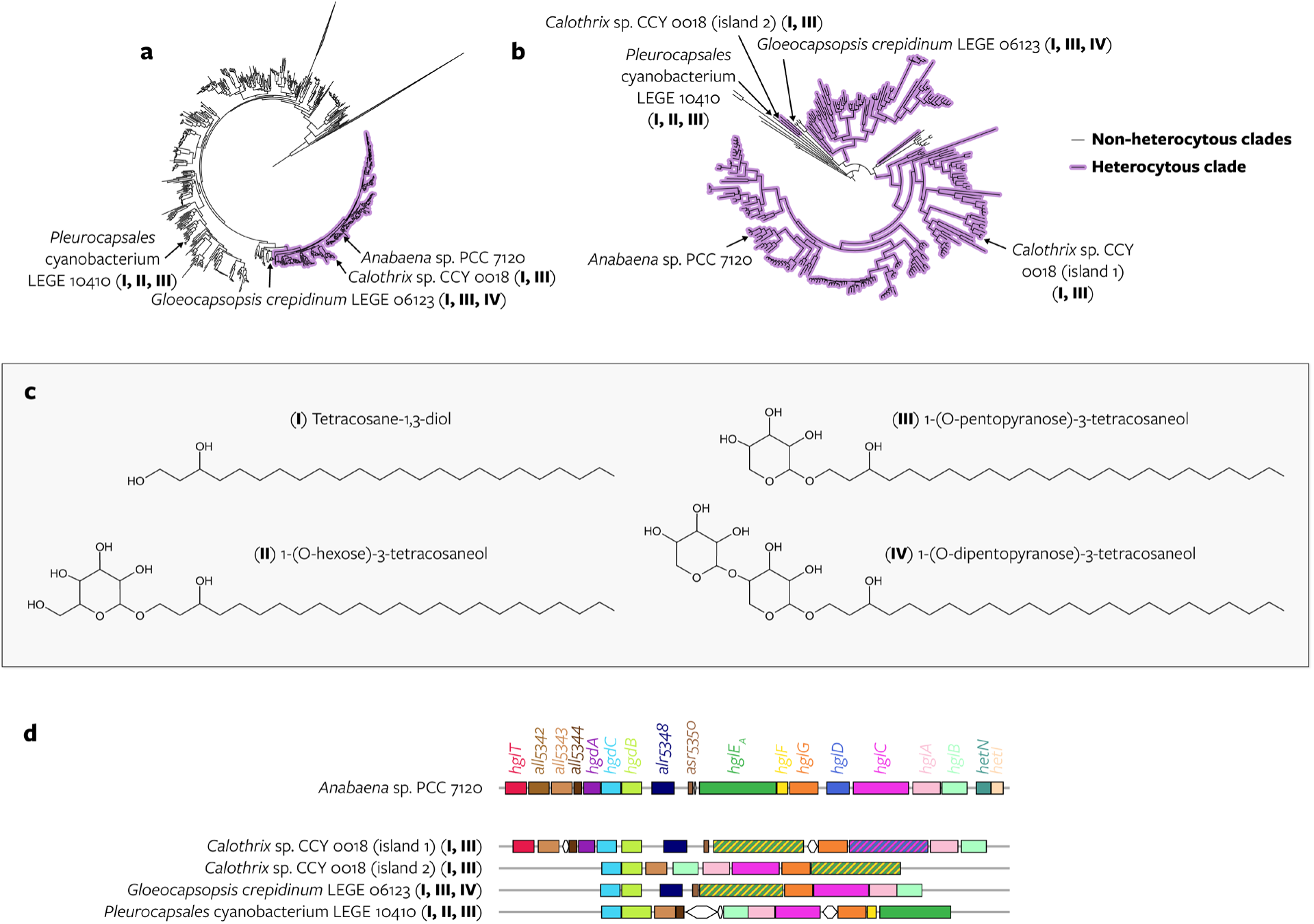
Two non-heterocytous and one heterocytous cyanobacteria produce HG analogs. **a**, Position of cyanobacteria that produce HG analogs in the phylogeny based on the concatenated alignment of 24 core vertically-transferred genes (Fig. 2a). The HG analogs that they produce are depicted in panel c and indicated with roman numerals I-IV throughout the figure. *Pleurocapsales* cyanobacterium LEGE 10410 and *Gloeocapsopsis crepidinum* LEGE 06123 are the only non-heterocytous cyanobacteria whose lipids were analyzed. *Anabaena* sp. PCC 7120 does not produce HG analogs but is indicated for reference. **b**, Position of the *hgl* islands of the same cyanobacteria in the concatenated alignment of homologous sequences of 7 HG biosynthesis genes (Fig. 4). Both *hgl* islands of *Calothrix* sp. CCY 0018 are indicated. **c**, Tentative structure of HG analogs. **d**, *Hgl* islands encoded on the genomes of *Anabaena* sp. PCC 7120 and the three cyanobacteria that produce HG analogs.

Although no C_26_ - C_32_ HGs nor derived AGs were detected, both strains synthesize novel lipids that structurally resemble canonical HGs and AGs, but with a shorter C_24_ alkyl chain and lacking a characteristic functional group at the ω-1 position (Supplementary Information, Supplementary Figs. 19-22). Whilst some are observed in diol form (tetracosane-1,3-diol), thus resembling AGs, others consist of this same C_24_ alkyl chain compound bound to different sugars (1-(O-hexose)-3-tetracosaneol, 1-(O-pentopyranose)-3-tetracosaneol, and 1-(O-dipentopyranose)-3-tetracosaneol) resembling HGs (hereafter ‘HG analogs’) (Fig. 6c, Supplementary Tables 11, 19 and 20). We speculate that these novel lipids are likely produced by the enzymes encoded by the *hgl* islands of the non-heterocytous strains (Fig. 6d, Supplementary Information). Analysis of the lipid extracts of the heterocytous cultures revealed that *Calothrix* sp. CCY 0018, which encodes an additional *hgl* island closely related to the *hgl* island of *Pleurocapsales* cyanobacterium LEGE 10410 (Fig. 6b), also produces HG analogs (Supplementary Fig. 23, Supplementary Table 11), providing further evidence for the biosynthesis of these lipids by an ancient phylogenetic lineage of the *hgl* island (Supplementary Information).

As the ancestor of these islands was likely present in *Cyanobacteriia* before the emergence of the heterocyte (see above), we propose that LHeCCA was already capable of synthesizing and exporting C_24_ alkyl chain lipids resembling HGs and AGs for still unknown purposes (Supplementary Information). Because these lipids were detected under all growth conditions including in non-limited media containing nitrogen (therefore not promoting nitrogen fixation), and because *hgl* islands are also present in non-diazotrophic cyanobacteria outside the heterocytous clade (Fig. 2a, Supplementary Table 7) the HG analogs produced by contemporary cyanobacteria, and conceivably by extension by LHeCCA, are probably not involved in nitrogen fixation nor in (a process similar to) akinete formation.

In this scenario, the modified biosynthetic steps encoded by one of the copies of the *hgl* island of LHeCCA would have allowed for the emergence of HGs from HG analogs by chain length extension and the addition of new keto, alcohol, and methyl groups (Supplementary Information), and facilitated their new role as nitrogenase protectors in an increasingly oxygenated atmosphere. The absence of contemporary HGs with a C_24_ alkyl chain may reflect selection for longer chain length potentially providing a higher oxygen impermeability to the heterocyte membrane throughout the course of the evolution of heterocytous cyanobacteria.

## Conclusions

The broad screening of HG composition in cultures revealed a large and previously unknown structural diversity of HGs within heterocytous cyanobacteria, exposing 30 novel HGs—nearly tripling known HG structural diversity. Our findings also highlight the reduced taxonomic value of HGs, which complicates their use to identify their producers in present and past environments.

Our results support the origin of HGs from ancient HG analogs that were not involved in nitrogen fixation—remnants of which may still be found in specific non-heterocytous and heterocytous cyanobacteria today. Genomic clusters of HG biosynthesis genes were probably present in *Cyanobacteriia* before the rise of the heterocyte. Neofunctionalization of this existing genomic machinery for the production and export of 1,3-diols and related sugar-bound lipids allowed for the biosynthesis of HGs and the formation of an oxygen-impermeable membrane layer to further protect nitrogenase, the key enzyme for nitrogen fixation. These findings suggest the usefulness of AG, HGs, and HG analogs to trace the evolution of multicellularity and division of labor in cyanobacteria, which remains challenging to pinpoint due to scarcity of fossil evidence. Future studies will need to evaluate the preservation potential and diagenetic transformations of these molecules to fully conclude their use as fossils for molecular clock calculations of key events in Earth’s history and microbial evolution.

## Data availability

Sequencing data and genome assemblies were deposited in the European Nucleotide Archive (ENA) under project number PRJEB67822. UHPLC-HRMS*^n^*(Orbitrap) and GC-MS datafiles, phylogenetic trees raw files and plasmid maps can be found in supplied Supplementary Data Files and/or in the Zenodo repository https://doi.org/10.5281/zenodo.10444558.

## Code availability

The code used for this manuscript is available from Zenodo at https://doi.org/10.5281/zenodo.10444558 and can be used to reconstruct all results presented here.

## Acknowledgments

We thank Samuel Cirés for providing biomass of the *Umezakia ovalisporum* strains UAM 290 and UAM 292, Koichiro Awai for providing the strains *Anabaena* sp. PCC 7120 *wt* and Δ*hglT*, Michele Grego and Eveline Garritsen for assistance handling the cyanobacterial cultures, Anchelique Mets and Jessica Riekenberg for assistance during lipid extraction, Denise Dorhout and Monique Verweij for running and maintaining the UHPLC-HRMS*^n^* analytic equipment, Marcel van der Meer for assistance on the interpretation of the GC-MS data, Judith van Bleijswijk for arranging the required permits to work with *Anabaena* sp. PCC 7120 mutants, Lora Strack van Schijndel for assistance extracting DNA from *Anabaena* sp. PCC 7120 mutants, Aniek van der Woude for helpful cloning suggestions, and Tracy Villareal and Peter C. Wolk for helpful insights on heterocytous cyanobacteria.

## Online Methods

### Bacterial strains and growth conditions

Most cyanobacterial biomass from heterocytous strains was obtained from frozen (−80 °C) culture pellets stored in the former Culture Collection Yerseke (CCY), and from starter cultures provided by the Pasteur Culture Collection (PCC) (Supplementary Tables 1 and 2). The exceptions were *Umezakia ovalisporum* strains UAM290^47^ and UAM292 (formerly known as *Chrysosporum ovalisporum* or *Aphanizomenon ovalisporum*), kindly provided by Dr. Samuel Cirés, and *Anabaena* sp. PCC 7120 *wild type* (*wt*) and Δ*hglT*^15^, kindly provided by Dr. Koichiro Awai. Non-heterocytous strains were kindly provided by the Blue Biotechnology and Ecotoxicology Culture Collection (LEGE-CC)^48^.

*Escherichia coli* strains NEB® 5-alpha (New England Biolabs) and TOP10 (ThermoFisher Scientific) were used for plasmid manipulation, grown at 37 °C in LB (Miller) Broth (Sigma Aldrich, St. Louis, MO, USA) or in solid LB (Miller) supplemented with bacteriological agar (VWR, Solon, OH, USA). pRL443 (Addgene plasmid # 70261; http://n2t.net/addgene:70261; RRID:Addgene_70261) and pRL623 (Addgene plasmid # 58494; http://n2t.net/addgene:58494; RRID:Addgene_58494) were a gift from Peter Wolk, and pAM5404 was a gift from Susan Golden (Addgene plasmid # 132660; http://n2t.net/addgene:132660; RRID:Addgene_132660) obtained through Addgene (Watertown, MA, USA).

*Anabaena* sp. PCC 7120 *wild-type (wt)* and *ΔhglT*^15^ were routinely grown at 30 °C in liquid BG11 medium or BG11 without nitrogen (BG11_0_)^46^ (see Supplementary Table 2), supplemented with appropriate antibiotics, and incubated under constant white light illumination (60-70 μE/m^2^/s) with or without agitation at 120 rpm (INFORS, Bottmingen, Switzerland). Cells were also grown in solid BG11 medium supplemented with bacteriological agar (VWR, Solon, OH, USA) and the appropriate antibiotics. When required, the following antibiotic concentrations were used: ampicillin (100 μg/ml), kanamycin (20 μg/ml), spectinomycin (25 μg/ml), streptomycin (10 μg/ml), and chloramphenicol (25 μg/ml). Growth of *Anabaena* strains was monitored by measuring OD_730_ in a V-1200 Spectrophotometer (VWR International Europe, Leuven, Belgium). To induce heterocyte formation, cells were either grown directly in BG11_0_ media or transferred from BG11 to BG11_0_ by either: (1) washing the culture three times with BG11_0_ (*wt* and Δ*hglT*) (2) a minimum of five successive 1:10 culture dilutions in BG11_0_ (ARP001 and ARP003), (3) two to five rounds of cell collection and transfer to BG11_0_ medium (ARP002n, ARP004-007).

*Gloeocapsopsis crepidinum* LEGE 06123 and *Pleurocapsales* cyanobacterium LEGE 10410 were routinely grown at 21 °C in MN and MN_0_ media (see Supplementary Table 2) and incubated under a 12:12 light:dark regime, with white light illumination (30-40 μE/m^2^/s) and without agitation (SANYO MLR-350, Japan). Cultures were either grown directly in MN media or transferred from MN to MN_0_ using a 1:10 dilution. After three successive transfers in MN_0_, cultures were incubated for 38 days and then harvested. To evaluate the effect of other stressors such as nutrient limitation and culture age, the first transfer of the cultures was also harvested after 77 days.

Biomass was harvested by centrifugation at 2500 rpm for 10 min, washed 3 times with bi-distilled water and stored at −20 °C until further processing.

### Microscopic analysis

Prior to harvesting, strains *Gloeocapsopsis crepidinum* LEGE 06123 and *Pleurocapsales* cyanobacterium LEGE 10410 were analyzed under the microscope. To visualize the presence of external polysaccharide a fresh aliquot of each culture was incubated with an equal volume of a 0.5% solution of Alcian blue (Sigma Aldrich, Steinheim, Germany) at room temperature for 10 min, which was then visualized using bright field microscopy (Axio Imager M2, Zeiss, magnification x200 - x400). Cell lipids were visualized using Nile Red (Sigma Aldrich, Steinheim, Germany), in brief, 40 µl of culture were incubated with 50 µl DMSO 40% and 10 µl Nile Red (10 µlg/ µl in 40% DMSO) at room temperature for 5 to 15 min. Stained samples were then visualized using epifluorescence microscopy on an Axio Imager M2 microscope (Zeiss, magnification x400) coupled to a Colibri LED light source (Zeiss, 555 nm excitation wavelength).

### Nucleic acids extraction and sequencing

Genomic DNA (gDNA) was extracted from strains described in Supplementary Table 2 using the DNeasy PowerSoil kit (Qiagen, Hilden, Germany) with minor modifications, namely: samples were disrupted using the bead mill homogenizer ran twice for 10 seconds at a speed of 3.55 m s^-1^ with a 30 second dwell. Concentration of genomic gDNA extracts was analyzed using a Nanodrop ND-1000 Spectrophotometer (NanoDrop Technologies Inc. Wilmington, DE, USA). gDNA was sequenced using Illumina sequencing platform 150bp paired-end, achieving total of 13 to 22 million reads per strain. Library preparation using Illumina TruSeq Nano (350bp insert) library kit and sequencing were carried out by Macrogen Europe (Amsterdam, The Netherlands), whilst library preparation using Nextera XT DNA Library Preparation Kit (96 samples) and sequencing were carried out by Macrogen Korea (Seoul, Republic of Korea) (Supplementary Table 4).

### Genome assemblies

Adapter sequences and polyG tails were removed from Illumina reads using Trimmomatic (v0.36; using arguments ‘PE’, ‘-phred33’, and ‘ILLUMINACLIP: adapters.fa:2:30:10 LEADING:20 TRAILING:20 SLIDINGWINDOW:5:20 MINLEN:40’)^49^ and Cutadapt (v1.16; using arguments ‘--nextseq-trim=20’, ‘-m 40’, ‘-q 20’, ‘-n 5’, ‘--discard-trimmed’)^50^, respectively. Sequence quality was checked using FastQC (v0.11.9)^51^. Sequencing reads were assembled into draft genomes using BiosyntheticSPAdes (v3.14.1)^52^. Quality and potential contamination of the assemblies were assessed with BlobTools2 (v2.3.3)^53^ integrating the output of BLASTN (v2.10.1+)^54,55^ to NCBI nt downloaded on April 1^st^ 2021, DIAMOND blastx (v2.0.8.146)^56^ to UniProt 2021_01 database^57^, Minimap2 (v2.17-r941)^58^ to map the reads back to the assembly and BUSCO (v5.1.1)^59^ with database version odb10. Only contiguous sequences (contigs) assigned as ‘Cyanobacteria’, not assigned to any phylum (‘no-hit’), and to ‘Bacterial-undefined’ were retained. To confirm the absence of contigs of other phyla in our assemblies we used BlobTools2 integrating data of BLASTN, Minimap2 and BUSCO. This procedure was repeated for four genomes (CCY 9926, CCY 0103, UAM 290 and UAM 292) where proteobacterial contigs had been detected. Additionally in two of these genomes (UAM290 and UAM292) only contigs with a minimum coverage of 3x and a GC percentage of 30-50% were retained.

### Selection of publicly available cyanobacterial genome sequences

We downloaded all genomes from the PATRIC genome database^60^ (currently part of the BV-BRC database^61^) that had a taxonomic assignment as ‘phylum *Cyanobacteria*’ (taxonomy ID 1117 according to NCBI taxonomy) based on the ‘genome_lineage’ file on the PATRIC ftp server (ftp.patricbrc.org) from April 30, 2022. As some of the genome sequences in PATRIC were identical, we identified replicates based on concatenated DNA sequences and concatenated sorted DNA sequences and kept one genome per group of replicates.

The final selection of 3,657 genomes (Supplementary Data 1) included 258 genomes with genome status ‘complete’, 3,021 with genome status ‘WGS’, 342 with genome status ‘Plasmid’ and 36 with unknown genome status. The genomes were taxonomically reannotated with GTDB-Tk (v2.1.0)^62^ using release207_v2 of the Genome Taxonomy Database (GTDB)^63^, revealing that most of the chromosomal genomes were from the class *Cyanobacteriia*. The annotation algorithm of GTDB-Tk is based on phylogenetic placement of core genome marker genes and thus does not allow for taxonomic annotation of plasmids. GTDB-Tk placed 52 and 5 genomes in the non-photosynthetic cyanobacterial classes *Vampirovibrionia* and *Sericytochromatia*, respectively, and 20 genomes outside cyanobacteria (Supplementary Data 1). Genome quality was assessed with CheckM (v1.1.3)^64^ in the lineage-specific workflow (Supplementary Data 1). Genomes from strains with known morphology (unicellular, baeocytous, filamentous, heterocytous, and ramified) based on ref. ^31^ were identified based on their taxonomy ID.

### Query of HG biosynthesis genes and *hgl* island definition

Proteins were predicted on the PATRIC genomes and on the 14 newly sequenced genomes with Prodigal (v2.6.3)^65^ in single genome mode. The 19 protein sequences that are encoded by the *hgl* island of *Anabaena* sp. PCC 7120 (see Supplementary Fig. 1) were extracted from the protein files of GCA000009705.1_ASM970v1 on GenBank^66,67^ and queried against a single concatenated protein file containing all proteins of the PATRIC genomes and of the 14 newly sequenced genomes with BLASTP (v2.12.0+)^54^. No protein-coding sequences were predicted on 78 PATRIC genomes, which were all plasmids according to the PATRIC metadata table, and the final set of proteins thus encompassed 3,579 genomes. Hits with an *e*-value ≤ 1^e-5^ and query coverage ≥ 50% were considered homologous.

Because of the potential of gene fusions and fissions, we allowed for multiple hits per open reading frame (ORF) if these hits did not overlap substantially. Hits were defined as ‘overlapping’ if at least one of the hits had ≥ 50% overlap on the subject sequence with another hit. For a group of overlapping hits, the single best hit was chosen according to bit-score. ‘Non-overlapping’ hits—i.e. hits that had < 50% overlap on the subject sequence with any other hit—where all considered present on the subject sequence. ORFs with hits to HG biosynthesis genes were considered part of a genomic cluster if they were separated by ≤3 ORFs on the contig. For example, the first predicted ORF and the fifth predicted ORF on a contig were considered connected by this definition, but not the first and the sixth unless an ORF in between also contained a hit to an HG biosynthesis gene. An ORF was considered close to a contig edge if it was within 3 ORFs of the edge, for example the fourth ORF on a contig was considered close to the edge but not the fifth. We examined genomic clusters containing homologs of at least 7 of the queried HG biosynthesis genes—i.e. a genomic cluster of ORFs with hits to at least 7 different HG biosynthesis genes irrespective of the number of ORFs on which they were encoded or the copy number of the gene on the cluster—as ‘*hgl* islands’.

### Query of *nif* genes and identification of glycosyltransferases

*Nif* genes and genomic clusters of *nif* genes were defined similar to the HG biosynthesis genes and genomic clusters described above. We queried nitrogenase protein sequences encoded by *nif* genes from *Anabaena* sp. PCC 7120 of GCA000009705.1_ASM970v1 that are part of the nitrogenase gene cluster depicted in Figure 2 of ref. ^68^: alr1407 (*nifV1*), asr1408 (*nifZ*), asr1409 (*nifT*), all1433 (*nifW*), all1436 (*nifX*), all1437 (*nifN*), all1438 (*nifE*), all1440 (*nifK*), all1454 (*nifD*), all1455 (*nifH*), all1456 (*nifU*), all1457 (*nifS*), alr1459 (*xisF*), and all1517 (*nifB*). Overlapping and non-overlapping hits were defined and dealt with as above for the HG biosynthesis genes, as were genomic clusters of ORFs. The length of a *nif* genomic cluster was defined as the number of queried genes with hits on the genomic cluster irrespective of the number of ORFs on which they were encoded or the copy number of the gene on the cluster. We considered genomic clusters containing homologs of at least 5 of the queried *nif* genes ‘*nif* islands’ and an indication of the genomic capability of nitrogen fixation.

To identify glycosyl transferases, we annotated the genomes with dbCAN2, using dbCAN HMMdb (v10.0)^69^ and hmmscan, with an *e*-value < 1e-15 and coverage > 0.35.

### Cyanobacterial core gene phylogeny

We selected the 2,777 genomes that were taxonomically annotated as ‘phylum *Cyanobacteria*’ according to GTDB-Tk, and had at least a medium-quality draft status according to the minimum information about metagenome-assembled genome (MIMAG) criteria^70^ (estimated completeness ≥ 50% and contamination < 10%). Similar genomes were identified with dRep (v3.4.0)^71^ using the fastANI algorithm and a similarity cut-off of 95% average nucleotide identity (ANI), which has been suggested as species-level boundary^72^.

The representative genome of each dRep cluster was used for constructing a phylogeny based on core genes. We started with the 27 Clusters of Orthologous Gene (COG) families that showed evidence of being primarily vertically transferred in ref. ^42^. For identification of these genes in the genomes, fasta files from the COG 2020 database were downloaded from the NCBI ftp server (ftp.ncbi.nih.gov/pub/COG/COG2020/data/fasta/), and aligned with MAFFT (v7.505)^73^ with the ‘--anysymbol’ flag, using the L-INS-i algorithm if the COG family contained ≤ 800 sequences and default parameters otherwise. HMM profiles were constructed with hmmbuild and hmmpress from the HMMER package (v3.2.2)^74^. The representative genomes, with proteins longer than 100.000 amino acids removed, were annotated with the COG database using hmmscan from the HMMER package and an *e*-value < 1e-5. The best hit was selected per protein sequence based on *e*-value. We counted the occurrence of the COG families in each genome and selected the 24 COG families that were present in single-copy in ≥ 75% of the representative genomes as our final set of core genes. Genes that were single-copy in a genome were extracted and aligned with MAFFT using the L-INS-i algorithm. The alignments were trimmed with trimAl (v1.4.rev15)^75^ in gappyout mode. If a sequence contained ≥ 50% gaps after trimming, it was removed from the alignment. The aligned sequences were concatenated per genome, filling in gaps when a gene was absent from the alignment—i.e. when it was not present on the genome or present in multiple copies or when it was removed after trimming. We removed the 18 representative genomes whose concatenated alignment was based on <7 genes. The final concatenated alignment contained 1,260 representative genomes (representing 2,758 genomes) and 6,933 amino acids.

A phylogenetic tree was constructed with IQ-TREE (v1.2.1)^76^, using 1,000 ultrafast bootstraps^77^, and model selection^78^ based on nuclear models. The best-fit model (LG+R10) was chosen based on the Bayesian Information Criterion (BIC). The phylogenetic tree was visualized and decorated in Interactive Tree of Life (iTOL)^79^. Pruning of the phylogenetic tree for figures that contained a selection of cyanobacteria was also done in iTOL.

### Visualization of *hgl* and *nif* islands on core gene phylogeny

The branches of the core gene phylogeny are representative genomes of dRep clusters that sometimes contain multiple genomes. We decorated each branch on the core gene phylogeny with *hgl* and *nif* islands of one selected genome, and the genome that was chosen for this visualization was not always the dRep representative genome that was used to construct the core gene phylogeny. When the dRep cluster contained a genome from a culture with associated lipid data, this genome was chosen. Alternatively, the genome from the dRep cluster that had the most extended *hgl* island in terms of number of queried genes with hits on the island was chosen.

The most extended *hgl* island in terms of number queried genes with hits on the island was drawn on the phylogeny. Presence and absence of a *nif* island in a branch of the core gene phylogeny was based on the same selected genome. Drawing direction of the *hgl* islands was based on the orientation of the *hgdC* / *hgdB* pair, or if the island had no hits to both genes of the *hglG* / *hglC* pair. The oriented *hgl* islands were aligned based on their nucleotide position relative to *hgdB* (no offset) or *hglC* (an offset of 10,135 nucleotides because this is the distance between *hgdB* and *hglG* on the genome of *Anabaena* sp. PCC 7120 (BV-BRC genome identifier 103690.50)).

### Phylogenies based on 7 *hgl* island genes

We extracted all hits of 7 HG biosynthesis genes—*hgdCB* and *hglEAFGCA*—from the 324 *hgl* islands that were drawn on the core gene phylogeny. Only the protein region of the BLASTP hit was extracted, and when an island contained multiple hits, we picked the best hit in terms of bit-score. 69 *hgl* islands did not contain a hit to all 7 genes and were discarded. The protein sequences were aligned with MAFFT using the E-INS-i algorithm, trimmed with trimAl in gappyout mode, and the aligned sequences were concatenated per *hgl* island. A phylogenetic tree was constructed with IQ-TREE, using 1,000 ultrafast bootstraps, and model selection based on nuclear models. The best-fit model (JTTDCMut+F+R8) was chosen based on the BIC. The phylogenetic tree was visualized and decorated in iTOL. Pruning of the tree to generate Fig. 5 and Supplementary Figs. 16, and 23 was done in iTOL.

In addition, we made a second phylogeny which included the same hits but also incorporated the hits from the *hgl* islands of 4 cyanobacteria that were not drawn on the core gene phylogeny (Fig. 2). These cyanobacteria were chosen because they are non-diazotrophic cyanobacteria from within the heterocytous clade (*Raphidiopsis curvata* NIES-932, *Cylindrospermopsis raciborskii* CENA303, and *Raphidiopsis brookii* D9) or because they possess an *hgl* island but without *hglT* homologs (*Cylindrospermopsis raciborskii* CYRF; the highest-scoring *hglT* hit anywhere on its genome has a bit-score of 68.6). For these 4 genomes, we included *hgl* islands with hits to at least 6 of the queried genes, and reconstructed an alignment and phylogenetic tree as described above, filling in gaps when a gene was absent from the alignment. The best-fit model (JTTDCMut+F+R9) was chosen based on the BIC. The phylogenetic tree was visualized and decorated in iTOL. Pruning of the tree to generate Supplementary Figs. 15 and 17 was done in iTOL.

### Cloning strategy and recombinant plasmid construction

Prior to further genetic modifications we confirmed the substitution of *hglT* for the *npt* kanamycin resistance cassette in the *ΔhglT Anabaena* strain via PCR using two different primer sets (Supplementary Fig. 12 and Supplementary Table 12). The presence of a kanamycin resistance cassette in place of *hglT* was analyzed using primers Fw3_Hali14 and Rv2_Hali14_mod ^15^, which produced the expected 990 bp fragment (data not shown). Whilst deletion of *hglT* in all genome copies of the *ΔhglT Anabaena* mutant strain was determined using primers HR1_hglT_F and HR2_hglT_R, if *hglT* was present a ∼1700 bp band would be seen, whist if *hglT* had been successfully replaced by *npt* we would observe a ∼2000 bp band (Supplementary Fig. 12). Our results confirmed that *hglT* had been successfully deleted in all genome copies. However, in addition to the expected ∼2000 bp band we also observed a smaller (∼550 bp) band (data not shown), which might be caused by the partial loss of the *npt* kanamycin cassette (∼800 bp) after kanamycin stepdown upon receival of the strain.

The PCRs described above were carried out using 1 µl gDNA extracted as described in the Material and Methods in a 25 µl polymerase chain reaction (PCR) containing, 10 µl PCR water, 5 µl Q solution, 2.5 µl Qiagen Buffer, 2 µl dNTPs (0.25 mM), 1.5 µl MgCl2, 1.25 µl forward primer (4 mM), 1.25 µl reverse primer (4 mM), 0.25 µl BSA (20 mg/mL) and 0.125 µl Qiagen *Taq* polymerase. The conditions for the reaction were as follows: a 10’ initial denaturation step at 95 °C was followed by 1’ at 95 °C, 1’ at 55 °C (Fw3_Hali14 and Rv2_Hali14_mod) or 58 °C (HR1_hglT_F and HR2_hglT_R), 2’ at 72 °C, these steps were repeated 30 times, and were followed by a final extension step of 10’ at 72 °C.

The following genes of interest (GOIs) were selected from the *Richelia intracellularis* HH01 genome (taxon ID 2579778779) and their sequence obtained by using The Integrated Microbial Genomes & Microbiomes system (hereafter IMG/M)^80^: RINTHH_5560, RINTHH_5570, RINTHH_17770 (Ri70) and RINTHH_20790 (Ri84-long). Because RINTHH_5560 and RINTHH_5570 were very closely located, a construct containing both genes and their intergenic region (42bp) (RINTHH_5560_5570) was also included. Additionally, *R. intracellularis* HH01 genome was downloaded and annotated using RAST^81^.

From this annotation an open reading frame (ORF) annotated as a glycosyltransferase that was highly similar to RINTHH_20790 (Ri84), albeit shorter (39 bp at the 5’) was selected. All the aforementioned genes were synthesized by Baseclear BV (Leiden, the Netherlands) under the control of P*_glnA_*, a constitutive promoter active in heterocyte and vegetative cells^82^.

The streptomycin/spectinomycin resistance cassette *aadA*^83^, used as selective marker, was also synthesized by Baseclear B.V. and inserted into pBluescript II KS (+). All constructs were flanked by the appropriate GC-adaptors in order to make them compatible for cloning using the CYANO-VECTOR approach^84^. The genes of interest (GOIs) and the resistance cassette were inserted in pAM5404 (Supplementary Fig. 13), an RSF1010-based plasmid containing a mutation in *mobA*Y25F to improve cloning efficiency and an additional RK2-*bom* site to increase its mobilization efficiency^83^, as described in ref ^84^. In brief, plasmid and inserts were digested using *ZraI I* (New England Biolabs) according to the manufacturer’s instructions. Fragments containing the GOIs were purified using a PCR purification kit (QIAquick PCR Purification kit, Qiagen, Hilden, Germany). To remove potential undigested plasmid containing *aadA* in the assembly reaction, the restriction digest of pRP002 was run on a 1% agarose gel, and the appropriate fragment (1279 bp) was cut and purified using QIAquick gel extraction kit (Qiagen, Hilden, Germany).

Final constructs were assembled using NEBuilder® HiFi DNA Assembly Cloning Kit (New England Biolabs) according to the manufacturer’s instructions, transformed into NEB® 5-alpha chemically competent *E. coli* cells and plated on LB agar plates containing spectinomycin and streptomycin. The resulting plasmids were extracted using QIAprep Spin Miniprep Kit (Qiagen, Hilden, Germany) and the presence of the correct insert was confirmed by Sanger sequencing (Baseclear B.V, Leiden, The Netherlands) using the primers PaadA_out, PglnA_F and GOI_R (Supplementary Table 12). In order to make them suitable for tri-parental mating, plasmids (hereafter referred to as cargo plasmids, Supplementary Fig. 14, Supplementary Table 13) were then subcloned in TOP10 cells (Invitrogen, Carlsbad, CA, USA).

### Recombinant plasmid transfer via triparental mating

Genetic modification by triparental mating of *Anabaena* sp. PCC 7120 Δ*hglT* was carried out as described in refs ^85,86^ with a few modifications. As helper strain, we used an *E.coli* strain containing the helper plasmid pRL623 and the conjugal plasmid pRL443^87^, hereafter referred to as helper strain. This strain was generated by biparental mating (see below) of an *E.coli* DH5alphaMCR strain containing pRL623 and a DH5alpha strain containing pRL443. *E.coli* cultures of the helper strain and TOP10 cells strains carrying the cargo plasmids were grown overnight at 37 °C, shaking at 225 rpm (Innova 43, New Brunswick Scientific), diluted 1:20 in LB without antibiotics and then grown for 2.5h at 37 °C shaking at 200 rpm in an Erlenmeyer flask of volume 10 times larger than the volume of the culture. Cells were then harvested by centrifugation for 5 min at 2500 rpm and resuspended gently in 1 mL LB per every 10 mL of initial culture. Cargo and helper plasmid were then mixed in a 1:1 ratio in a total volume of 2 mL, harvested by centrifugation for 5 min at 2500 rpm, resuspended in 100 μl LB and incubated at 30 °C for 1 h without agitation. For each mating we harvested 1.8-3.5 mL of an *Anabaena* sp. PCC 7120 *ΔhglT* (OD_730_ 0.25-0.5) culture grown in BG11 media plus kanamycin by centrifugation for 5 min at 2500 rpm and resuspended it in 100 μl BG11. *Anabaena* cell concentrates were then mixed with the biparental mating of the *E.coli* strains containing the cargo and helper plasmids, harvested by centrifugation and resuspended in 30 μl BG11. Upon resuspension, cells were plated onto a Supor® Membrane Disc Filter (Pall Laboratories) on BG11 agar plates supplemented with 5% LB. The filters were allowed to dry before incubation at 30°C under light dimmed by covering the plates with a tissue paper. After approximately 24h, the filters were washed with BG11 media (Supplementary Table 2) and the resuspended mating’s were plated onto BG11 agar plates supplemented with kanamycin, streptomycin and spectinomycin. Upon appearance of *Anabaena* mutant colonies, single colonies were first replated in BG11 agar plates supplemented with antibiotics and then transferred to liquid BG11 media also with appropriate antibiotics. See Supplementary Table 13 for a summary of all the resulting *Anabaena* strains.

To check that the desired plasmids had been acquired by the host strain, plasmids were extracted using QIAprep Spin Miniprep Kit (Qiagen, Hilden, Germany) following the manufacturer’s instructions with minor modifications, namely: cells from liquid cultures were harvested by centrifugation, resuspended in 500 µl P1 and disrupted as previously described (see Materials and Methods). Upon centrifugation the supernatant was processed as described in the manufacturer’s protocol. Then the GOI was amplified via 50 µl polymerase chain reaction (PCR) containing, 21 µl PCR water, 10 µl Q solution, 5 µl Qiagen Buffer, 4 µl dNTPs (0.25 mM), 3 µl MgCl_2_, 2.5 µl PaadA_out (0.2 mM), 2.5 µl GOI_R (0.2 mM), 0.5 µl BSA (20 mg/mL), 0.25 µl Qiagen *Taq* polymerase and 1 µl plasmid DNA. The conditions for the PCR reaction were as follows: a 10’ initial denaturation step at 95 °C was followed by 10 cycles of 1’ at 95 °C, 1’ at 68.4 °C, 2’ at 72 °C, 20 cycles of 1’ at 95 °C, 1’ at 67 °C, 2’ at 72 °C followed by a final extension step of 10’ at 72°C.

To verify the sequence of the inserted GOIs in the mutant strains, the PCR products of five identical reactions for each strain were pooled and analyzed on a 1% agarose gel. The resulting bands were then purified and concentrated using QIAquick gel extraction kit (Qiagen, Hilden, Germany). The sequences of the concentrated PCR products were confirmed via Sanger sequencing (Baseclear B.V, Leiden, The Netherlands) using the primers PaadA_out, PglnA_F and GOI_R (Supplementary Table 12).

### Lipid extraction and analysis

Extraction of intact polar lipids (IPL) from freeze dried biomass was carried out using a modified Bligh Dyer (BD) extraction as described in ref. ^88^. A known amount of deuterated diacylglyceryltrimethylhomoserine (DGTS D-9, Avanti® Polar Lipids, USA) dissolved in dichloromethane (DCM): methanol (MeOH) (1:9, v:v) was added to the extracts as internal standard and then filtered through a true regenerated cellulose 4 mm syringe filter (0.4 µM, BGB, USA). Filtered extracts were analyzed on an Agilent 1290 Infinity I ultra-high performance liquid chromatographer (UHPLC) with a thermostatted auto-injector, coupled to a Q Exactive Orbitrap MS with an Ion Max source and heated electrospray ionization probe (HESI; ThermoFisher Scientific, Waltham, MA) according to ref. ^89^ (modified from ref ^90^). Briefly, chromatographic separation was achieved with an Acquity BEH C18 column (2.1 × 150 mm, 1.7 µm, Waters), with A) MeOH:H_2_O:formic acid: (14.8M) NH_3_aq (85:15: 0.12:0.04 [v:v]) and B) IPA:MeOH:formic acid:(14.8 M) NH_3_aq (50:50:0.12:0.04 [v:v]) at a flow rate of 0.2 mL min^−1^. Compounds were eluted with 5% B for 3 min, followed by a linear gradient to 40% B at 12 min ending at 100% B at 50 min. Lipids were detected using positive ion monitoring of *m/z* 350–2000 (resolution 70,000 ppm at *m/z* 200), followed by data dependent MS^2^ (isolation window 1 m/z; resolution 17,500 ppm at *m/z* 200) of the 10 most abundant ions. HGs were identified using a targeted approach, each sample was screened using the combined molecular mass of the protonated ([M+H]^+^), ammoniated ([M+NH_4_]^+^) and sodiated ([M+Na]^+^) adducts and were identified by comparison with published MS^2^ spectra (known HGs) and based on theoretical fragmentation for the novel HGs and AGs (extrapolated from the fragmentation of closely related HGs) (Supplementary Table 8).

In order to screen for aglycone-like components in the two LEGE strains, their BD extracts were also analyzed by GC-MS after methylation and silyilation. Methylation was carried out using diazomethane (CH_2_N_2_), then extracts were dried under N_2_, cleaned over a small silica gel column (pore size 60 Å, 0.063-0.2mm, 70-230 mesh, Merck), eluted with 3 times the column volume of ethyl acetate (EtAc) and dried under N_2_. Silylation was caried out by dissolving the methylated extract in and pyridine (10 µL) and allowed to react with N,O-Bis(trimethylsilyl)trifluoroacetamide (BSTFA, Regis Technologies Inc., IL, USA) at 60 °C for 40 min. Samples were then diluted in EtAc and analyzed using a gas chromatographer (Agilent 7990B GC) coupled to a mass spectrometer (Agilent 5977A MSD; GC-MS) equipped with a fused silica capillary column (Agilent CP Sil-5, 25 m x 0.32 mm x 0.12 µm). The temperature program was as follows: start at 70 °C, increased to 130 °C at 20 °C min^−1^, increased to 320 °C at 4 °C min^−1^, held at 320 °C for 25 min. Flow was held constant at 2 mL min^−1^. For further analysis, some extracts were de-silylated by washing 5 times in DCM (500 µL) and drying under N_2_, and then re-silylated by the method above, but with addition of homemade deuterated BSTFA (5 µL) before re-analysis.

### Visualization of lipid data on phylogenetic trees

The relative abundances of each HG in relation to the sum of all HGs produced by each strain were plotted as a heatmap on the phylogenetic trees with iTOL. Data from the literature represented with symbols was converted to percentages as follows; “+++”, 90%; “++”, 40%, “+”, 15%; “tr.”, 1%.

