## Supplementary Information for "Emergence and evolution of heterocyte glycolipid biosynthesis enabled specialized nitrogen fixation in cyanobacteria"

Includes

Supplementary Results and Discussion,

Supplementary Figs. 1-23,

and 42 Supplementary References.

Supplementary Tables 1-20 and Supplementary Data 1-4 are supplied as additional files.

### **Supplementary Results and Discussion**

##### **Genomic prediction of heterocyte glycolipid (HG) biosynthesis in *Cyanobacteriia***

###### **Homology search of HG biosynthesis genes**

The HG gene cluster on the genome of the model cyanobacterium for heterocyte formation *Anabaena* sp. PCC 7120 was initially referred to as an ‘expressed island’, a group of physically clustered genes encompassing multiple transcription units that were regulated in a coordinated manner^1^. However, since it included known ‘*hgl*’ genes, it was later renamed to ‘*hgl* island’^2^. Here, we use the latter name.

The *hgl* island has traits of iterative type I and type II polyketide synthases (PKSs), containing large enzymes consisting of several functional domains (for example ketoacyl synthase, acyl transferase, and acyl carrier protein (ACP) domains on *hglE_A_*^2^) as well as monofunctional enzymes (for example glycosyltransferase *hglT*^3,4^ and ketoacyl synthase *hglD*^2^) (Fig. 1b). Other PKSs in cyanobacteria are responsible for the production of a wide variety of secondary metabolites with diverse functions, such as cytotoxins (e.g. cylindrospermopsin, lyngbyatoxin), antimycotics (e.g. toyocamycin, tubercidin), and photoprotectors (e.g. shinorin), and some even have anticancer properties (e.g. dolastatin, apratoxin)^5,6^.

We hypothesized that genomic colocalization of homologs of these genes in cyanobacterial genomes implies similar function in HG biosynthesis to the *Anabaena* sp. PCC 7120 *hgl* island, discerning them from PKSs that share protein domains but produce other products. Using a subset of 143 genomes of cyanobacteria with known morphological traits including their capacity for heterocyte formation^7^, we confirmed that most heterocytous cyanobacteria (26 out of 28 in the subset) encode an island with homologous hits to at least 13 genes of the queried *Anabaena* sp. PCC 7120 *hgl* island (allowing for at most three open reading frames (ORFs) in between hits and multiple non-overlapping hits per ORF, see Online Methods; Supplementary Fig. 2). The two exceptions are *Raphidiopsis brookii* D9 (homologs of 7 different genes) and *Calothrix parietina* PCC 6303 (homologs of 8 different genes). *Raphidiopsis brookii* D9 has heterocytous ancestors and is therefore characterized as heterocytous cyanobacterium in the subset^7^, but has lost the ability to fix nitrogen and is incapable of heterocyte formation^8^. The absence of a complete *hgl* island in its genome therefore likely reflects gene loss or genomic rearrangements due to loss of function. The *hgl* island of *Calothrix parietina* PCC 6303 is interrupted by several CRISPR-associated proteins flanked by two CRISPR arrays, and its complete *hgl* island (homologs of 14 different genes; Supplementary Table 6) is therefore not detected by our genomic island search, which only allows for at most 3 ORFs in between hits.

In contrast, 109 of 115 non-heterocytous cyanobacteria in the subset have at most homologs of 3 or less of the queried genes clustered on their genome (Supplementary Fig. 2). Only two of the 115 genomes have an island with homologs of more than 5 of the queried genes: the unicellular *Gloeobacter violaceus* PCC 7421 (homologs of 7 different genes), whose homologs of *hgl* island genes with unknown function have been noticed before^2^, and the unicellular *Gloeocapsa* sp. PCC 7428 (homologs of 10 different genes). We conclude that the ability to make heterocytes can be identified with high certainty from the genome sequence based on the presence of the here-defined *hgl* island. Based on this benchmark, we investigated islands with homologs of at least 7 of the 19 queried genes as potentially involved in the biosynthesis of HGs, which we called *hgl* islands from hereon. This relatively permissive definition allowed us to detect PKSs that are likely involved in the biosynthesis of unknown HG-related products and reconstruct the evolutionary history of HG biosynthesis.

Next, we explored the *hgl* islands in our full set of cyanobacterial genomes (Supplementary Data 2). These included 3,579 genomes and plasmids from the PATRIC genome database (Supplementary Data 1) and 14 genomes that were newly sequenced here (see Online Methods). The 14 sequenced genomes were from cultures of which we analyzed the lipids (see below) but that did not have their genome sequenced yet. Of the 3,593 genomes, 3,339 contained at least one hit to a queried protein, and 500 contained one or more *hgl* island(s) as defined here (i.e. a genomic cluster of homologs of at least 7 of the queried HG biosynthesis genes) (Supplementary Data 2).

For further confirmation that the presence of an *hgl* island is associated with heterocyte formation, we screened the genomes for the presence of nitrogenase (*nif*) genes involved in the process of nitrogen fixation, which are also encoded by a genomic island in *Anabaena* sp. PCC 7120 (Supplementary Table 5; ref. ^9^). This screen revealed that most genomes with an *hgl* island also encode a *nif* island (here defined as a genomic cluster of at least 5 *nif* genes; Supplementary Fig. 3), and are thus capable of nitrogen fixation.

##### ***Hgl* and *hgl-like* islands are present throughout *Cyanobacteriia***

###### **Identification of monophyletic heterocytous clade within Cyanobacteriia**

We reconstructed the cyanobacterial species phylogeny based on a concatenated alignment of 24 conserved vertically transferred genes (selected from ref. ^10^) including one representative genome per species based on sequence similarity (average nucleotide identity >= 95%) and using *Vampirovibrionia* and *Sericytochromatia* genomes as outgroup for rooting^11,12^ (see Online Methods). Plotting the *hgl* islands on this phylogeny (Fig. 2a) revealed a large monophyletic group of cyanobacteria in which almost all genomes contain an *hgl* island composed of homologs of at least 10 of the queried genes, and which includes the genomes of known heterocyte-producing cyanobacteria mentioned earlier. We call this clade the ‘heterocytous clade’. The clade contains uncultured genomes with unknown morphology. The most closely related sister clade to the heterocytous clade contains a single uncultured cyanobacterium from Antarctic soil (the metagenome-assembled genome *Nostocaceae* cyanobacterium MGR_bin409) that does not encode an *hgl* nor *nif* island and we thus assume is non-heterocytous. The most closely related sister clade to the heterocytous clade and *Nostocaceae* cyanobacterium MGR_bin409 contains genomes with known unicellular and baeocytous morphology and contains mainly non-diazotrophic cyanobacteria based on the absence of a *nif* island (Fig. 2a). Nevertheless, some of the inferred non-heterocytous genomes in this sister clade contain an *hgl* island (Fig. 2a; see Results and Discussion and below for a discussion on *hgl* islands in non-heterocytous cyanobacteria).

###### **Gene conservation on hgl islands with homologs of at least 10 HG biosynthesis genes**

First, we explored the genetic content of the identified islands. The genomic compositional variation within the islands includes gene absences and fusions/fissions, gene duplications and some positional rearrangements (Fig. 2a). For the *hgl* islands containing homologs of at least 10 queried genes, we examined the most common genes present. Homologs of 13 genes were present on ≥88% of these islands (Fig. 2c), and their position on the island was largely conserved (Supplementary Fig. 4). In previous mutagenesis studies, 12 of the 13 genes have been shown to be essential to *Anabaena* sp. PCC 7120 for survival and growth in nitrogen-depleted conditions (Supplementary Table 4; refs. ^2–4,13–18^). They include the epimerase *hgdA* and two genes encoding ATP-binding cassette (ABC) transporter proteins, *hgdC* and *hgdB*, responsible for transport of the HG and its deposition in the cell envelope; the gene encoding ketoacyl synthase, acyl transferase, and ACP domains, *hglE_A_*; the gene *hglF* of unknown function; the gene *hglG* containing ketoreductase and dehydratase domains; the ketoacyl synthase *hglD*; the gene *hglC* containing a chain length factor and acyl transferase domains; the enoyl reductase *hglA*; and the gene *hglB* (also known as *hetM*), containing ACP and thioester reductase domains (Fig. 2c, Fig. 1, Supplementary Fig. 1). Additional studies have shown that although *hglT*, which encodes the glycosyltransferase (GT) that attaches the glucose headgroup to the aglycone (AG), and *all5343*, a gene of unknown function, are required for proper heterocyte development and HG layer deposition, they are not essential for survival and growth in nitrogen-depleted conditions (Supplementary Table 4, refs. ^2–4^). The role of *all5344* in HG biosynthesis has not yet been elucidated with mutagenesis studies. The 13 genes are likely evolutionary conserved on the *hgl* island to preserve function in heterocyte formation. The three genes *all5343*, *all5344*, and *hglF* have yet unknown function but are present in 88%, 90%, and 99% of the *hgl* islands with homologs of at least 10 queried genes, respectively (Fig. 2c), which may further indicate an essential role in HG biosynthesis.

Genes that are often absent from the *hgl* islands may be non-essential for HG biosynthesis, be present somewhere else on the genome (Supplementary Fig. 5), or their catalytic function may be replaced by unrelated genes. For example, homologs of *alr5348,* encoding a protein with SAM-dependent methylase domains, and *asr5350*, encoding a protein of unknown function, are both clade-specific and only present in approximately half of the *hgl* islands containing homologs of at least 10 queried genes (Fig. 2a,c). The 4’-phosphopantetheinyltransferase (PPTase) *hetI* is present on 52% of these *hgl* islands. PPTases are essential enzymes required for the biosynthesis of a wide variety of compounds such as fatty acids, polyketides, and non-ribosomal peptides^19^, and all genomes with diverse *hgl* islands encode *hetI* homologs, even if not always in the vicinity of the other HG biosynthesis genes (Supplementary Fig. 5). Many members of subclade 6 (see Fig. 2a) lack homologs of GTs and of proteins involved in HG deposition (*hgdACB*) on their *hgl* island. The GT *all5342* is rarely present on *hgl* islands (Fig. 2c), and *asr5349* is restricted to *Anabaena* sp. PCC 7120, reflecting clade-specific genes in our *hgl* island query.

Even though our protein queries represented individual open reading frames (ORFs), homologous sequences of *hglD* and *hglC*, and of *hglE_A_* and *hglF*, were often encoded by the same ORF (90% and 48% respectively, Fig. 2a,c), which was previously reported as anecdotical^2^. The two independent ORFs of *hglE_A_* and *hglF* in *Anabaena* sp. PCC 7120 thus likely represent a rare gene fission from a single ancestral ORF containing the domains encoded by both genes.

**The role of *hgl*-like island in heterocytous cyanobacteria is unclear**

Our analysis thus reveals that HG biosynthesis genes are evolutionary conserved as a cluster closely resembling the *Anabaena* sp. PCC 7120 *hgl* island—containing homologs of at least 10 of the queried genes—within a monophyletic heterocytous clade in *Cyanobacteriia*. These *hgl* islands are likely involved in HG biosynthesis and heterocyte formation. Moreover, various heterocytous cyanobacteria also encode an ‘*hgl*-like’ island with a highly conserved gene composition containing homologs of only 7 HG biosynthesis genes (the transporters *hgdCB* and PKS *hglE_A_FGCA*) encoded by 5 ORFs in addition to a more extended *hgl* island (see Results and Discussion and Fig. 2). Given that many heterocytous strains do not possess an *hgl*-like island (Fig. 2) we suspect that it does not always have an essential function. However, its preservation during over 2 billion years of evolution in *Cyanobacteriia* suggests an important function in those strains where it is present.

Based on its prevalence in heterocytous cyanobacteria we hypothesized that the *hgl*-like island might produce and export compounds structurally similar to HGs which may also be involved in nitrogen fixation. However, genomic analysis revealed that the *hgl*-like island is likely not involved in nitrogen fixation. Certain strains such as *Raphidiopsis curvata* NIES-932, *Cylindrospermopsis raciborskii* CENA303, and the earlier discussed *Raphidiopsis brookii* D9 do not possess a *nif* island (Supplementary Data 2) and are thus incapable of nitrogen fixation even though they evolutionary branch from within the heterocytous clade (see also refs. ^8,20^). These non-diazotrophic strains possess a single *hgl* island that is more similar in sequence to the *hgl*-like islands of heterocytous cyanobacteria than to the more extended *hgl* island in those strains (see section ‘Evolutionary reconstruction of HG biosynthesis within *Cyanobacteriia*’ below). Thus, even though they have lost their extended *hgl* island likely in relation to the loss of heterocyte formation, they have kept their *hgl*-like island, suggesting that it conveys a function independent of nitrogen fixation.

Since akinetes are thought to be the precursors of heterocytes^21^ and are shown to be surrounded by the same layer of glycolipids and polysaccharides in some cultures^22,23^ we also speculated a potential role of the *hgl*-like island in akinete formation. However, mutational studies have shown that the HGs found in akinetes and heterocytes are synthesized by the same set of genes^17^. In addition, our data show that not all strains that form akinetes possess an *hgl*-like island (e.g. *Anabaena variabilis* ATCC 29413). Therefore, we deemed this hypothesis unlikely. Thus, the potential role of the *hgl*-like island within heterocytous cyanobacteria is still unclear.

##### **Heterocytous cyanobacteria produce a wide diversity of new HGs**

###### **Identification of new HGs in heterocytous cyanobacterial cultures**

We analyzed HGs in the biomass of 26 heterocytous cyanobacterial cultures whose HG profiles were previously published (Supplementary Table 10). The cultures were evenly selected throughout the heterocytous cyanobacterial phylogeny and included representatives from all subclades in Fig. 2a with the exception of subclade 1.

HG biosynthesis involves several rounds of condensation, reduction, and dehydration of acyl-CoA precursors (Supplementary Fig. 1), and different combinations of these steps result in a large theoretically possible number of structures. We screened the lipid extracts of the cultures for 88 theoretically possible HG structures including 19 HGs previously described in the literature, using ultra-high-performance liquid chromatography coupled with multistage high-resolution mass spectrometry (UHPLC-HRMS*^n^*). A wide range of HGs were detected (Supplementary Table 11), including the 19 HGs previously described in the literature, which had been identified from their accurate masses and fragmentation using other techniques such as high performance liquid chromatography coupled to electrospray ionization tandem mass spectrometry (HPLC–ESI-MS^2^, refs. ^24,25^) or ultra-high performance liquid chromatography coupled to an ultra-high-resolution quadrupole time-of-flight (QToF) mass spectrometer^26^. The previously reported HGs included thirteen hexose HGs with either 26, 28, 30 or 32 n-alkyl chains and with either two (keto-ol, diol, or diketone) or three (triol and keto-diol) oxygen moieties on their alkyl chain^24–28^. Additionally, we detected several HGs with alternative sugar moieties that had been previously described in the literature, including four pentose HGs (pentose HG_26_ diol, pentose HG_30_ diol, pentose HG_30_ triol, and pentose HG_32_ triol^26,29,30^, plus the previously described methyl-hexose HG_28_ (ref. ^31^), and deoxy-hexose HG_26_ (ref. ^26^). We note that while ‘pentose’ refers to a C_5_ sugar, the pentoses detected in the cultures in this study may be in the pentopyranose form (see also section ‘Potential origin of HGs from ancient 1,3-diols’) instead of the furanose detected in *Richelia* species^32^ (see below).

In addition, 30 HGs were detected that, to the best of our knowledge, have not previously been described in the literature (Supplementary Tables 10 and 11). 22 of these novel HGs had a known sugar moiety and either a 26, 28, 30 or 32 carbon alkyl chain with alcohol or keto moieties—as described above for the known HGs, but in combinations not previously described. These were thus straightforward to identify from their accurate masses, their fragmentation patterns, and their elution order (Supplementary table 9). 6 of these were hexose HGs, 6 were pentose HGs, 8 were methyl-hexose HGs and 1 was a deoxy-hexose HG. Finally, we found nine novel HGs with more unusual structures (Supplementary Table 11, Supplementary Fig. 6), which we discuss below.

One of the unusual novel HGs gave rise to a dominant ion at mass-to-charge ratio (*m/z*) 576.483, assigned as C_32_H_66_O_7_N. Under MS^2^ fragmentation (Supplementary Fig. 7a), this ion underwent neutral loss of 179.080 Da (C_6_H_13_O_5_N), indicative of a hexose and NH_3_ loss from a [M+NH_4_]^+^ ion, to produce a fragment ion at *m/z* 397.403 (C_26_H_53_O_2_). This fragment ion also underwent a subsequent loss of H_2_O to produce an ion at *m/z* 379.393 (C_26_H_51_O), followed by another loss of H_2_O resulting in an ion at *m/z* 361.382 (C_26_H_49_). The combination of these neutral losses provides evidence that the parent ion at *m/z* 576.483 is the [M+NH_4_]^+^ of a hexose HG_26_ keto. To the best of our knowledge, HGs with a single oxygen moiety on the alkyl chain have not previously been reported. Similarly, a second novel HG gave rise to an ion at *m/z* 578.499 (C_32_H_68_O_7_N), showed a similar fragmentation in MS^2^ (Supplementary Fig. 7b) as the previously described compound, but eluted several minutes earlier. Hence, this component was assigned as a hexose HG_26_ ol (see also section ‘Potential origin of HGs from ancient 1,3-diols’ below). One or both of these two components were detected in low levels (<2% of total HG sum) in 6 cyanobacterial cultures.

Three structurally similar components gave rise to ions at *m/z* 617.462, *m/z* 645.494 and *m/z* 673.525. All three contained a hexose sugar moiety, as evidenced by the primary neutral loss of 162.053 Da (C_6_H_10_O_5_) in MS^2^ (Supplementary Fig. 8). The neutral loss of the sugar gave rise to primary fragments ions at *m/z* 455.409 (C_28_H_54_O_4_), *m/z* 483.441 (C_30_H_58_O_4_), and *m/z* 511.472 (C_32_H_62_O_4_), respectively. These all underwent four sequential losses of H_2_O. Based on their accurate masses and their fragmentation patterns, these three components were assigned as hexose HG_28_ diketo-ol, hexose HG_30_ diketo-ol, and hexose HG_32_ diketo-ol, respectively.

Another unusual HG gave rise to an assumed [M+H]^+^ ion at *m/z* 591.484 (C_33_H_67_O_8_) and was detected in eight cultures. This HG contained a hexose sugar moiety (as revealed from the 162.053 Da neutral loss from the parent ion) in MS^2^ (Supplementary Fig. 9a). This neutral loss led to the formation of a fragment ion at *m/z* 429.430, equivalent to C_27_H_57_O_3_. This suggests that this HG contains a 27-carbon alkyl chain, in contrast to all previously described HGs that have even-numbered alkyl chains. From the *m/z* 429.430 fragment ion there were consecutive losses of H_2_O giving rise to ions at *m/z* 411.418 (C_27_H_55_O_2_), *m/z* 393.408 (C_27_H_53_O) and *m/z* 375.397 (C_27_H_51_), a fragmentation pattern similar to that regularly described for HG diols^24^ . Hence, this novel HG was assigned as a hexose HG_27_ diol. Similarly, another novel ‘HG-like’ component gave rise to a [M+H]^+^ at *m/z* 617.499 (C_35_H_69_O_8_), which underwent loss of a hexose moiety in MS^2^ (Supplementary Fig. 9b), resulting in fragment ions with a 29 carbon chain. This novel component, which was detected in two cultures (*Tolypothrix tenuis* PCC 7101 and *Fortiea contorta* PCC 7126), was assigned as a hexose HG_29_ keto-ol.

Another unusual novel HG had an [M+H]^+^ at *m/z* 633.530 (C_35_H_69_O9) and was detected in just one culture (*Scytonema* sp. PCC 10023). In MS^2^ (Supplementary Fig. 10a) this ion gave rise to a fragment ion at *m/z* 501.488 (C_30_H_61_O_5_), due to a neutral loss of 132.043 Da (C_5_H_8_O_4_), characteristic of a pentose sugar moiety^32^. The fragment ion at *m/z* 501.488 unusually underwent 5 subsequent losses of H_2_O, resulting in fragment ions at *m/z* 483.440 (C_30_H_59_O_4_), *m/z* 465.429 (C_30_H_57_O_3_), *m/z* 447.419 (C_30_H_55_O_2_), *m/z* 429.409 (C_30_H_53_O), and finally *m/z* 411.399 (C_30_H_51_), indicative of a component with four oxygen moieties on the alkyl chain. Hence this novel HG was assigned as a pentose HG_30_ keto-triol.

Finally, a novel HG-like compound was detected in three cultures (*Chlorogloeopsis fritschii* PCC 6912, *Chlorogloeopsis fritschii* CCY 9923, and *Fischerella muscicola* PCC 73103) and gave rise to an apparent [M+H]^+^ ion at *m/z* 613.504. The MS^2^ spectrum arising from fragmentation of this ion (Supplementary Fig. 10b) revealed a fragment pattern characteristic of an HG with a C_30_ keto-ol component, e.g., dominant ions at *m/z* 469.463 (C_30_H_61_O_3_), *m/z* 451.450 (C_30_H_59_O_2_), *m/z* 433.440 (C_30_H_57_O) and *m/z* 415.429 (C_30_H_55_). However, the initial neutral loss normally associated with loss of the sugar moiety was 144.043 Da, equivalent to C_6_H_8_O_4_. Structural identification of the assumed sugar moiety was not possible based on its accurate mass and hence is assigned as an HG-like, C_30_ keto-ol with an unknown headgroup.

##### **HG structure is mostly independent of *hgl* island gene composition**

###### **HG abundances and distribution throughout the heterocytous cyanobacterial phylogeny**

Next, we studied the distribution of the detected HGs across the 26 cultures analyzed. We focused on the three characteristics of the HG structures: headgroup, functional groups on the alkyl chain, and alkyl chain length.

*Headgroups*. In all the cultures analyzed, hexose was the most abundant HG headgroup, followed by pentose (possibly in pentopyranose form), deoxyhexose, and methyl hexose (Supplementary table 10). The capacity to produce HGs with three different sugar headgroups was present in strains scattered across the heterocytous cyanobacterial phylogeny (Fig. 3). In general, HGs with alternative headgroups to hexose appeared in small amounts (<0.1 - 3.8%) and usually only in conjunction with the abundant production (>20%) of that same HG with a hexose headgroup. However, in some cases HGs with headgroups other than hexose were found in relatively high abundances such as pentose HG_26_ diol in *Nostoc* sp. CCY 0012 (14.4%), deoxy-hexose HG_26_ diol in *Anabaena* sp. CCY 0017 (20.1%) and *Nodularia chucula* CCY 0103 (7.9%), methyl-hexose HG_28_ triol in *Calothrix* sp. CCY 0202 (17.2%), and methyl-hexose HG_30_ triol in *Tolypothrix tenuis* PCC 7110 (16.8%). In addition, pentose HG_30_ keto-triol was the only HG with a pentose headgroup for which a hexose equivalent has not been detected, neither here nor in previous studies (Supplementary Tables 10 and 11).

*Functional groups*. Overall, most cultures analyzed synthesized most of their HGs (>93%) with either 2 (diol and keto-ol) or 3 (keto-diol and triol) functional groups on the alkyl chain besides their sugar moiety. Only one of the analyzed cultures, *Scytonema* sp. PCC10023, produced an HG with four functional groups (pentose HG_32_ keto-triol, 2.7%). Cultures of strains found in subclades 2-4 (Fig. 3, Supplementary Fig. 6) produced mostly HGs with three functional groups, usually keto-diols and triols, but also in some cases diketo-ols. Cultures of strains in subclades 5-9 (Supplementary Fig. 6) mostly produced HGs with only two functional groups, such as diols and keto-ols, but they also synthesized diketones and HGs with only one keto or alcohol group. Even though we found clade-specific dominance of the number of functional groups, HGs with two and three functional groups were also found in low abundance throughout subclades 2-4, and subclades 5-9, respectively. Strains capable of producing diketones (with or without an additional alcohol group) were also found throughout the phylogeny.

*Chain length*. Although most cultures analyzed produced most (>91%) of their total HGs with only one specific alkyl chain length, they were also capable of producing HGs with different alkyl chain lengths. One exception to the dominance of a single chain length was *Tolypothrix tenuis* PCC 7101, which produced hexose HG_28_ and hexose HG_30_ in similar proportions (42% and 57%, respectively). Notably, cultures of strains from subclades 2-4 were capable of producing HG_28,30,32_, whilst cultures of strains from subclades 5-9 usually produced HG_26,28_ (Supplementary Fig. 6). An exception was again *Tolypothrix tenuis* PCC 7101, a strain found in subclade 6 (Supplementary Fig. 6), which besides hexose HG_28_ also produces hexose HG_30_. Although cultures from closely related strains usually produced HGs of the same length, this was not always the case. For example, *Nostoc* sp. CCY 9925 and *Nostoc* sp. CCY 9926 (both from subclade 9) synthesized hexose HG_28_ and hexose HG_26_, respectively. Additionally, odd-chain HGs were only found in cultures of strains from subclades 5-9 (Fig. 3, Supplementary Fig. 6, Supplementary Table 11).

###### **HglT is absent from pentose HG producing symbiotic cyanobacteria**

As mentioned above, the protein product of *hglT* is responsible for attaching the sugar headgroup to the AG in the last step of HG biosynthesis by *Anabaena* sp. PCC 7120. However, homologs of this gene are not always present on the *hgl* islands (Fig. 2c).

The alignment scores of hits obtained during our homology search of the *hglT* of *Anabaena* sp. PCC 7120 show a bimodal distribution, where hits with a bit-score >350 are mostly located on the *hgl* island (Supplementary Fig. 11, orange bars). All *hglT* hits that are on an *hgl* island have a high bit-score, suggesting that the bit-score reflects conserved enzymatic functionality. In contrast, most genomes in which the best hit for *hglT* has a bit-score <200 do not contain an *hgl* island (Supplementary Fig. 11, purple bars). Hence, hits with low-scoring alignments may be indicative of distant homologs with a potentially divergent function unrelated to HG biosynthesis. Incidentally, there are some genomes with high-scoring *hglT* hits that do not contain the homolog on their island (Supplementary Fig. 12, cyan bars with bit-score >350) or that do not contain an *hgl* island (Supplementary Fig. 11, purple bars with bit-score >350). This could be due to (based on manual curation) fragmented assemblies of genomes (in which case the *hgl* island is absent or located close to a contiguous sequence (contig) edge), or disruption of the *hgl* island for example by transposases or by CRISPR arrays (e.g *Calothrix* *parietina* PCC 6303 discussed above, Supplementary Table 6). We assume that these genomes encode conventional *hglT* activity even if its homolog is not found within an *hgl* island based on our automated searches. However, there are some cyanobacteria that do contain an *hgl* island but lack a high-scoring *hglT* hit anywhere on their genome (Supplementary Fig. 11, cyan bars with bit-score < 200). For example, high-scoring *hglT* hits are absent from *Richelia euintracellularis* HM01*,* *Richelia* *intracellularis* RC01 and *Richelia* *rhizosoleniae* SC01 genomes. These three cyanobacteria form symbiotic relationships with marine diatoms, and previous studies have shown that *Richelia euintracellularis* HM01 and *Richelia* *intracellularis* RC01 exclusively produce HGs with pentose headgroups in the furanose form^29,30^, in contrast to other analyzed cultures which predominantly produce hexose HGs (Fig. 3). The HG profile of *Richelia* *rhizosoleniae* SC01—a symbiont of the marine diatom *Chaetoceros compressus*—is not known, but since it also does not encode a high-scoring *hglT* hit on its genome, we predict that it also produces pentose HGs like the other two *Richelia* species.

We hypothesized that a gene other than a high-scoring *hglT* homolog with substrate specificity for pentose instead of hexose (hereafter *hglT2*) is responsible for the addition of the sugar headgroup to the AG in these species. To find the gene responsible in *Richelia euintracellularis* HH01, an endosymbiont of *Hemiaulus hauckii*, we used two approaches: (a) sequence homology to *hglT*, and (b) functional annotation as a GT and colocalization on the *hgl* island. Each approach returned two candidate genes in *Richelia euintracellularis* HH01: (a) *RINTHH_17770* and *RINTHH_20790*, and (b) *RINTHH_5560* and *RINTHH_5570.* The candidate genes were heterologously expressed in an *Anabaena* sp. PCC 7120 strain that could only produce AGs and no HGs due to the deletion of *hglT* (∆*hglT*)^4^ (see Online Methods, Supplementary Figs. 12-14 and Supplementary Tables 12-16). Since *RINTHH_5560* and *RINTHH_5570* were only separated by 42 base pairs (bp), we expressed them together including their intergenic region. *Anabaena* sp. PCC 7120 *wild-type* (*wt*) was reported to only produce hexose HGs^33^, while here for the first time we also detected the presence of a very low amount of pentose HGs in the *wt* phenotype (0.28%; Supplementary Table 16), although possibly in the pentopyranose form instead of the furanose found in *Richelia* species. Hence, upon expression of *hglT2* in the HG-deficient strain*,* we expected mainly production of HGs with a pentose headgroup.

However, although the control strain expressing the deleted *hglT* on a plasmid (ARP001) did recover the original *wt* phenotype, producing hexose HGs (97%) and small amounts of pentose HG_26_ (0.19%), none of our candidate genes led to the generation of pentose HGs (Supplementary Table 16). Low amounts of hexose HG_26_ diol (< 0.17%) were still detected in strain ∆*hglT* and in all mutants generated in this study (Supplementary Table 16), potentially caused by spontaneous reaction or by the unspecific activity of another GT. There are several possible explanations for why no pentose HGs were detected in any of the mutants: (1) none of the candidate genes is *hglT2*; (2) we did not express the functional sequence, as for example the ORF of *RINTHH_20790* differs when using different gene calling methods (See Online Methods); (3) *Richelia euintracellularis* HH01 proteins are not functional in *Anabaena* sp. PCC 7120 due to physiological differences potentially affecting their folding and catalytic activity (e.g. intracellular pH, redox potential, absence of required chaperones, cofactors or ligands, etc.); (4) limited availability of free pentose sugars in *Anabaena* sp. PCC 7120, and (5) *hglT2* acts as a molecular caliper and can only catalyze the reaction when the substrate AG has the correct chain length (note that *Richelia euintracellularis* HH01 produces HG_30,32_, whilst *Anabaena* sp. PCC 7120 synthesizes HG_26,28_). Given that strains where *hglT* is expressed (*wt* and ARP001) do synthesize small amounts of pentose HGs, and that *Anabaena* sp. PCC 7120 is known to have abundant pentoses^34^ we consider explanation (4) unlikely. To test explanation (5), one could express the GT of a known hexose HG_30_ or hexose HG_32_ producer (such as *Chlorogloeopsis fritschii* PCC 6912) in the ∆*hglT* background strain. If no hexose HGs are produced, this suggests that the chain length of the AG is a determining factor in the GT activity, but it would not rule out possible explanations (1), (2) or (3).

##### **HG structure evolved convergently**

###### **Re-evaluating HGs as biomarkers**

For years, the chemical structure of HGs and their relative abundance were thought to have chemotaxonomical value and were used to distinguish between cyanobacteria of different orders and families^7,14,36^. However, the increasing number of strains analyzed and the recent changes in cyanobacterial taxonomy, combined with our high-resolution lipid analysis of 26 heterocytous cultures, are challenging this approach. For example, large concentrations of hexose HG_26_'s coupled with small amounts of hexose HG_28_'s were associated with the former *Nostocaceae* family^36^ (currently *Nostocaceae* and *Aphanizomenonaceae* families according to the latest taxonomic classification^35^ (Supplementary Table 17)), however, in some members of these families HG_28_'s account for a large proportion of the total HGs (e.g. *Dolichospermum* sp. BIR169 and *Aphanizomenon* sp. TR83, 46% and 12%, respectively)^36^. Another example is the recent identification of a hexose HG_28_ diketone in *Microchaete* sp. PCC 7126 (currently *Fortiea contorta*)^40^ and its proposal as potential biomarker to track the *Microchaete* genus (currently representing members of the *Microarchaete* and *Fortiea* genera)^41^. However, here we show that this diketone is also produced by *Anabaena* and *Nostoc* strains (Supplementary Table 11), hence rendering it unusable as an exclusive taxonomic biomarker for neither the *Fortiea* nor the *Microchaete* genera. Additionally, the evolutionary evidence presented in this study suggests that the HG profiles observed in extant cyanobacteria resulted from convergent evolution (Fig. 3 and Supplementary Figs. 6 and 16), a process where the same trait evolves independently in different species likely triggered by environmental factors, thus further challenging their use as taxonomy biomarkers. For instance, survival under extreme temperatures or high oxygen concentrations might have been only possible in strains capable of producing HGs with ‘long’ alkyl chains which might provide increased structural integrity and impermeability^32,37^.

##### ***Hgl* islands predate heterocyte formation**

###### **Evolutionary reconstruction of HG biosynthesis within Cyanobacteriia**

As mentioned above (see also Results and Discussion), HG biosynthesis genes were evolutionary conserved as a cluster closely resembling the *Anabaena* sp. PCC 7120 *hgl* island—containing homologs of at least 10 of the queried genes—within a monophyletic heterocytous clade in *Cyanobacteriia*. In addition, several distantly related non-heterocytous cyanobacteria also encode *hgl* islands with a more diverged gene composition, and various heterocytous cyanobacteria encode an additional *hgl*-like island with a highly conserved gene composition containing homologs of only 7 HG biosynthesis genes encoded by 5 ORFs (see Results and Discussion and Fig. 2). The broad presence of *hgl* and *hgl*-like islands in heterocytous and non-heterocytous cyanobacteria provides genomic evidence to elucidate the origin of HGs within *Cyanobacteriia*. To reconstruct the evolutionary history of HG biosynthesis, we constructed a phylogeny based on the concatenated alignment of 7 genes (*hgdCB* and *hglE_A_FGCA*) that are shared among most *hgl* and *hgl*-like islands (Fig. 4, Supplementary Fig. 15).

In this phylogeny, the more extended *hgl* islands with homologs of least 10 genes discussed above and more compact *hgl*-like islands each formed distinct groups (Fig. 4, Supplementary Fig. 17, Supplementary Table 18). This shows that a duplication that gave rise to the two types of islands on single genomes of heterocytous cyanobacteria (*hgl* and *hgl*-like islands) predates the Last Heterocytous Cyanobacterial Common Ancestor (LHeCCA). When a root was placed in between these groups (Fig. 4), both had a similar tree topology to that of the phylogeny created using the core vertically-transferred genes (Fig. 2a)—subclades that were monophyletic in the concatenated core gene tree were also largely monophyletic in the concatenated HG biosynthesis gene tree—suggesting primarily vertical inheritance of both island types (Fig. 5). Differences in branching of deeper clades between the two phylogenetic trees may be explained by the shorter alignment length of the concatenated HG biosynthesis gene tree (3,666 amino acids) compared to concatenated core gene tree (6,933 amino acids) or by different evolutionary histories of individual genes. Alternatively, horizontal transfer of the complete island early in heterocytous cyanobacterial radiation can explain the observed discrepancies. We identified one plasmid (pNFSY07) of *Nostoc flagelliforme* CCNUN1, a strain isolated from desert soil of Inner Mongolia^38^, containing a cluster of homologs of 5 queried genes (*hglE_A_FGCA*; see Supplementary Data 1 and 2, genome ID 2038116.9), showing that HG biosynthesis genes occasionally transfer together horizontally in mobile genetic elements.

Some heterocytous cyanobacteria from subclades 4 and 9 contain additional *hgl* islands with a gene composition that differs from the *hgl*-like islands (Fig. 4 and Supplementary Fig. 17). These islands did not cluster with the other *hgl* and *hgl*-like islands from their respective subclade. Instead, they were phylogenetically placed—together with the *hgl* islands of non-heterocytous cyanobacteria—between the two broad heterocytous phylogenetic clusters discussed above (i.e. the more extended *hgl* island and *hgl*-like islands of heterocytous cyanobacteria, whose evolutionary histories were similar to that of the concatenated core gene tree) (Figs. 4 and 5, and Supplementary Fig. 17). Even though the detailed evolutionary history of these *hgl* islands is unclear, we assume that their phylogenetic placement suggests an origin predating LHeCCa.

The phylogenetic placement of *hgl* islands belonging to non-heterocytous cyanobacteria in the concatenated HG biosynthesis gene tree between the two broad heterocytous phylogenetic clusters (Fig. 4 and Supplementary Fig. 17) depicts an ambiguous evolutionary history. The islands of a closely related sister clade of heterocytous cyanobacteria cluster closely together with *hgl*-like islands instead of with the more extended heterocytous *hgl* islands, and the *hgl* islands of, for example, the more distantly related filamentous *Moorea* species are more similar to the more extended heterocytous *hgl* islands than these sister clade islands are (Supplementary Fig. 17). The *Moorea* islands cluster with one of the two additional *hgl* islands found in the heterocytous *Chlorogloeopsis fritschii* PCC 6912 (Supplementary Fig. 17). These results support the presence of an *hgl* island in the last common ancestor of *Cyanobacteriia*, or alternatively multiple horizontal transfer events predating LHeCCA. In both cases, widespread loss explains the sparse distribution of *hgl* islands in *Cyanobacteriia* today.

##### **Potential origin of HGs from ancient 1,3-diols**

###### **Two cultures of non-heterocytous cyanobacteria with an hgl island do not produce HGs**

To further investigate the origin of HG biosynthesis, we screened two strains from outside the heterocytous clade that contained an *hgl* island—*Pleurocapsales* cyanobacterium LEGE 10410 (*hgl* island with homologs of 10 different genes; *hgdCB*, *all5343*, *all5344*, and *hglBACGFE_A_*; Supplementary Data 2) and *Gloeocapsopsis crepidinum* LEGE 06123 (*hgl* island with homologs of 10 different hits; *hgdCB*, *alr5348*, *asr5350*, and *hglE_A_FGCAB*; Supplementary Data 2)—for the presence of HGs and HG-related compounds. *Gloeocapsopsis crepidinum* LEGE 06123 is part of a closely related sister clade of the heterocytous cyanobacteria, and *Pleurocapsales* cyanobacterium LEGE 10410 is much more distantly related (Figs. 2a and 6a). Based on the presence of a *nif* island (Supplementary Table 7) both these non-heterocytous strains are capable of nitrogen fixation. As the *hgl* islands in these genomes likely represent a phylogenetic lineage of the island that predates LHeCCA (see section ‘Evolutionary reconstruction of HG biosynthesis within *Cyanobacteriia*’ above), their biosynthetic products may be ‘remnants’ of molecules that were produced before the emergence of the heterocyte, and thus give insights into the evolutionary acquisition of HGs for heterocyte formation.

The gene compositions of the *hgl* islands of both strains were relatively similar (Fig. 6d). Out of the 13 genes that are commonly conserved on regular *hgl* islands (see section ‘Gene conservation on *hgl* islands with homologs of at least 10 HG biosynthesis genes’ above), homologs of 8 genes were present on the *hgl* islands of both strains (*hgdCB* and *hglE_A_FGCAB*), and *Pleurocapsales* cyanobacterium LEGE 10410 also encoded homologs of *all5343* and *all5344*. Absent from both islands were homologs of *hglT*, *hgdA*, and *hglD*. Whereas *Pleurocapsales* cyanobacterium LEGE 10410 does not contain a high-scoring *hglT* homolog anywhere on its genome (bit-score of best hit: 137; see section ‘*HglT* is absent from pentose HG producing symbiotic cyanobacteria’ above for a discussion of ‘high-scoring’ *hglT* hits), a high-scoring homolog of *hglT* is encoded close to the *hgl* island of *Gloeocapsopsis crepidinum* LEGE 06123 (6 ORFs from the *hglB* homolog, with a bit-sore of 549; Supplementary Data 2).

In culture, daughter cells of both strains remained attached to the parent cell upon division, and were surrounded by a polysaccharide layer, thus appearing as aggregates under the microscope (Supplementary Fig. 18), under all growth conditions. We did not identify HGs in any of the tested culturing conditions, neither after ageing nor in media without nitrogen (Supplementary Fig. 18) which would induce akinete and heterocyte formation (respectively) in heterocytous cyanobacteria. Based on the absence of HGs, we hypothesized that the *hgl* islands of the two strains produce compounds that resemble HGs or AGs but are structurally different, which we call ‘HG analogs’ from hereon. Based on the absence of *hglT*—the GT that attaches the glucose headgroup to the AG in canonical HG biosynthesis (Supplementary Fig. 1)—from the *hgl* islands of both strains, our initial targets for potential HG analogs produced by both non-heterocytous cultures were the AGs (Fig. 1a).

###### **Identification of HG analogs in non-heterocytous cyanobacteria**

As C_26_ - C_32_ alkyl diols, keto-diols etc. are more amenable to analysis by gas chromatography (GC) than by liquid chromatography (LC)^24^, we first examined the lipid extracts of the biomass of *Gloeocapsopsis crepidinum* LEGE 06123 and *Pleurocapsales* cyanobacterium LEGE 10410 by gas chromatography mass spectrometry (GC-MS). Based on published spectra of acid hydrolyzed HGs^24^ we examined extracted ion chromatograms (MS^2^) of *m/z* 219, a diagnostic fragment found in both silylated triols, tetrols and triol-ketos. In both strains we detected a compound of interest eluting at 29.9 min, which gave rise to a spectrum (Supplementary Fig. 19a) with a diagnostic *m/z* 219 fragment (indicative of a 1,3-diol) but lacking a *m/z* 117 fragment, diagnostic for a component with a hydroxy moiety at the ω-1 position, and lacking ions at *m/z* 58 and 130, diagnostic of a keto group at the ω-1 position^24^. To further elucidate the structure of this component, we carried out re-analysis after silylation with deuterated N,O-bis(trimethylsilyl)trifluoroacetamide (BSFTA). This gave rise to a mass spectrum (Supplementary Fig. 19b) with an ion of interest that contained either 9 or 18 deuterations, indicative that the fragment ion contained either one or two silylated hydroxy (OH) groups. Of particular interest was the ion at *m/z* 397 in the non-deuterated component, which appeared at *m/z* 406 after deuterated silylation. This 9 Da increase indicates that this fragment contains one hydroxy moiety, and hence the ion at *m/z* 397 was assigned as a C_22_ with a single silylated OH group. This ion assignment, in combination with the presence of the *m/z* 219 ion indicative of a 1,3-diol and the absence of diagnostic ions for oxygen moieties at the ω-1 position allowed us to tentatively assign this component as tetracosane-1,3-diol.

Additionally, a later eluting component was detected in *Pleurocapsales* cyanobacterium LEGE 10410, eluting at 45.1 min, while two similar other late eluting components were detected in *Gloeocapsopsis crepidinum* LEGE 06123, eluting at 43.6 and 52.9 min. All three late eluting components gave rise to spectra indicative of compounds containing a silylated sugar moiety, e.g. with dominant ions at *m/z* 204 and 217 (ref. ^39^) (Supplementary Fig. 20)*.* Of note was the presence in all three spectra of an ion at *m/z* 397, providing evidence that these four components were also composed of tetracosane-1,3-diols. The sugar in *Pleurocapsales* cyanobacterium LEGE 10410 was identified (Supplementary Fig. 20a) as a hexose on comparison of its mass spectrum with library mass spectra (NIST Mass Spectral Library, Version 2.0, 2012). The two sugars in *Gloeocapsopsis crepidinum* LEGE 06123 (Supplementary Fig. 20b,c) were identified as pentose sugars, in pentopyranose form, based on spectral library matches (NIST).

To further probe into the structure of these sugar-bearing tetracosane-1,3-diols, we examined the extracts by ultra-high-performance liquid chromatography coupled to high-resolution mass spectrometry (UHPLC–HRMS). In both the lipid extracts of *Pleurocapsales* cyanobacterium LEGE 10410 and *Gloeocapsopsis crepidinum* LEGE 06123, we first screened for the presence of tetracosane-1,3-diol ([M+H]^+^ C_24_H_51_O_2_, *m/z* 371.389). This component was present in both extracts, as two isomers, eluting at 16.3 min (*Pleurocapsales* cyanobacterium LEGE 10410 and *Gloeocapsopsis crepidinum* LEGE 06123) and 19.3 min (*Pleurocapsales* cyanobacterium LEGE 10410 only), both of which gave rise to similar but not identical MS^2^ spectra (Supplementary Fig. 21). In addition, we detected three larger components which gave rise to fragment ions at *m/z* 371.389, distributed across the two strains. The first of these components, present only in *Gloeocapsopsis crepidinum* LEGE 06123, eluted at 14.67 min and gave rise to a [M+NH_4_]^+^ ion at *m/z* 652.499. Based on its accurate mass and its MS^2^ spectrum (Supplementary Fig. 22a), and based on the above-mentioned sugar assignment from GCMS analysis, this was tentatively assigned as a 1-(O-dipentopyranose)-3-tetracosanol, e.g. a tetracosane-1,3-diol attached to two pentopyranose sugars. The next component eluted at 14.81 min and gave rise to a [M+NH_4_]^+^ ion at *m/z* 550.468. Based on its accurate mass and its MS^2^ spectrum (Supplementary Fig. 22b) this was tentatively assigned as a 1-(O-hexose)-3-tetracosaneol, e.g. a tetracosane-1,3-diol attached to one hexose sugar, thus resembling the slightly longer H HG_26_ ol detected in several heterocytous strains (see section ‘Heterocytous cyanobacteria produce a wide diversity of new HGs’ above). This component was only detected in the lipid extracts of *Pleurocapsales* cyanobacterium LEGE 10410. Finally, the third component eluted at 15.22 min in both *Pleurocapsales* cyanobacterium LEGE 10410 and *Gloeocapsopsis crepidinum* LEGE 06123 and gave rise to a [M+NH_4_]^+^ ion at m/z 520.457. Based on its accurate mass and its MS^2^ spectrum (Supplementary Fig. 22c) this was tentatively assigned as a 1-(O-pentopyranose)-3-tetracosaneol, e.g. a tetracosane-1,3-diol attached to one pentopyranose sugar. The distribution of the tetracosane-1,3-diols and the 1-(O-sugar)-3-tetracosaneols arising from UHPLC–HRMS analysis agreed well with the analysis by GC-MS (Supplementary Table 19). However, it should be noted that intact polar lipid species have diverse degrees of ionization efficiencies during UHPLC–HRMS and hence the peak areas, in response units, of different components do not always accurately reflect their relative abundances in the source material.

###### **Potential production of HG analogs by the PKSs encoded by hgl islands of non-heterocytous cyanobacteria**

Thus, both non-heterocytous strains produced tetracosane-1,3-diol under all tested culturing conditions, a molecule that structurally resembles an AG of canonical HG biosynthesis but with a shorter 24-carbon chain and no keto or alcohol group at the ω-1 position (Fig. 6c, structure I). In addition, sugar-bound compounds of these molecules were present in higher abundance in both cultures and under all tested conditions, representing structural analogs to canonical HGs (Fig. 6c, structures II, III, and IV). In *Pleurocapsales* cyanobacterium LEGE 10410, these HG analogs contained mainly hexose (mean 64.8% ± 6.5% standard deviation) with a low abundance of pentopyranose (6.8% ± 1.4%), and in *Gloeocapsopsis crepidinum* LEGE 06123 pentopyranose (64.7% ± 3.2%) and dipentopyranose (26.2% ± 2.9%) (Supplementary Tables 9, 11 and 20). Given the structural resemblance to AGs and HGs of the molecules identified in these two phylogenetically distant strains (see Fig. 6a), we hypothesize that their *hgl* islands encode the enzymes responsible for their biosynthesis.

For further confirmation, we screened the heterocytous cultures for the presence of the newly identified HG analogs, and found them only in a single strain, *Calothrix* sp. CCY 0018 (Fig. 6a,b), which produced tetracosane-1,3-diol (3.0% of total HGs + HG analogs) and in higher abundance 1-(O-pentopyranose)-3-tetracosaneol (15.7%; Supplementary Table 11). The genome of this strain contains, in addition to an expected extended *hgl* island (homologs of 15 different genes) which is likely responsible for canonical HG biosynthesis, an additional *hgl* island with a gene composition that is similar to the *hgl* islands of the two non-heterocytous strains discussed above (*hgl* island with homologs of 9 different genes; *hgdCB*, *all5343*, and *hglBACGE_A_F*; Fig. 6d). In the concatenated HG biosynthesis gene tree, this additional island is closely related to the *hgl* island of *Pleurocapsales* cyanobacterium LEGE 10410 (Fig. 6b, Supplementary Fig. 16). Thus, the additional *hgl* island of *Calothrix* sp. CCY 0018 is not closely related to the more extended *hgl* islands or *hgl*-like islands of other heterocytous cyanobacteria but instead to the *hgl* islands of non-heterocytous cyanobacteria (see section ‘*Hgl* and *hgl*-like islands are present throughout *Cyanobacteriia*’ above). The close evolutionary association between the additional *hgl* island of *Calothrix* sp. CCY 0018 and that of *Pleurocapsales* cyanobacterium LEGE 10410, in addition to the shared production of HG analogs by these strains, provides further evidence for the biosynthesis of these molecules by an ancient phylogenetic lineage of the *hgl* island. Whereas the canonical HGs of *Calothrix* sp. CCY 0018 are likely produced by the enzymes encoded by its more extended *hgl* island (containing homologs of 15 different genes), the HG analogs may thus be produced by the enzymes encoded by its additional *hgl* island (with homologs of 9 different genes).

The structural differences between the HG analogs and canonical HGs—an alkyl chain containing only 24 carbon atoms compared to at least 26 in canonical HGs and the absence of a keto or alcohol group at the ω-1 position—may derive from different enzymatic activity of the encoded PKSs. Compared to the enzymatic activity encoded by the more extended *hgl* island in heterocytous cyanobacteria (Supplementary Fig. 1), where canonical HGs result from the omission of the first dehydration and reduction steps, the first condensation round of the PKS encoded by the *hgl* islands of *Pleurocapsales* cyanobacterium LEGE 10410, *Gloeocapsopsis crepidinum* LEGE 06123, and the additional *hgl* island of *Calothrix* sp. CCY 0018 could include the reduction of the keto group to an alcohol group, its dehydration into a double bond, and its subsequent reduction to a saturated carbon chain, thus resulting in a lipid lacking the ω-1 alcohol group. In addition, a lower number of extension cycles compared to canonical HG biosynthesis would result in a shorter alkyl chain. Alternatively, the HG analogs could result from modification of a canonical AG after biosynthesis by the PKS via a yet unidentified mechanism. In both cases, the attachment of the glucose headgroup to the tetracosane-1,3-diol could be carried out by a GT encoded by the island, for example the *all5343* homologs on the *hgl* island of *Pleurocapsales* cyanobacterium LEGE 10410 and on the additional *hgl* island of *Calothrix* sp. CCY 0018, or encoded anywhere else on the genome, for example the high-scoring *hglT* hit of *Gloeocapsopsis crepidinum* LEGE 06123.

Even though the exact enzymatic activity of the encoded PKS remains unknown, we note that the *hgl* islands of both non-heterocytous strains and the additional *hgl* island of *Calothrix* sp. CCY 0018 do not encode a homolog of the ketoacyl synthase *hglD*. Moreover, the expression region may encode other genes that are unrelated to the queried *hgl* island genes of *Anabaena* sp. PCC 7120 and which are therefore not included in our *hgl* island definition, but which may be involved in the biosynthesis of the HG analogs—including potentially carrying out condensation reactions where no functional groups are introduced or shortening of the alkyl chain.

###### **Implications for the evolution of the heterocyte**

We thus for the first time identify HG analogs in non-heterocytous cyanobacteria, which we speculate are produced by the *hgl* islands encoded by their genomes. Given that the molecules are produced by the non-heterocytous strains irrespective of nitrogen source or age of the culture, they are likely not involved in nitrogen fixation nor in (a process similar to) akinete formation. Even though the function of the HG analogs thus remains unknown, the presence of *hgdC* and *hgdB* homologs (encoding potential ABC transporter proteins) on the *hgl* islands of the strains that produce them, suggests that the HG analogs are exported across the inner membrane and cell wall. Here, they may be incorporated into a cell envelope like in heterocytes, or be released in the environment. Future investigations into the non-heterocytous *Cyanobacteriia* carrying an *hgl* island that we identified here may shed light on the functional role of these novel natural products.

If the *hgl* islands in non-heterocytous strains are indeed responsible for the production of the here identified HG analogs, the molecules may be ‘remnants’ of a biosynthetic process that predates LHeCCA. Neofunctionalization of the already existing *hgl* island in LHeCCA, which was previously not used for nitrogen fixation, could have enabled the formation of a new type of cell envelope. Chain length extension and/or the addition of a keto or alcohol group at the ω-1 position may have improved oxygen impermeability and enabled nitrogen fixation by the oxygen-sensitive nitrogenase enzyme in an increasingly oxygenated atmosphere. Alternatively, the here identified lipids may be different from the molecules produced by the ancestors of LHeCCA. For example, the divergent gene composition of *hgl* islands in non-heterocytous cyanobacteria today may suggest that the ancestors of LHeCCA already contained an almost ‘complete’ *hgl* island to produce canonical HGs, and the *hgl* islands of contemporary non-heterocytous cyanobacteria reflect differential loss of individual genes from their island. Since none of the *hgl* islands of non-heterocytous strains contain *hglT* homologs, the inclusion of this GT into the island may have been a ‘late’ event relatively close in time to LHeCCA. However, since very similar molecules are biosynthesized by at least three distantly related strains in *Cyanobacteriia* (*Pleurocapsales* cyanobacterium LEGE 10410, *Gloeocapsopsis crepidinum* LEGE 06123, and *Calothrix* sp. CCY 0018), a scenario involving a change in biosynthetic product would require multiple similar independent changes from the ancestral enzymatic functionality to that of the contemporary *hgl* islands in the three strains, or alternatively, multiple horizontal transfer events. Comprehensive high-resolution characterization of the lipids produced by non-heterocytous *Cyanobacteriia* with an *hgl* island and elucidation of the enzymatic potential encoded by their genomes will further our understanding of the evolution of the heterocyte.

### **Supplementary Figures**

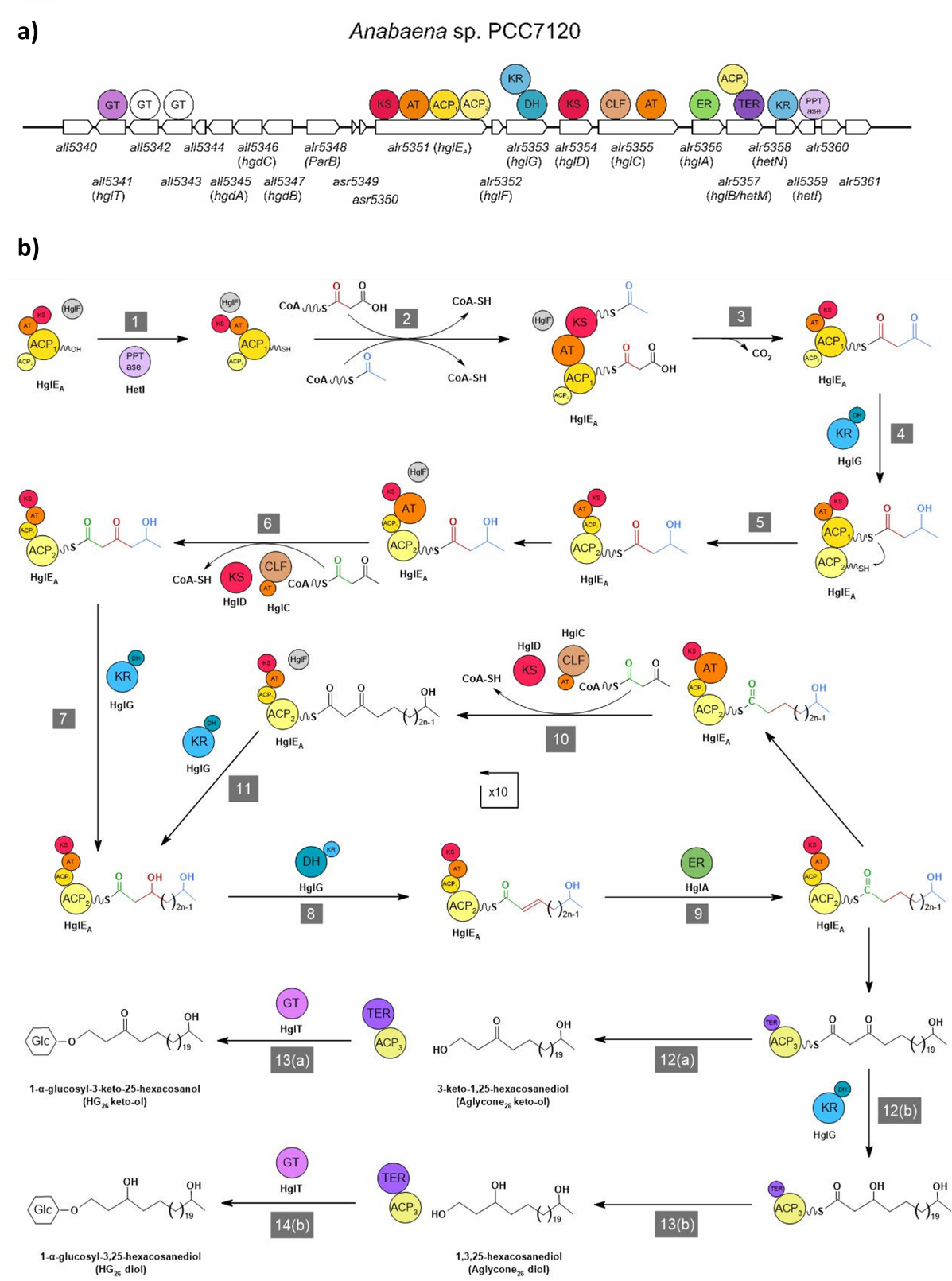

**Supplementary Fig. 1. Schematic representation of Anabaena sp. PCC7 7120 hgl island and the catalytic domains present in each gene (a) and representation of the proposed HG biosynthesis pathway carried out by the encoded proteins (b) (adapted from ref.** ^40^**).** (1) Activation of ACP domain in HglE_A_. (2) HglE_A_ acyl transferase (AT) adds malonyl group to ACP_1_ domain and β-ketoacyl synthase (KS) transfers acetyl residue to activated malonyl group. (3) Condensation catalyzed by HglE_A_ KS domain. (4) Reduction of ketone by HglG KR domain. (5) Chain translocation to the second ACP domain of HglE_A_. (6) Chain length checked by CLF domain of HglC and chain elongation catalyzed by HglE_A_ AT and HglD (KS). (7) Ketoreduction catalyzed by HglG KR domain. (8) Dehydration catalyzed by HglG DH domain. (9) Enoyl reduction catalyzed by HglA. (10) Chain elongation catalyzed by HglE_A_ AT and HglD (KS). Chain length checked by CLF domain of HglC. (11) Ketoreduction catalyzed by HglG KR domain. (12a) Chain termination catalyzed by HglB C-terminal thioester reductase (TER). (13a) Glucose moiety transferred to the aglycone by glycosyltransferase HglT. (12b) Ketoreduction catalyzed by HglG KR domain. (13b) Chain termination catalyzed by HglB C-terminal thioester reductase (TER). (14b) Glucose moiety transferred to the aglycone by glycosyltransferase HglT. GT, glycosyl transferase; KS, ketoacyl synthase; AT, acyl transferase; ACP, acyl carrier protein; KR, ketoreductase; DH, dehydratase; CLF, chain length factor; ER, enoyl reductase; TER, thioester reductase; PPTase, phosphopantetheinyltransferase.

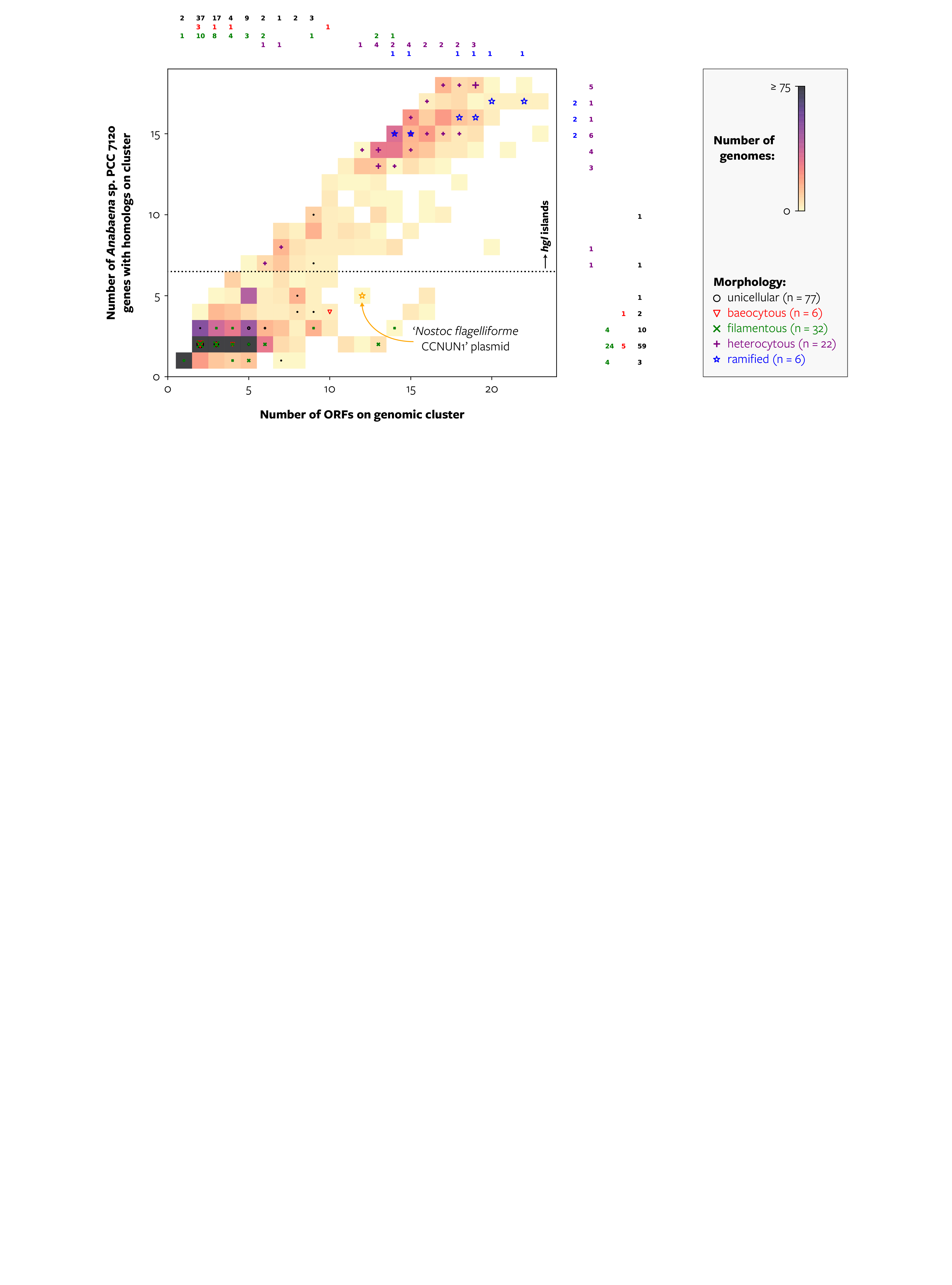

**Supplementary Fig. 2. The most extended genomic cluster of homologs of HG biosynthesis genes in genomes from the PATRIC genome database**^41^ **and in 14 newly sequenced genomes.** 3,579 genomes with the taxonomic assignment ‘phylum Cyanobacteria’ in the PATRIC ‘genome_lineage’ file and 14 genomes of heterocytous cultures were queried. The 3,339 genomes with ≥1 hit to an Anabaena sp. PCC 7120 HG biosynthesis gene are drawn in the plot, and only the most extended genomic cluster (in terms of number of Anabaena sp. PCC 7120 genes with homologs on the cluster) is shown per genome. X-axis represents the total length of the genomic cluster as number of open reading frames (ORFs), y-axis indicates the number of Anabaena sp. PCC 7120 genes with homologs on the cluster. The color of the squares indicates the number of genomes represented by that square. Colored numbers on the right and top of the graph indicate the number of genomes from ref. ^7^ present in each row and column, respectively, divided according to their morphology. Symbols within the square depict the same, where symbol size represents genome count. The orange star indicates a plasmid of strain Nostoc flagelliforme CCNUN1 (see Results and Discussion for details).

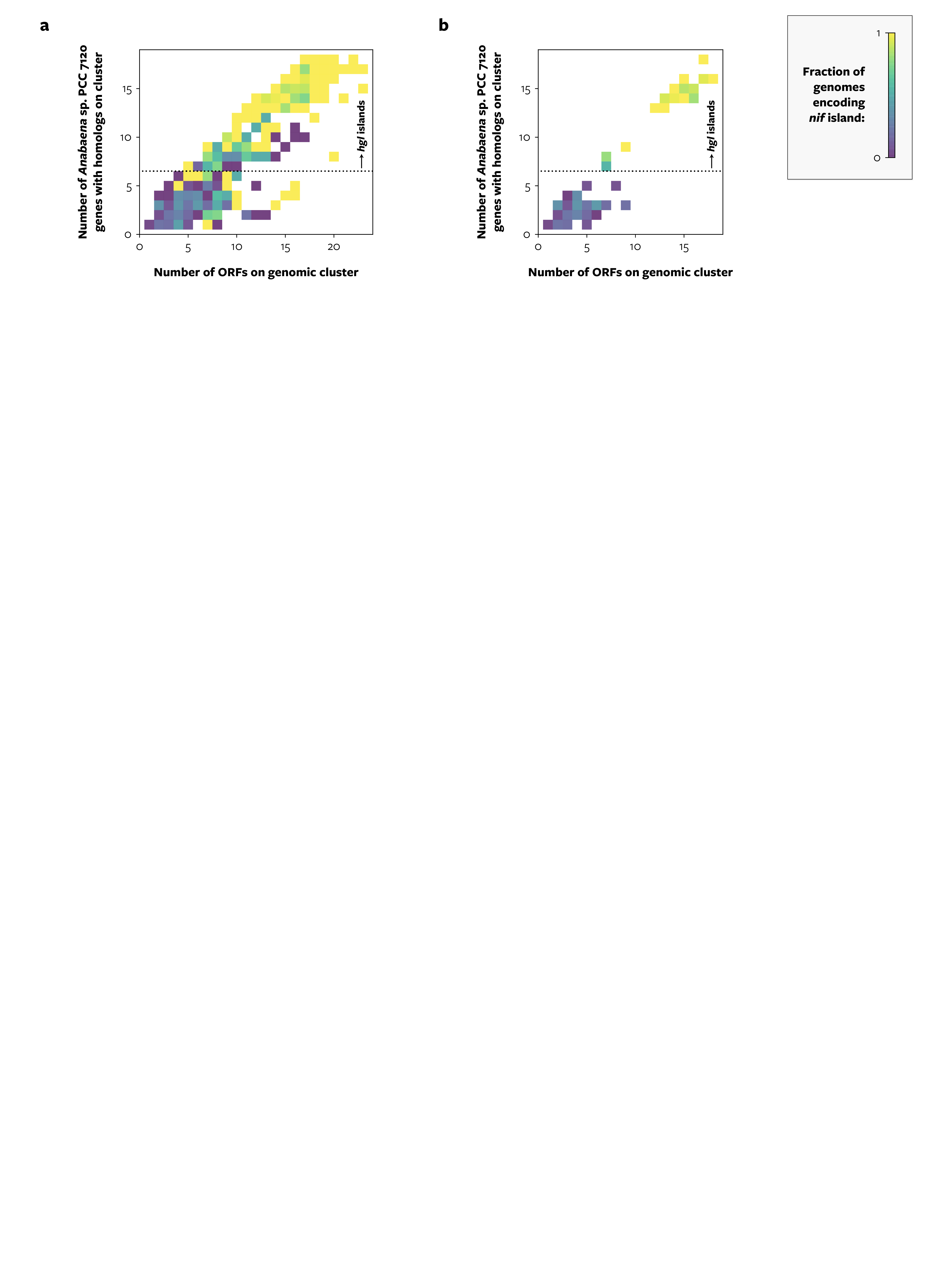

**Supplementary Fig. 3. Genomic co-occurrence of genomic clusters of homologs of HG biosynthesis genes and genomic clusters of nif genes.** **a**, The same plot as Supplementary Fig. 2 but with the squares colored according to the fraction of genomes in that square that encode a nif island (a genomic cluster containing homologs of ≥5 of the queried nif genes). **b**, The same plot as panel a but with only the squares that represent more than 10 genomes shown. Panel b represents 3,024 genomes, or 91% of the genomes depicted in panel a. ORF, open reading frame.

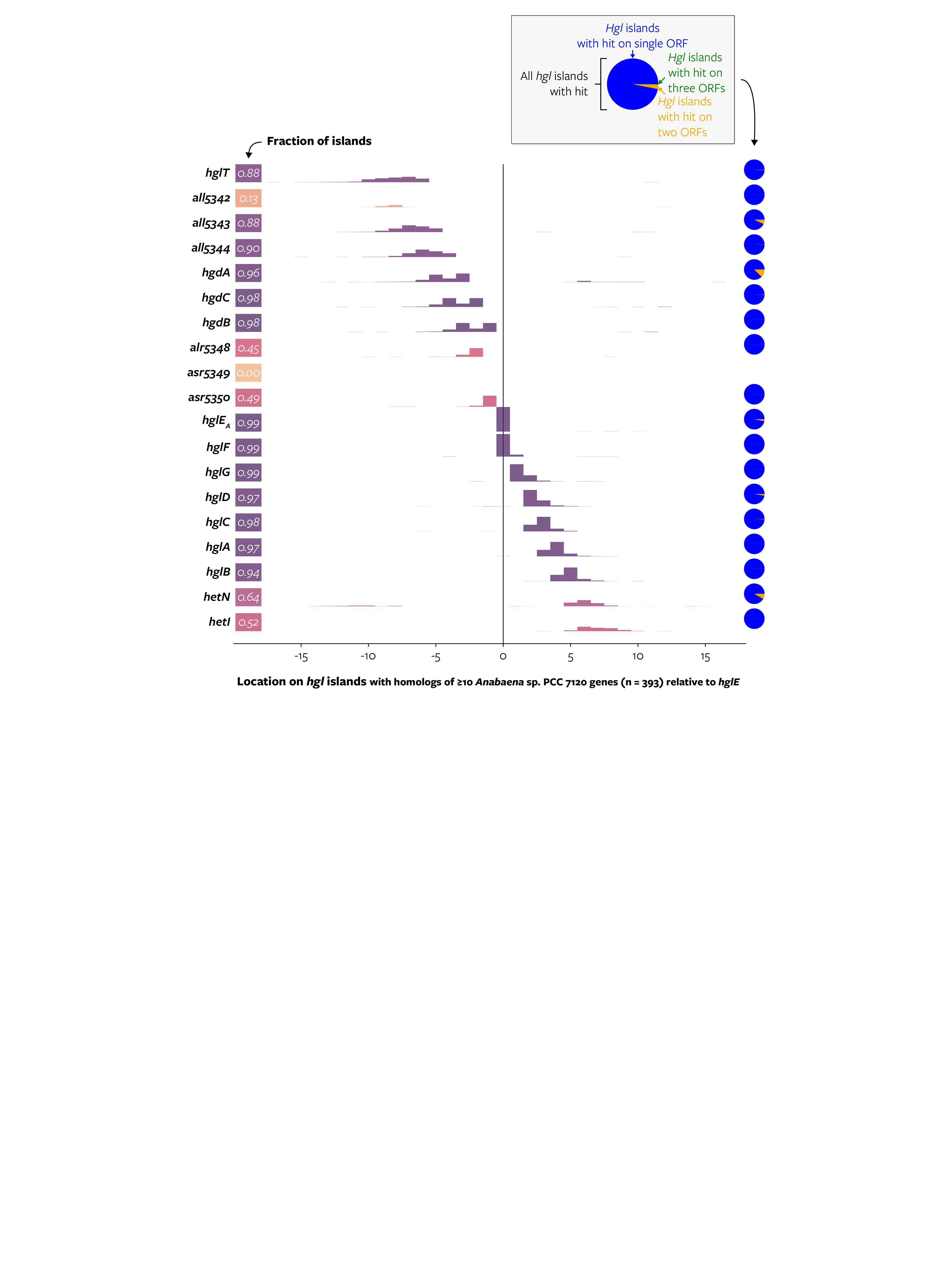

**Supplementary Fig. 4. Conservation of gene location on hgl islands with homologs of** ≥**10 Anabaena sp. PCC 7120 HG biosynthesis genes.** Histograms show the distribution of the location of hits for each queried gene. Location is measured as the distance in number of open reading frames (ORFs) between the hit and the ORF that contains an hglE_A_ hit, with orientation based on the hglE_A_ / hglG pair, or on the hglT / hglE_A_ pair if the hgl island did not contain an hglG hit. Three islands did not contain an ORF with an hglE_A_ hit, and in those cases, we based location on the hglG hit as if it was at location hglE_A_ + 2 with orientation based on the hglT / hglG pair. If an ORF contained multiple non-overlapping hits to the same gene, e.g. a duplication of hglE_A_, it was counted as 1 hit in this plot. Pie charts show the copy number of hits on the islands—for example, in 12% of the islands that contain an hgdA hit, two hgdA hits are present on two different ORFs. ORF, open reading frame.

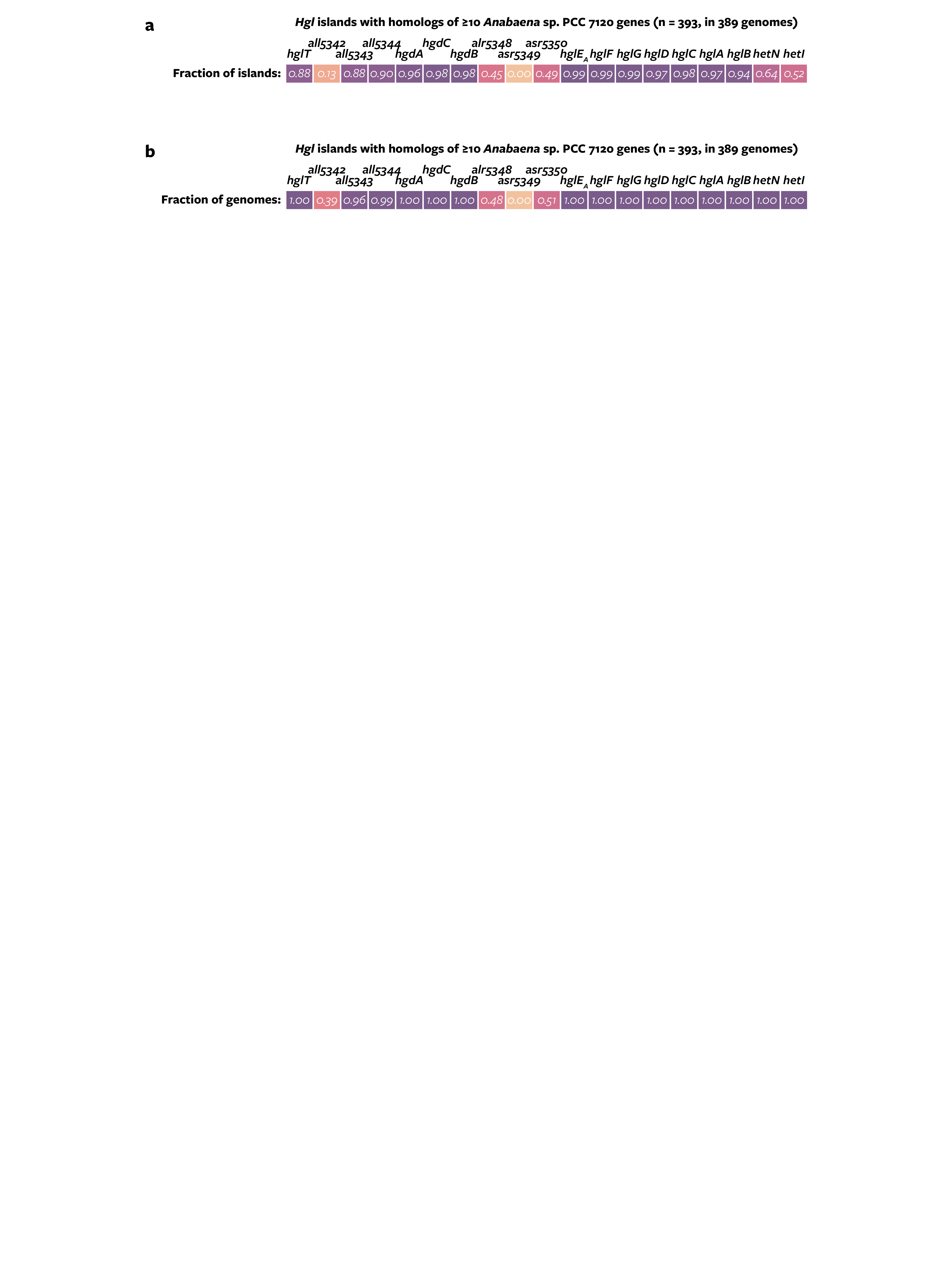

**Supplementary Fig. 5. Some genes are absent from the hgl island but present elsewhere on the genome. a**, Frequency of homologs of Anabaena sp. PCC 7120 HG biosynthesis genes on hgl islands containing homologs of ≥10 genes. The figure is identical to Fig. 2c. **b**, Frequency of homologs of HG biosynthesis genes in the genome of the cyanobacteria that encode the islands of panel a. Color-coding in panels a and b is for legibility only and reflects the numbers within the cells, with darker colors representing a higher fraction.

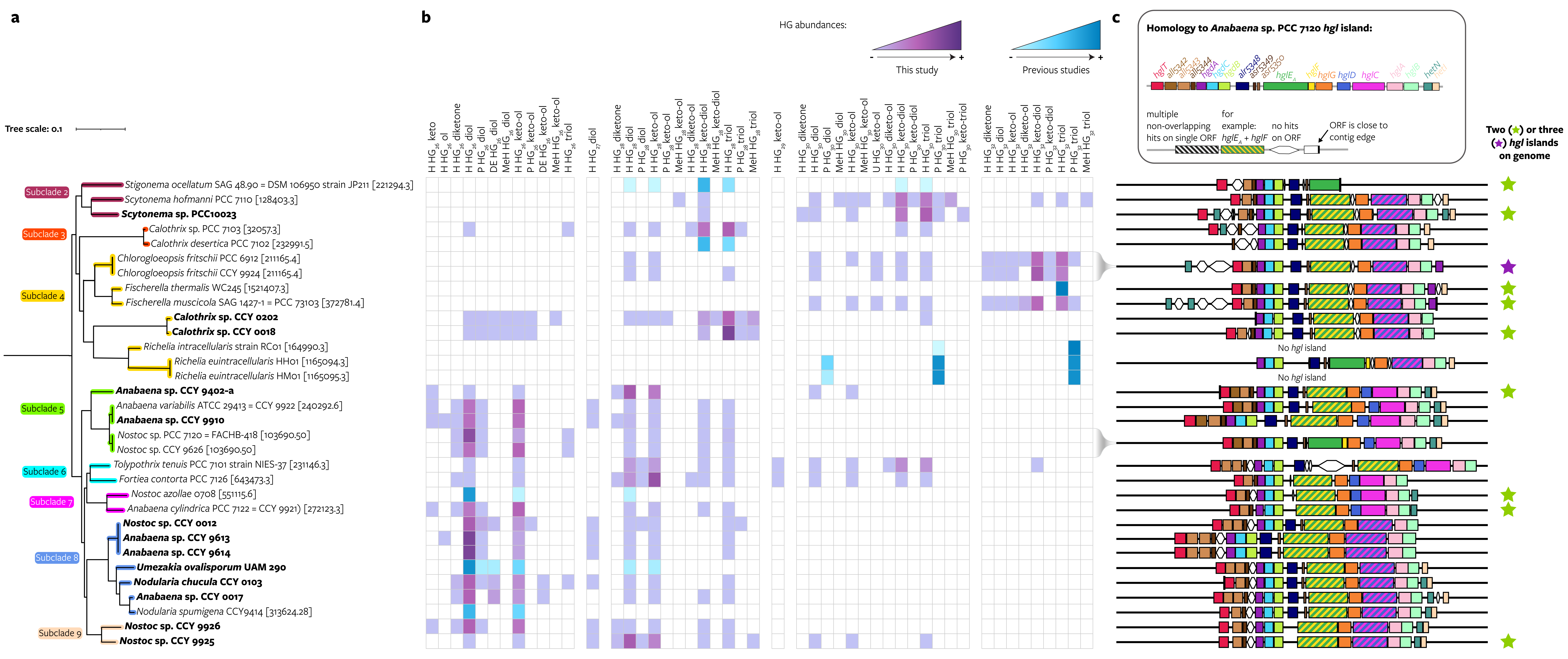

**Supplementary Fig. 6. Distribution of 49 HGs throughout heterocytous Cyanobacteriia.** The figure is identical to Fig. 3 except for panel b. **a**, Maximum likelihood cyanobacterial phylogeny based on 24 core vertically transferred genes including genomes with a known HG lipid profile (pruned from Fig. 2a). Genomes sequenced in this study are shown in bold. **b**, Heatmap of relative abundances of distinct HGs sorted according to their chain length. HG relative abundances obtained from this study are shown in purple and those obtained from literature are shown in blue. H, hexose; DeH, deoxyhexose; MeH, methyl-hexose; P, pentose; U, unknown. **c**, Schematic representation of the genes present on the most extended hgl island on the genome in terms of number of Anabaena sp. PCC 7120 genes with homologs on the island. Stars indicate the presence of two (green) or three (purple) hgl islands on the genome; these additional islands are not drawn. ORF, open reading frame; contig, contiguous sequence.

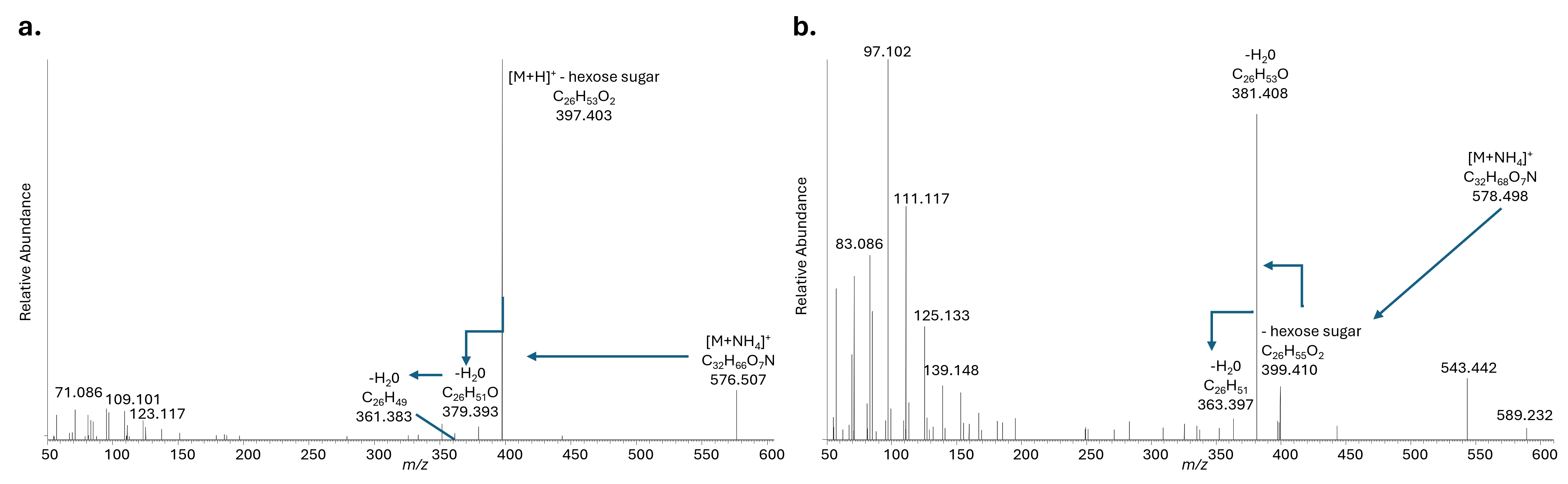
**Supplementary Fig. 7. UHPLC-HRMS MS^2^ spectra of unusual, novel HGs. a**, MS^2^ spectrum of the [M+NH_4_]^+^ ion at *m/z* 576.483 identified as hexose HG_26_ keto. **b**, MS^2^ spectrum of the [M+NH_4_]^+^ ion at *m/z* 578.499 identified as hexose HG_26_ ol.

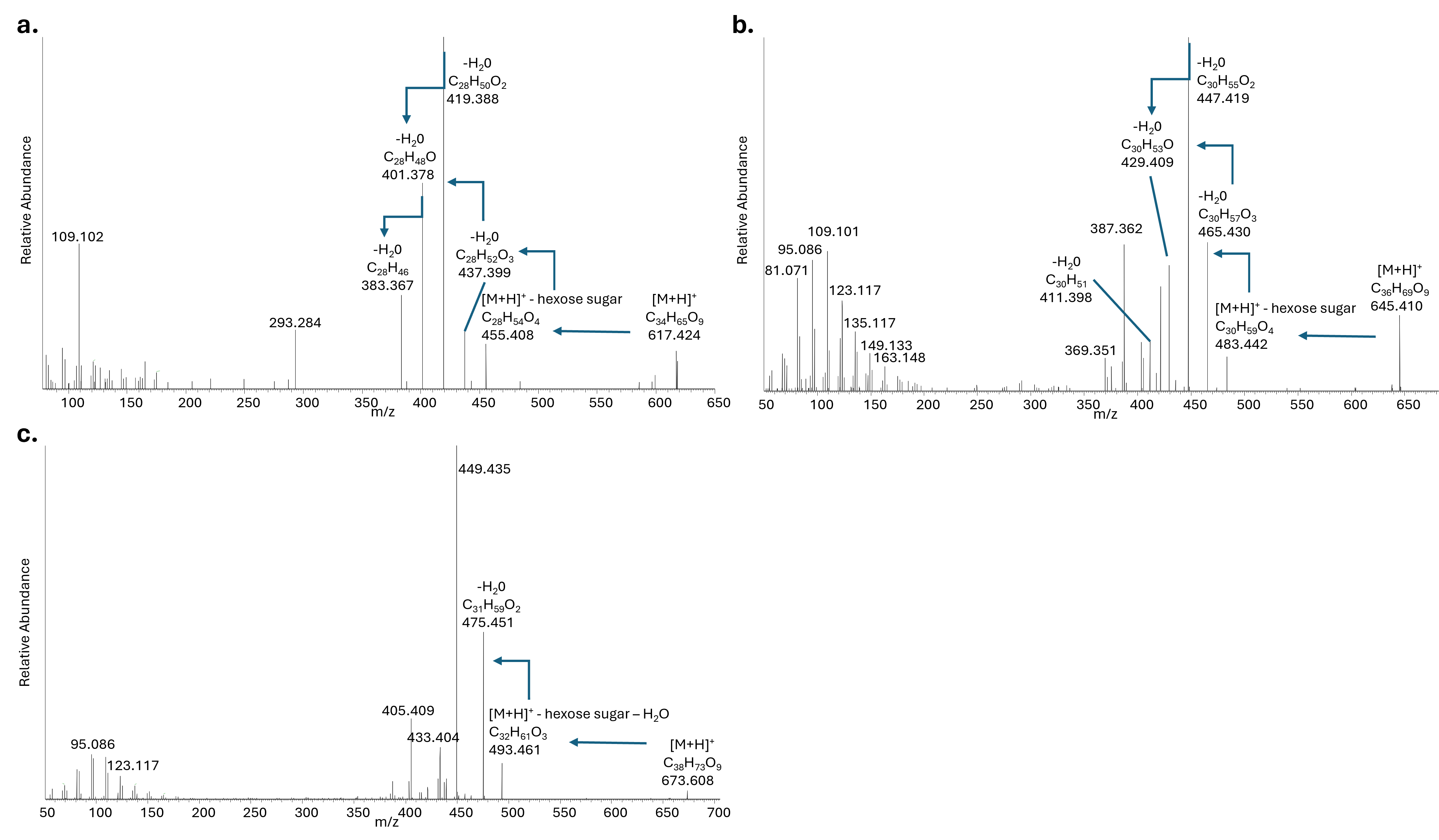
 **Supplementary Fig. 8. UHPLC-HRMS MS^2^ spectra of unusual, novel HGs. a**, MS^2^ spectrum of the [M+H]^+^ ion at *m/z* 617.462 identified as HG_28_ diketo-ol. **b**, MS^2^ spectrum of the [M+H]^+^ ion at *m/z* 645.494 identified as hexose HG_30_ diketo-ol. **c**, MS^2^ spectrum of the [M+H]^+^ ion at *m/z* 673.525 identified as hexose HG_32_ diketo-ol.

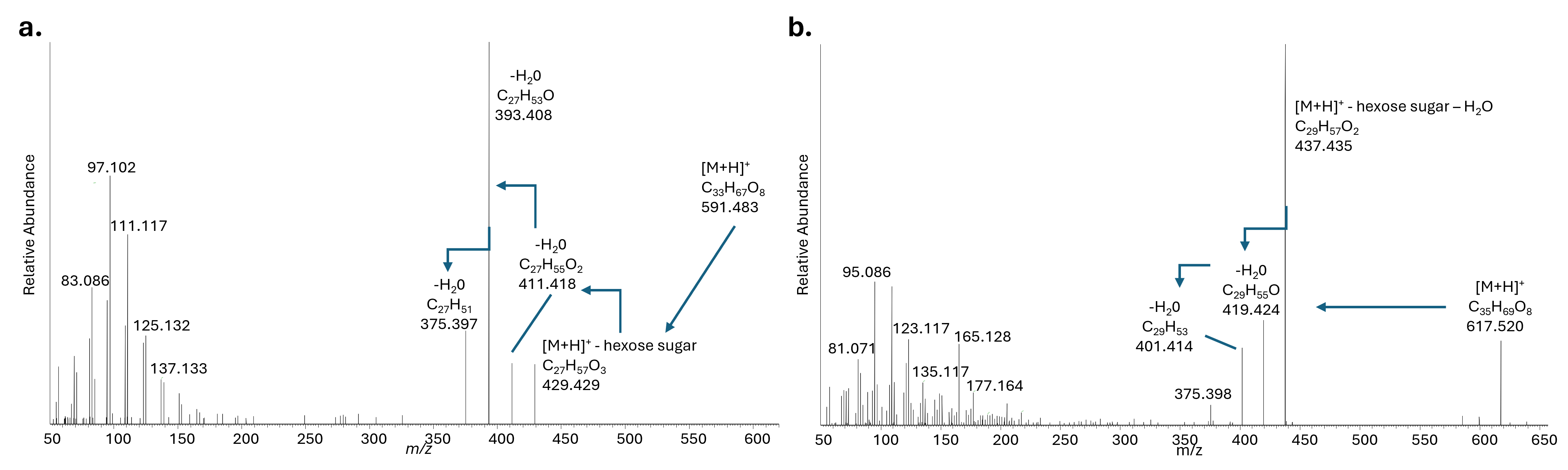
 **Supplementary Fig. 9. UHPLC-HRMS MS^2^ spectra of unusual, novel HGs. a**, MS^2^ spectrum of the [M+H]^+^ ion at *m/z* 591.484 identified as hexose HG_27_ diol. **b**, MS^2^ spectrum of the [M+H]^+^ ion at *m/z* 617.49 identified as hexose HG_29_ keto-ol.

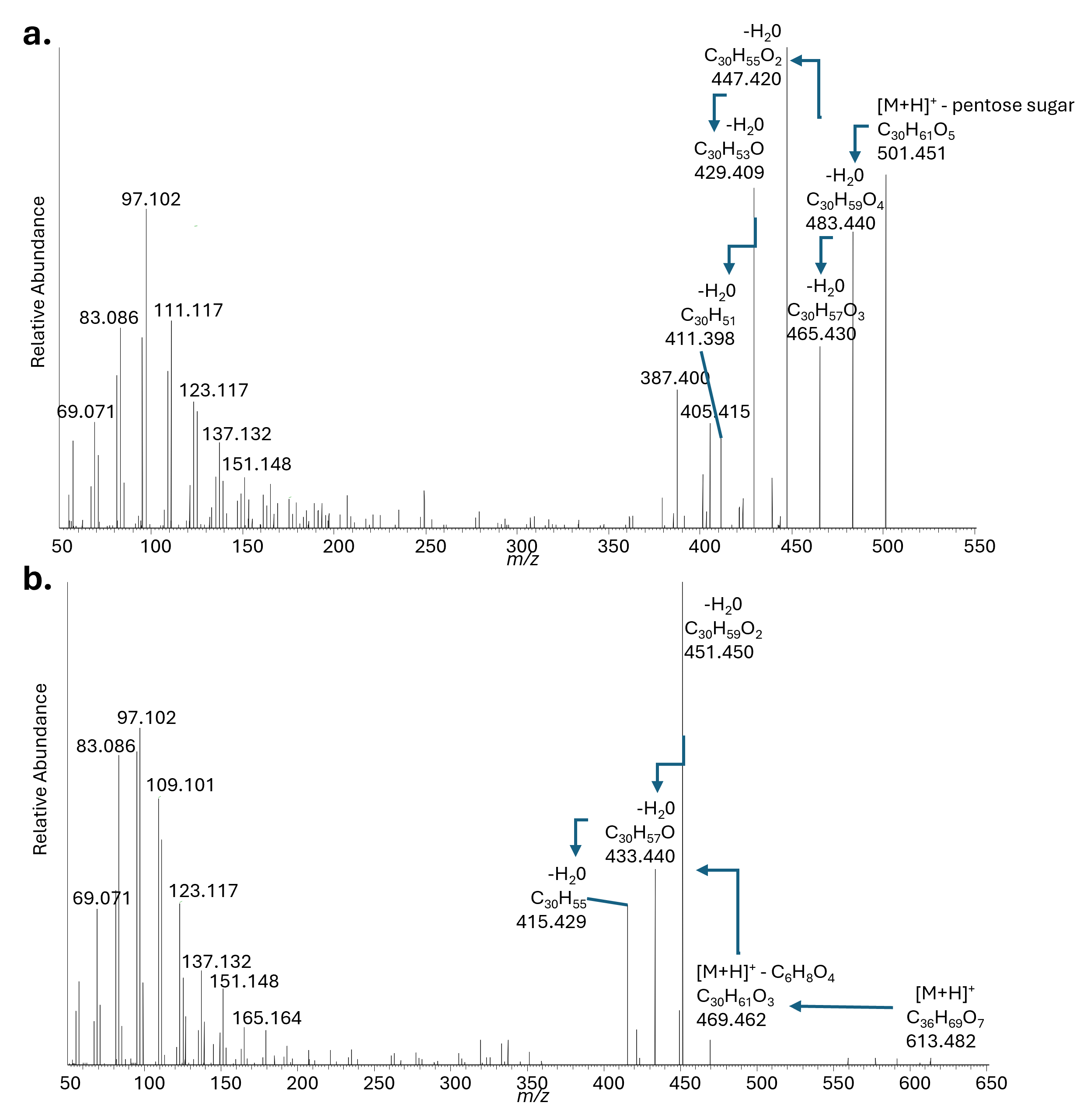
**Supplementary Fig. 10. a,** UHPLC-HRMS MS^2^ spectrum of the [M+H]^+^ ion at *m/z* 633.530 identified as pentose HG_30_ keto-triol. **b,** MS^2^ spectrum of the [M+H]^+^ ion at *m/z* 613.504 identified as HG-like compound, C_30_ keto-ol with an unknown headgroup.

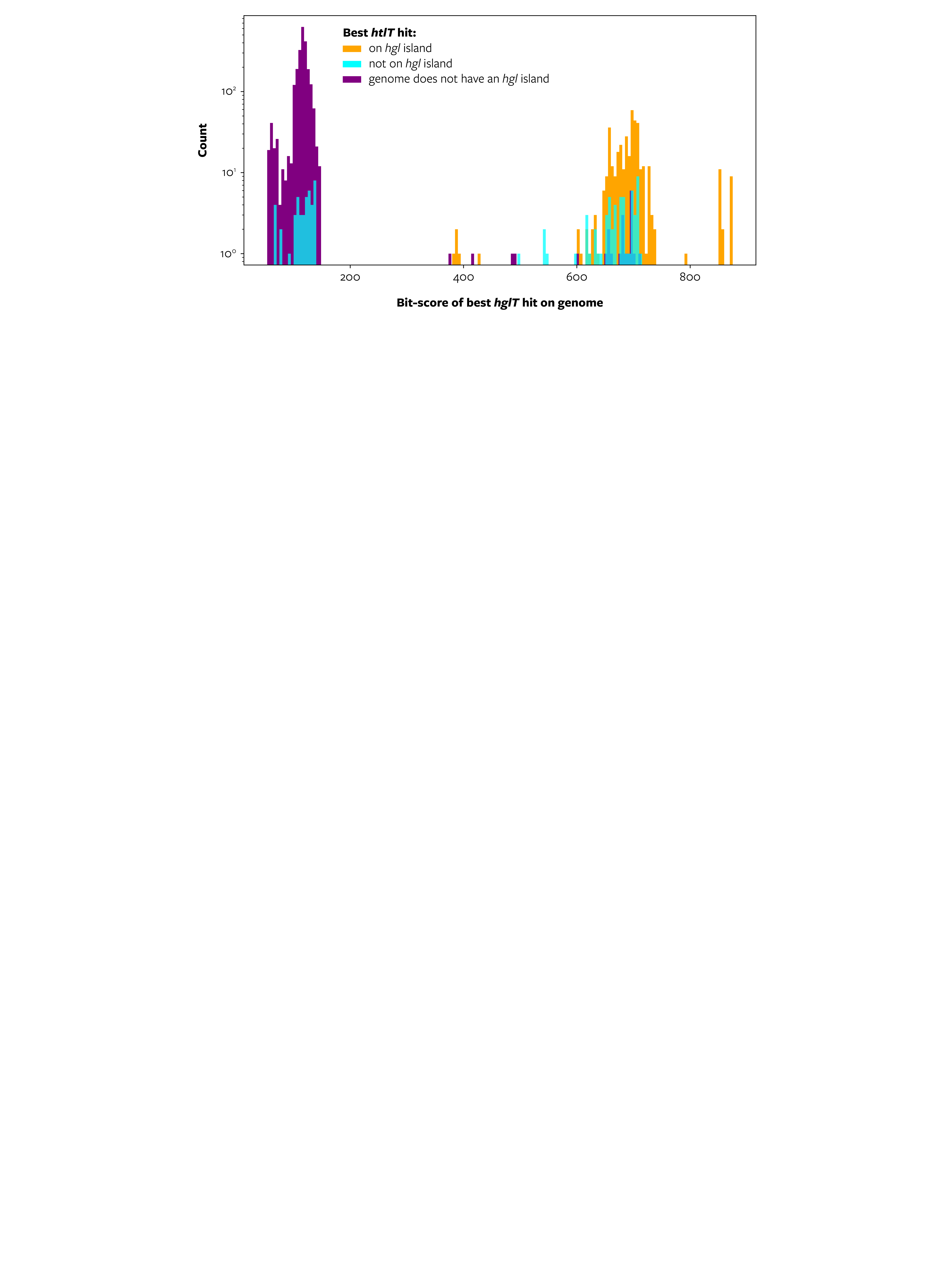

**Supplementary Fig. 11. Representation of the number of genomes (y-axis) for which the best hglT hit has a given bit-score (x-axis).** Hits with a bit-score <200 may be indicative of distant homologs with a potentially divergent function unrelated to HG biosynthesis. Colors represent whether the genome contains an hgl island as defined here (cyan and orange) and if so, if the best hglT hit is found within the island (orange) or elsewhere (cyan). Genomes without an island are shown in purple. Genomes were binned in cohorts of 5 bit-scores. Note the logarithmic scale of the y-axis.

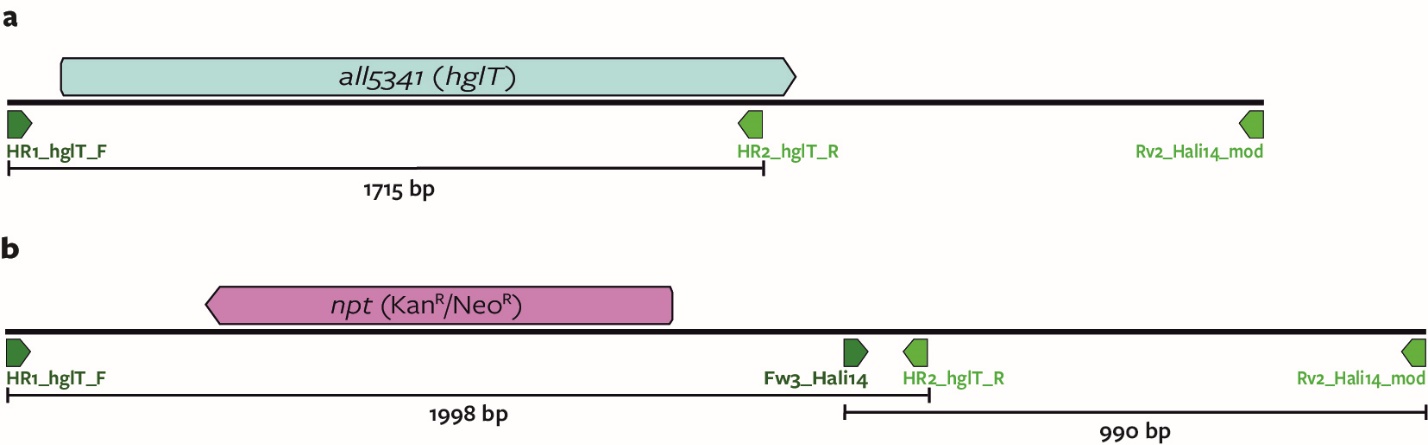

**Supplementary Fig. 12. Schematic representation of hglT (all5341). a,** in the wild-type Anabaena sp. PCC 7120 strain **b,** in the ∆hglT mutant strain where hglT is replaced by npt gene conferring resistance to kanamycin (Kan^R^) and neomycin, (Neo^R^)**.** HR1_hglT_F, HR2_hglT_R, Rv2_Hali14_mod and Fw3_Hali14, primer annealing sites. Lines and numbers indicate the expected length of the PCR product.

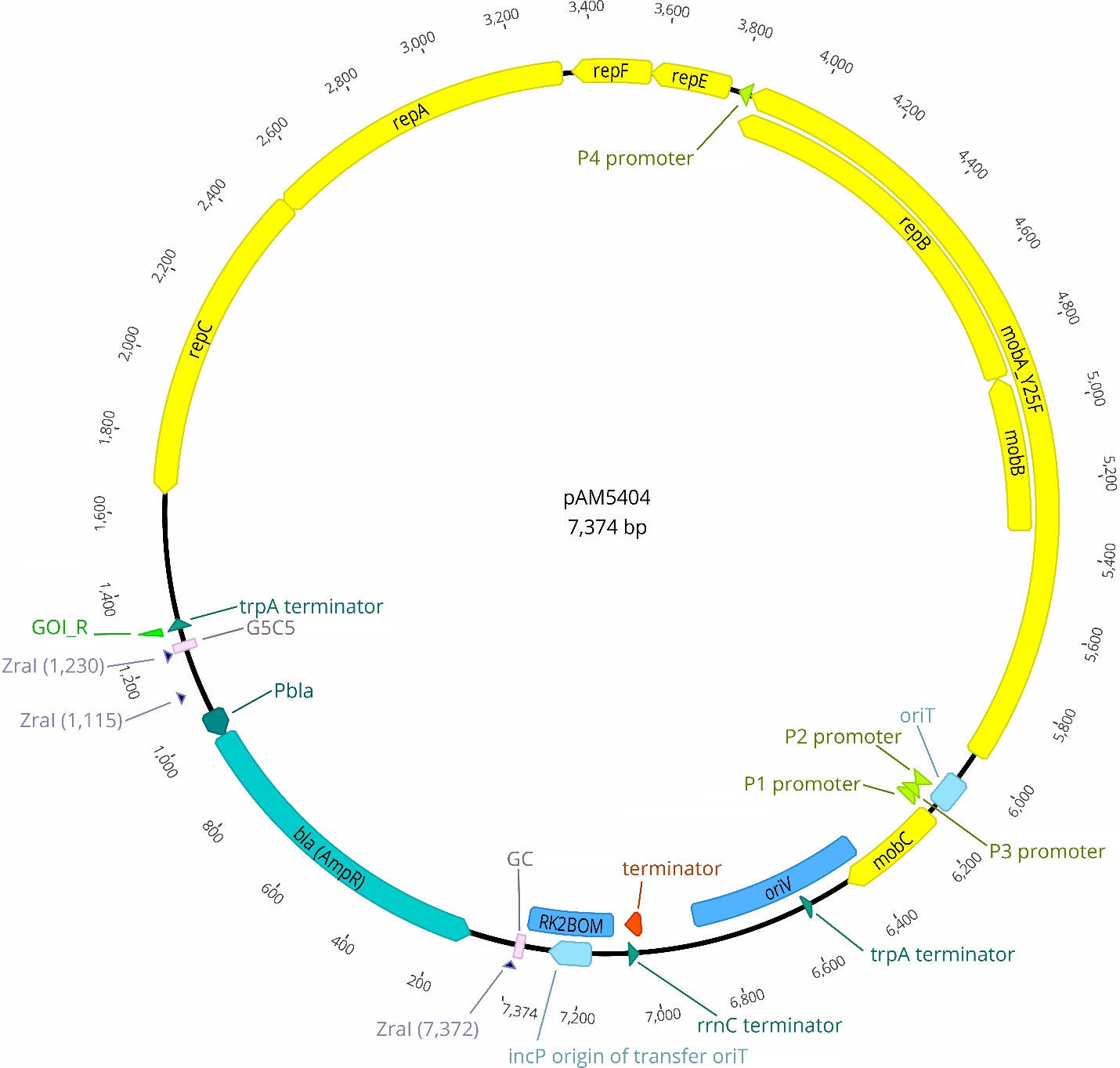

**Supplementary Fig. 13. Schematic representation of the pAM5404 cargo plasmid.** repA, repB, repC, repE and repF, genes required for plasmid replication; mobA, mobB and mobC, genes responsible for plasmid mobilization, mobA contains Y25F mutation responsible for improving cloning efficiency; oriV, RSF1010 broad-host-range plasmid origin of replication; oriT, RSF1010 plasmid origin of transfer; incP, RP4 conjugal plasmid origin of transfer; RK2bom, RK2 conjugal plasmid origin of transfer; P1, P2 and P3, RSF1010 native promoters; P_bla_ , ampicillin antibiotic resistance cassette promoter; bla (AmpR), ampicillin antibiotic resistance cassette; ZraI, restriction sites for ZraI enzyme; GOI_R, primer annealing site. GC and G5C5, GC-adaptor sequences described in ref. ^42^ and used for cloning of the GOIs and antibiotic resistance cassette.

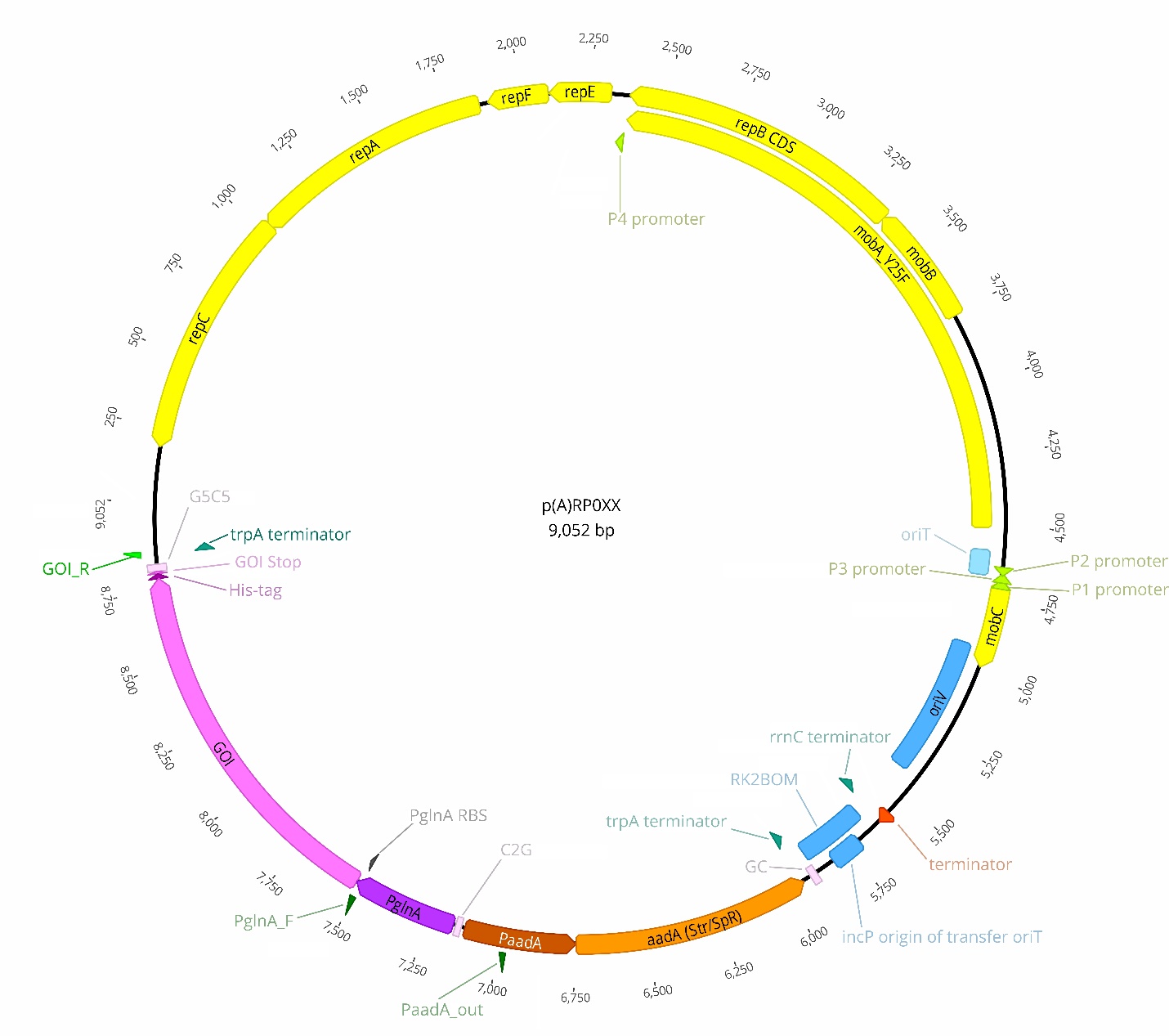

**Supplementary Fig. 14. Common schematic representation of plasmids pRP012-14, pRP019-022.** pRP019 and pRP020 do not possess a His-tag. GOI, gene of interest (all5341, RINTHH_17770, RINTHH_20790, RINTHH_5560-RINTHH_5570); P_glnA_, glnA promoter; P_aadA_, Streptomycin and spectinomycin resistance cassette promoter, aadA, spectinomycin and streptomycin resistance cassette. repA, repB, repC, repE and repF, genes required for plasmid replication; mobA, mobB and mobC, genes responsible for plasmid mobilization, mobA contains Y25F mutation responsible for improving cloning efficiency; oriV, RSF1010 broad-host-range plasmid origin of replication; oriT, RSF1010 plasmid origin of transfer; incP, RP4 conjugal plasmid origin of transfer; RK2bom, RK2 conjugal plasmid origin of transfer; P1, P2 and P3, RSF1010 native promoters; P_bla_ , ampicillin antibiotic resistance cassette promoter; bla (AmpR), ampicillin antibiotic resistance cassette; ZraI, restriction sites for ZraI enzyme; GOI_R, primer annealing site. GC and G5C5, GC-adaptor sequences described in ref. ^42^ and used for cloning of the GOIs and antibiotic resistance cassette.

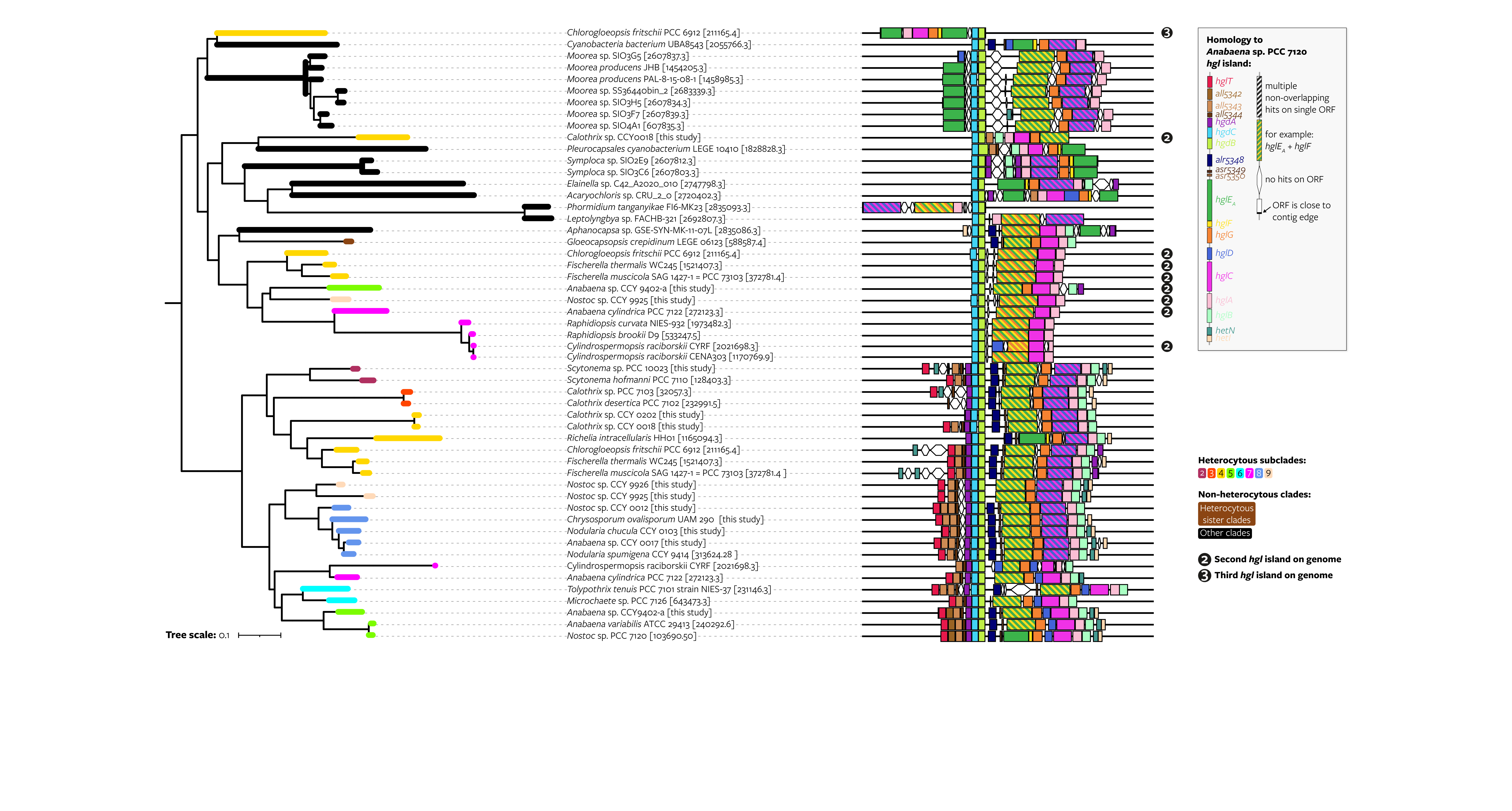

**Supplementary Fig. 15. Phylogenetic relationship between selected hgl and hgl-like islands.** The maximum likelihood phylogenetic tree of concatenated hgdCB and hglE_A_FGCA homologous sequences of hgl islands is pruned from Fig. 4. All hgl islands from strains with a known HG lipid profile and from all non-heterocytous cyanobacterial strains that contain an hgl island and that are shown in Fig. 4 are included. In addition, the islands of Raphidiopsis curvata NIES-932, Cylindrospermopsis raciborskii CENA303, and Raphidiopsis brookii D9 are included because they are non-diazotrophic cyanobacteria from within the heterocytous clade. Their single islands cluster with the hgl-like islands of heterocytous cyanobacteria instead of with their more extended hgl islands. In addition, the two islands of Cylindrospermopsis raciborskii CYRF are included because its hgl islands lack hglT. ‘First’, ‘second’, and ‘third’ hgl islands are based on the presence of other islands on the genome, where the ‘first’ hgl island (not marked) is the most extended island in terms of number of Anabaena sp. PCC 7120 genes with homologs on the island, the second hgl island (marked with a 2) the second-most extended island, and the third island (marked with 3) the third-most extended island. ORF, open reading frame; contig, contiguous sequence.

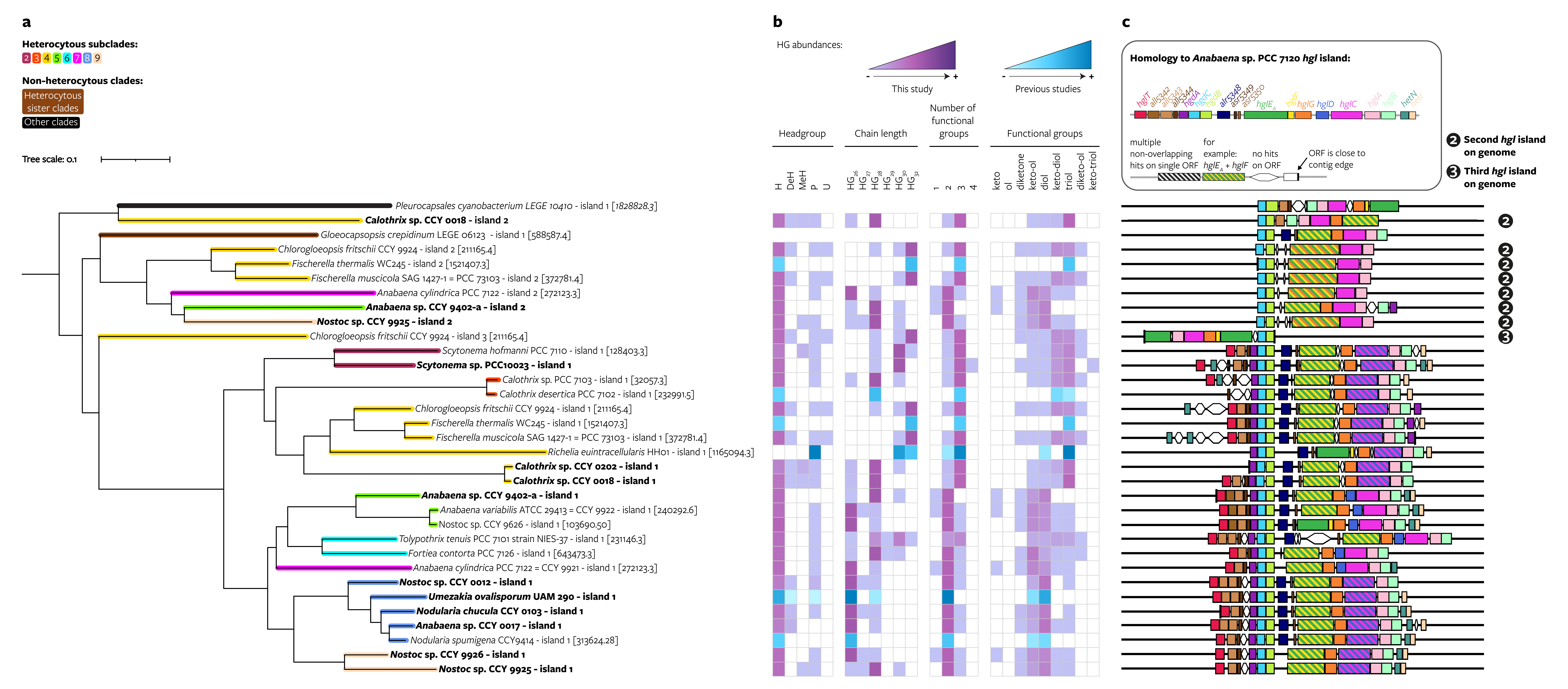

**Supplementary Fig. 16. Distribution of 49 HGs grouped according to their structure’s characteristics throughout Cyanobacteriia and plotted on the hgl island phylogeny. a**, Maximum likelihood phylogeny of the hgl island created using a concatenated alignment of homologous sequences of 7 HG biosynthesis genes (hgdCB and hglE_A_FGCA) that are often present on hgl islands, including only genomes with a known HG lipid profile (pruned, see Online Methods). Genomes sequenced in this study are shown in bold. Some genomes contain multiple hgl islands and are thus present in the tree multiple times. **b**, Heatmap of HG relative abundances grouped according to the headgroup (H, hexose; DeH, deoxyhexose; MeH, methyl-hexose; P, pentose; U, unknown), chain length, number and type of functional groups of each HG. HG relative abundances obtained from this study are shown in purple and those obtained from literature are shown in blue. **c**, Schematic representation of the genes present on the hgl island. When more than one island is present in the genome of the strain, the same HG abundances heatmap is shown for each island. ‘First’, ‘second’, and ‘third’ hgl islands are based on the presence of other islands on the genome, where the ‘first’ hgl island (not marked) is the most extended island in terms of number of Anabaena sp. PCC 7120 genes with homologs on the island, the second hgl island (marked with a 2) the second-most extended island, and the third island (marked with 3) the third-most extended island. ORF, open reading frame; contig, contiguous sequence.

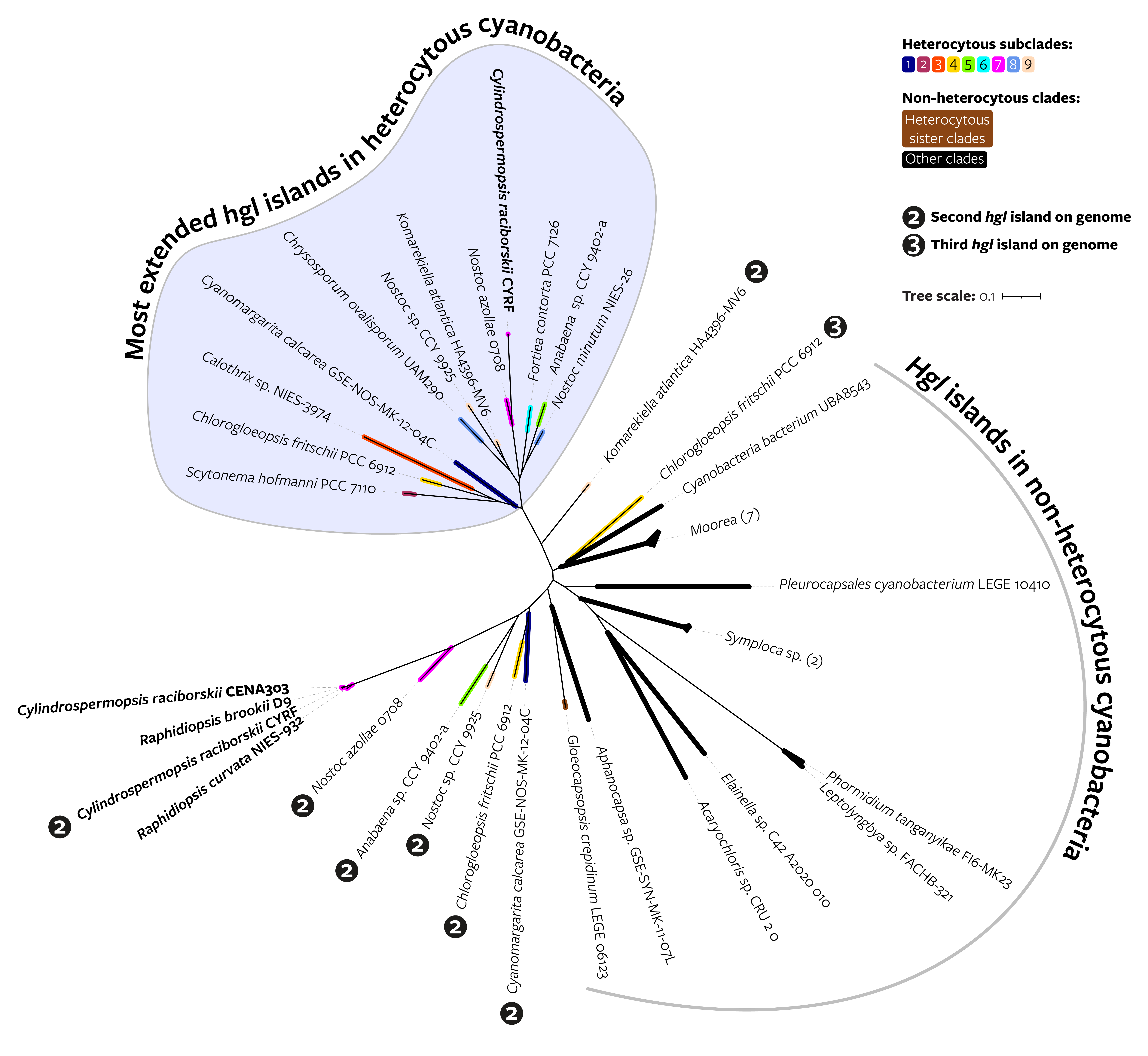

**Supplementary Fig. 17. Unrooted maximum likelihood phylogenetic tree of concatenated hgdCB and hglE_A_FGCA homologous sequences of hgl islands from selected genomes (pruned from Fig. 4).** One representative was chosen from each heterocytous subclade, and all the hgl islands from outside the heterocytous clade in Fig. 4 are included. In addition, the hgl-like islands of Raphidiopsis curvata NIES-932, Cylindrospermopsis raciborskii CENA303, and Raphidiopsis brookii D9 are included because they are non-diazotrophic cyanobacteria from within the heterocytous clade. ‘First’, ‘second’, and ‘third’ hgl islands are based on the presence of other islands on the genome, where the ‘first’ hgl island (not marked) is the most extended island in terms of number of Anabaena sp. PCC 7120 genes with homologs on the island, the second hgl island (marked with a 2) the second-most extended island, and the third island (marked with 3) the third-most extended island. Numbers within parentheses indicate the number of collapsed branches in that clade. BV-BRC genome identifiers of each leave are listed in Supplementary Table 18.

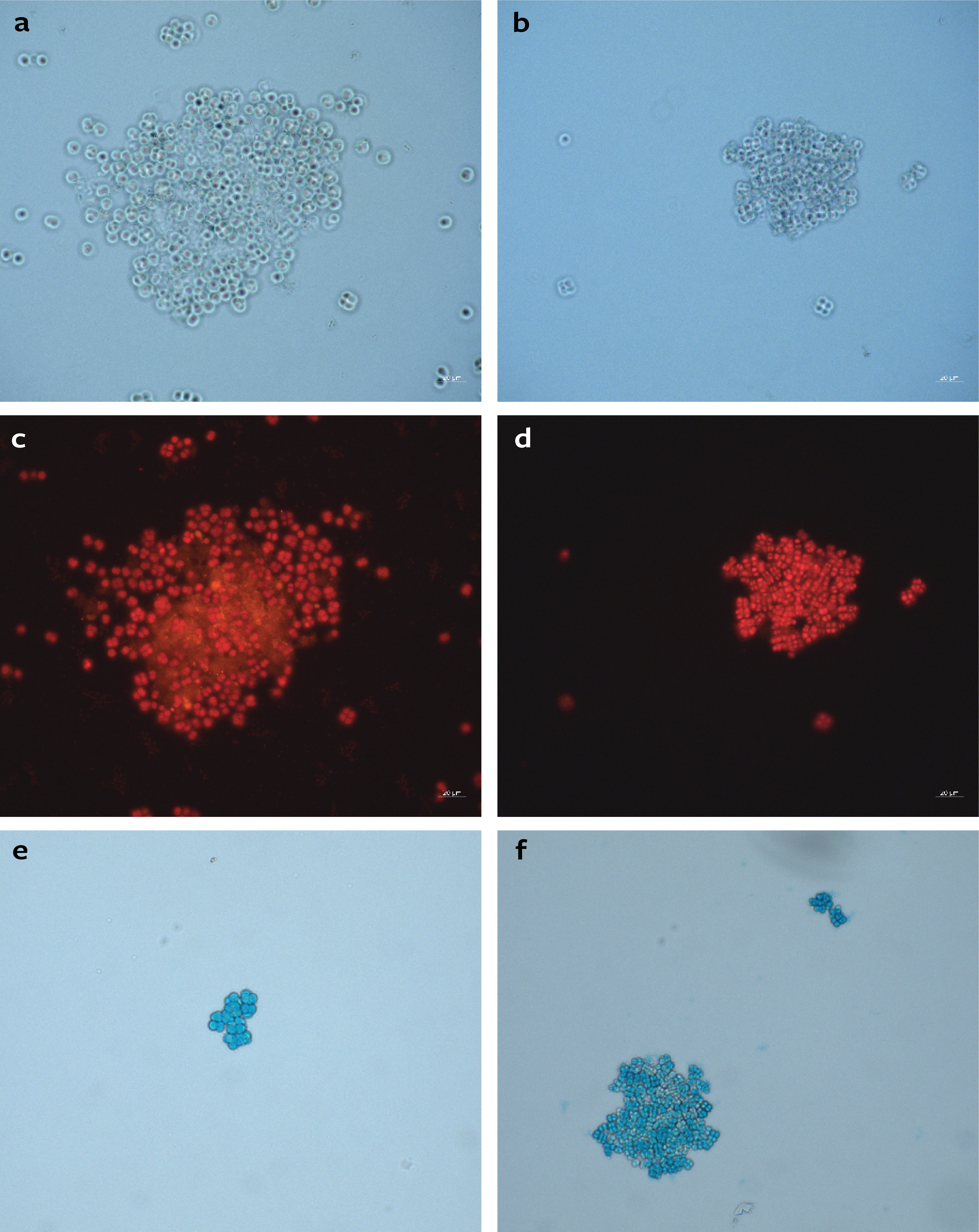

**Supplementary Fig. 18. Microscopic analysis of Gloeocapsopsis crepidinum LEGE 06123 (a,c,e) and Pleurocapsales cyanobacterium LEGE 10410 (b,d,f) grown in nitrogen-replete media for 38 days (a,b) and 77 days (c-f).** Panels a-d show lipids stained with Nile Red and analyzed using bright field (a-b) and epifluorescence microscopy (c-d). Panels e and f show polysaccharides stained with Alcian Blue.

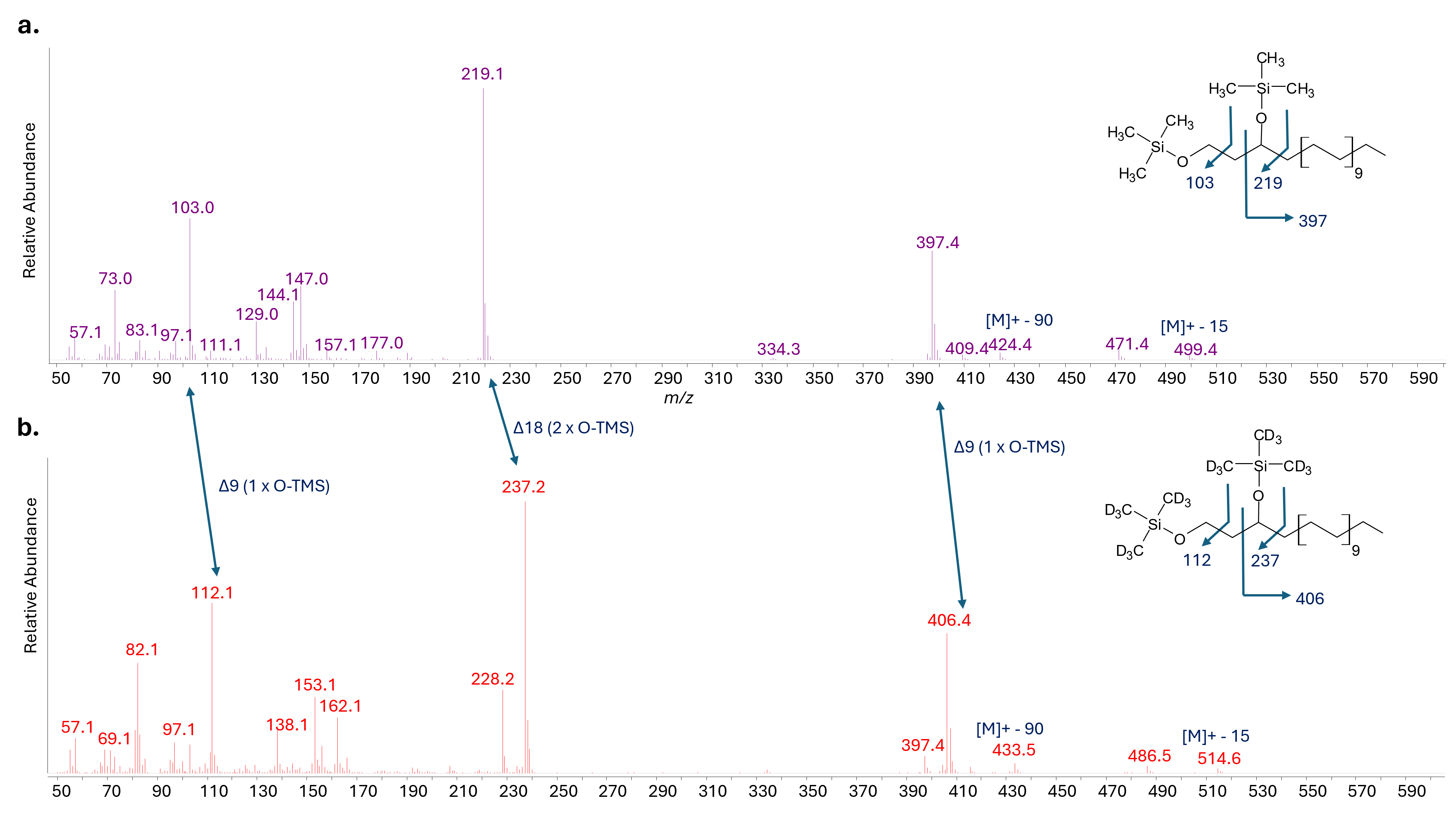

**Supplementary Fig. 19. GC–MS spectra of compound eluting at 29.9 min after** **a**, silylation with BSFTA and **b**, with deuterated BSTFA.

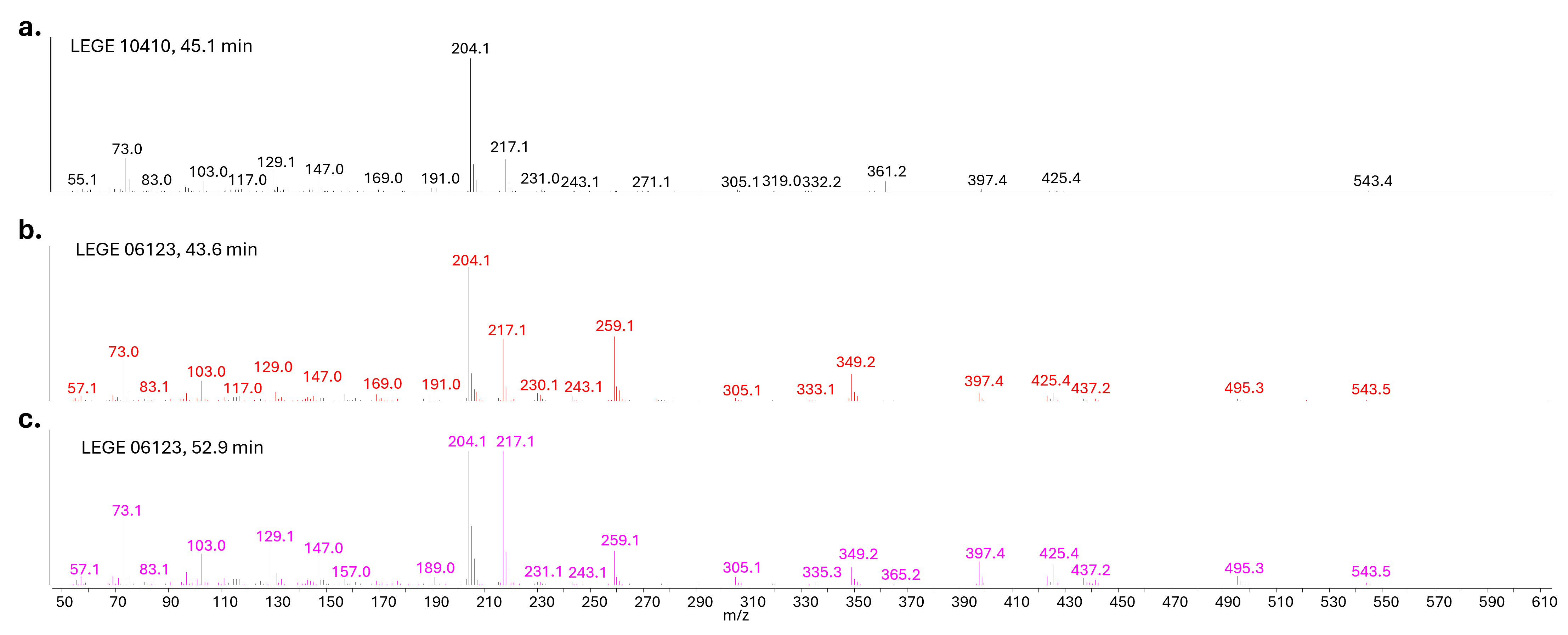

**Supplementary Fig. 20. GC–MS spectra of tetracosane-1,3-diols eluting at a**, 35.1 min with a hexose sugar headgroup, **b**, at 43.6 min and **c**, 52.9 min, both with a pentopyranose headgroup.

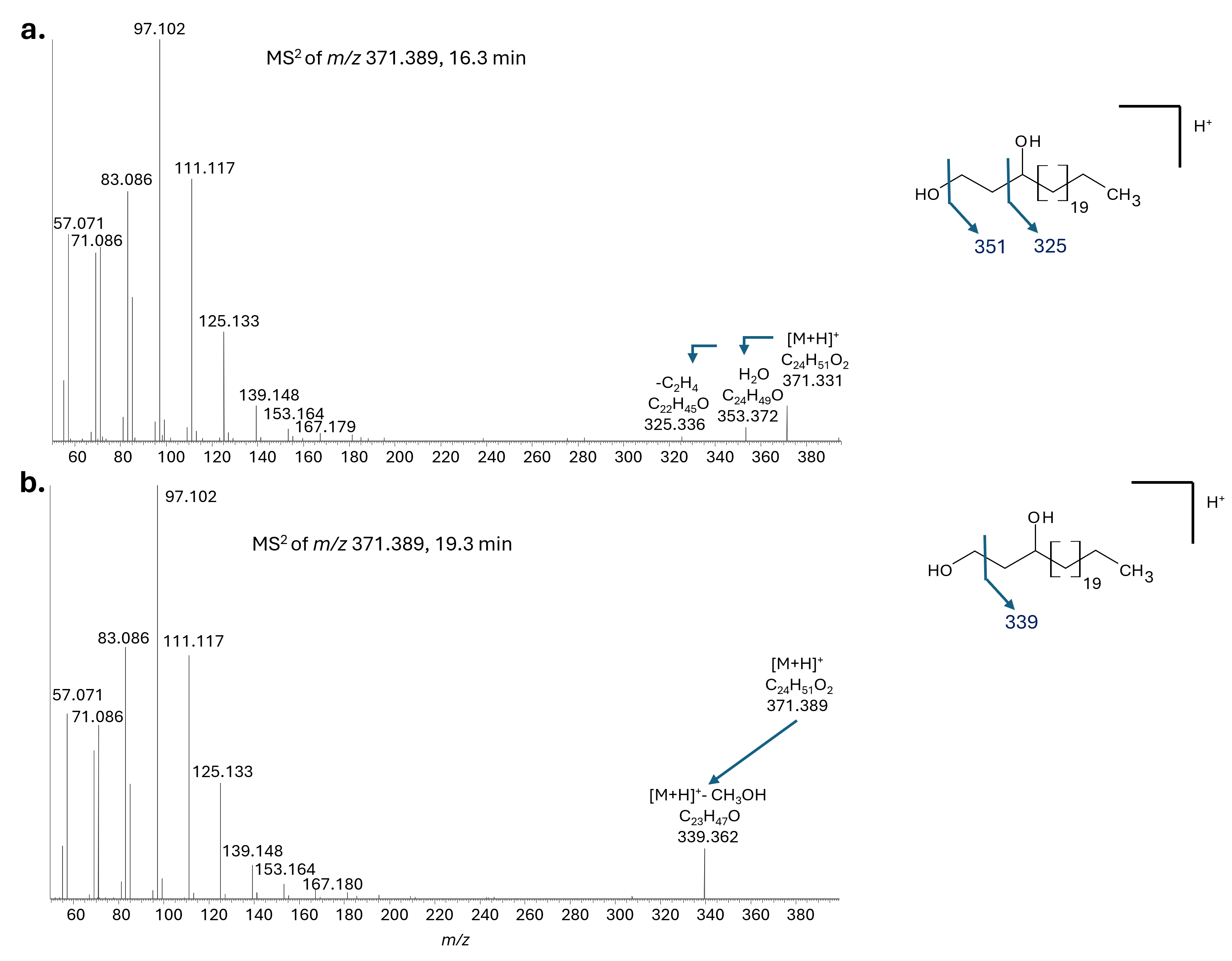

**Supplementary Fig. 21. UHPLC-HRMS MS^2^ spectra of tetracosane-1,3-diol ([M+H]^+^ C_24_H_51_O_2_, *m/z* 371.389) eluting at a**, 16.3 min and **b**, 19.3 min.

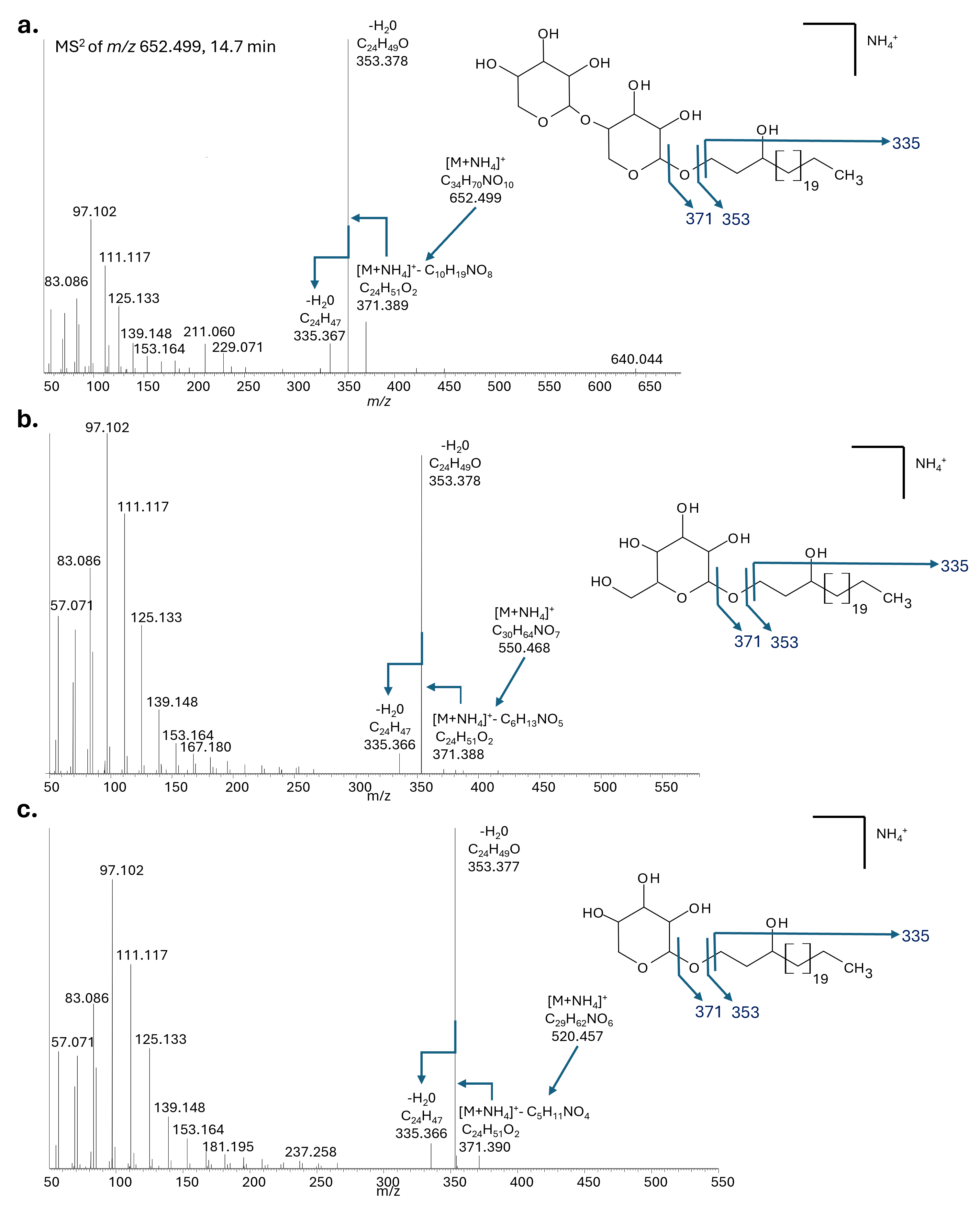

**Supplementary Fig. 22. UHPLC-HRMS MS^2^ spectra of larger components giving rise to fragment ions at *m/z* 371.389.** **a,** compound eluting at 14.67 min ([M+NH_4_]^+^ at *m/z* 652.499) tentatively assigned as 1-(O-dipentopyranose)-3-tetracosanol. **b,** compound eluting at 14.81 min ([M+NH_4_]^+^ at *m/z* 550.468) tentatively assigned as a 1-(O-hexose)-3-tetracosaneol. **c,** compound eluting at 15.22 min giving rise to an [M+NH_4_]^+^ ion at m/z 520.457 tentatively assigned as a 1-(O-pentopyranose)-3-tetracosaneol.

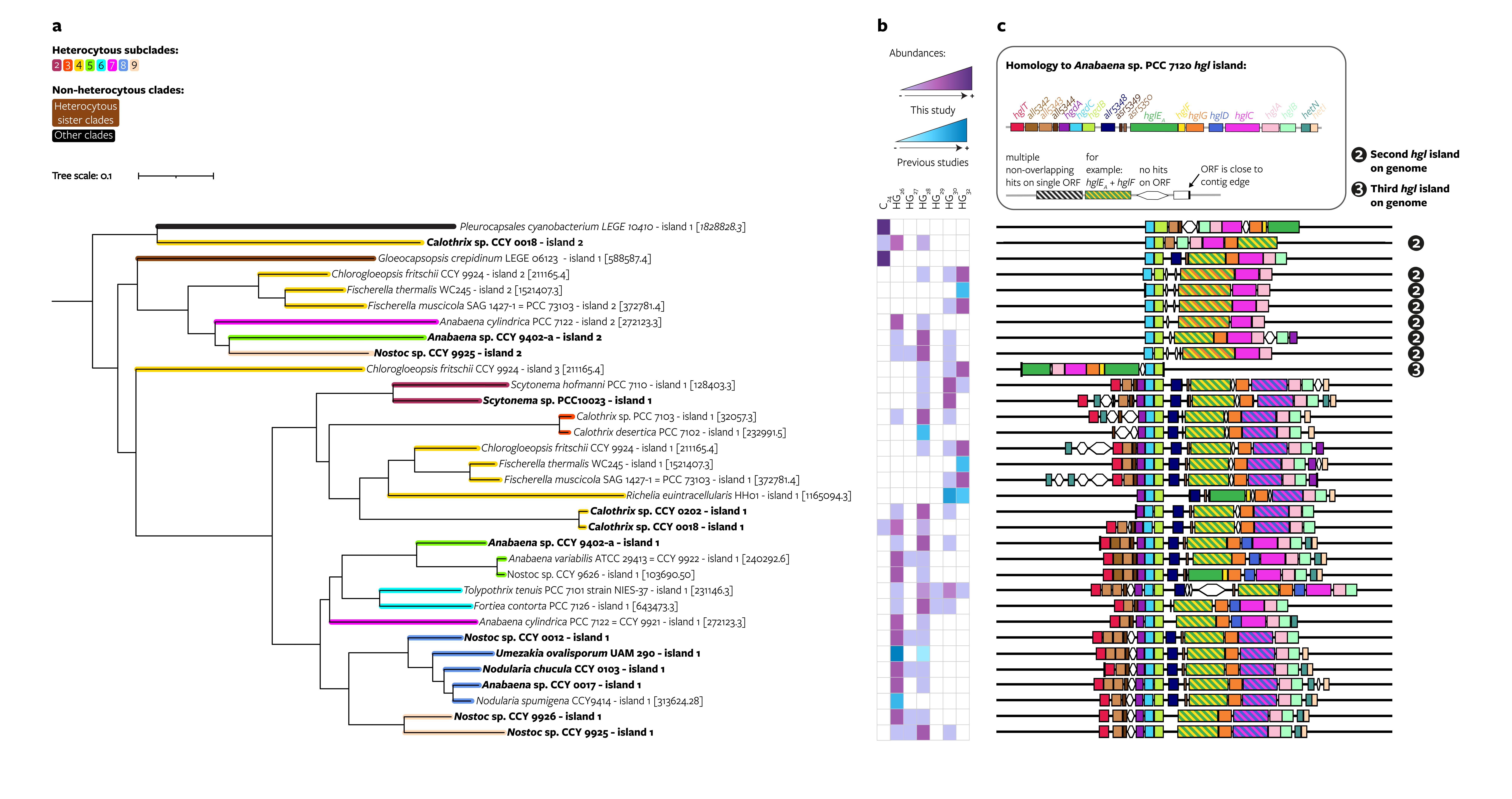

**Supplementary Fig. 23. Distribution of 49 HGs and 4 HG analogs grouped according to alkyl chain length throughout cultures of heterocytous and non-heterocytous Cyanobacteriia. a**, Maximum likelihood phylogeny of the hgl island created using a concatenated alignment of homologous sequences of 7 HG biosynthesis genes (hgdCB and hglE_A_FGCA) that are often present on hgl islands including only genomes with a known HG lipid profile (pruned, see Online Methods). Genomes sequenced in this study are shown in bold. Some genomes contain multiple hgl islands and are in the tree multiple times. **b**, Heatmap of HG and HG analog abundances grouped according to their alkyl chain length. HG analogs (marked as C_24_ in the panel) are the sum of tetracosane-1,3-diol, 1-(O-hexose)-3-tetracosaneol, 1-(O-pentopyranose)-3-tetracosaneol, and 1-(O-dipentopyranose)-3-tetracosaneol. Relative abundances of HGs in Calothrix sp. CCY 0018 are scaled according to the total sum of HGs + HG analogs, which differs from other figures in which HG abundances in Calothrix sp. CCY 0018 are shown. Relative abundances obtained from this study are shown in purple and HG abundances obtained from literature are shown in blue. **c**, Schematic representation of the genes present on the hgl island. When more than one island is present in the genome of the strain, the same HG abundances heatmap is shown for each island. ‘First’, ‘second’, and ‘third’ hgl islands are based on the presence of other islands on the genome, where the ‘first’ hgl island (not marked) is the most extended island in terms of number of Anabaena sp. PCC 7120 genes with homologs on the island, the second hgl island (marked with a 2) the second-most extended island, and the third island (marked with 3) the third-most extended island. ORF, open reading frame; contig, contiguous sequence.
